# A multispecies amplicon sequencing approach for genetic diversity assessment in grassland plant species

**DOI:** 10.1101/2021.07.26.453819

**Authors:** Miguel Loera-Sánchez, Bruno Studer, Roland Kölliker

## Abstract

Grasslands are widespread and economically relevant ecosystems at the basis of sustainable roughage production. Plant genetic diversity (PGD; i.e., within-species diversity) is related to many beneficial effects to the ecosystem functioning of grasslands. The monitoring of PGD in temperate grasslands is complicated by the multiplicity of species present and by a shortage of methods for large-scale assessment. However, the continuous advancement of high-throughput DNA sequencing approaches have improved the prospects of broad, multispecies PGD monitoring. Among them, amplicon sequencing stands out as a robust and cost-effective method.

Here we report a set of twelve multispecies primer pairs that can be used for high-throughput PGD assessment in multiple grassland plant species. The loci targeted by the amplicons were selected and tested in two phases: a “discovery phase” based on a sequence capture assay (611 target nuclear loci assessed in 16 grassland plant species), which resulted in the selection of eleven loci; and a “validation phase”, in which the selected loci were targeted and sequenced using twelve multispecies primers in test populations of *Dactylis glomerata* L., *Lolium perenne* L., *Festuca pratensis* Huds., *Trifolium pratense* L. and *T. repens* L. The resulting multispecies amplicons had overall nucleotide diversities per species ranging from 5.19 × 10^−3^ to 1.29 × 10^−2^, which is in the range of flowering-related genes but slightly lower than pathogen resistance genes. We conclude that the methodology, the DNA sequence resources, and the amplicon-specific primer pairs reported in this study provide the basis for large-scale, multispecies PGD monitoring in grassland plants.

## Introduction

Grasslands cover an estimated 40% of Earth’s land, fulfilling many ecosystem services including the preservation of soil integrity and regulation of water, carbon and nitrogren flows (Bengtsson et al., 2019; Reynolds, 2005; Zhao, Liu, & Wu, 2020). Multispecies grasslands are important sources of roughage for ruminant livestock and provide the basis for sustainable meat and dairy production (Zhao et al., 2020). They are also of unique cultural relevance and serve as places of recreation (Huber & Finger, 2020; Zhao et al., 2020). Plant biodiversity, including genetic diversity (i.e., within-species diversity), plays an important role in grassland ecology. Recent work has shown that high levels of plant genetic diversity in grasslands confer population resistance against invasive plants (Hadincová et al., 2020) and increase yield stability under environmental stress conditions, in part due to interactions between plant species richness and genetic diversity (Malyshev et al., 2016; Meilhac et al., 2019; Prieto et al., 2015). Furthermore, valuable genetic resources for forage breeding are to be found along the wide geographical range for grasses (Poaceae) and legumes (Fabaceae), the two most economically relevant families of forage crop species found in grasslands. Nevertheless, compared to species richness, genetic diversity is still underexplored for most taxonomic groups in all domains of life, including many grass and legume species found in semi-natural grasslands of temperate regions. This is mainly because most traditional assessment methods are time- and resource-demanding, particularly for constant and broad monitoring, which is the aim of current international biodiversity protection initiatives, such as the Convention of Biological Diversity of the United Nations (Hoban et al., 2020; Laikre et al., 2020; Pärli et al., 2021).

Recent advances in high-throughput sequencing technologies are rapidly improving this scenario. Such technologies have enabled genetic diversity assessments in non-model organisms, including the forage crop species from temperate regions (Loera-Sánchez, Studer, & Kölliker, 2019).

Compared to traditional methods for genetic diversity assessment (e.g., with SSRs), hundreds of samples and tens of thousands of markers may be processed simultaneously with high-throughput sequencing approaches. High-throughput approaches suitable for genetic diversity assessment in non-model organism include complexity reduction methods (e.g., genotyping-by-sequencing [GBS] and restriction site-associated DNA sequencing [RADseq]) and target enrichment methods (e.g., sequence capture and amplicon sequencing. Those methods differ in the amount of DNA they require, the marker density (i.e., the number of SNPs) they produce and the costs they imply (Carroll et al., 2018; Harvey et al., 2016).

The choice of a method to assess genetic diversity in grassland plant species in a large scale would require making a balance of its technical features along with the biological features of grassland plants. Grassland plant species often display high levels of intrapopulation genetic diversity and low levels of interpopulation differentiation, which is characteristic of outbreeding species (Hamrick & Godt, 1996). This is the case for many important forage crops, including ryegrasses (*Lolium* spp.), fescues (*Festuca* spp.), orchardgrass (*Dactylis glomerata* L.) and clovers (*Trifolium* spp.; (Collins et al., 2012; Cuyeu et al., 2013; Last et al., 2013; Liu et al., 2018). Because of such high levels of genetic diversity, large samples per population are usually necessary to estimate genetic diversity in these species (Kölliker et al., 2009). However, likely because of such high levels of diversity, a high marker density is not necessary to detect genetic differentiation in populations of such taxa. This has been observed in *Lolium perenne* L. (Liu et al., 2018), as well as in other highly diverse species, such as open-pollinated maize (*Zea mays* L.) landraces (Caldu-Primo et al., 2017) and *Arabidopsis halleri*, (L.) O’Kane & Al-Shehbaz, an outbreeding model species (Fischer et al., 2017).

In this study, we present a novel genetic diversity assessment approach based on amplicon sequencing that can be used in multiple forage grass and legume species. Taking advantage of the naturally high levels of genetic diversity and low levels of linkage disequilibrium of such outcrossing species, we hypothesized that a reduced set of nuclear loci would contain enough polymorphisms for genetic diversity analysis. We describe the steps we took to find such loci in sixteen forage crop species from sub-alpine grasslands (the “discovery” phase of this study) and how we evaluated them in test populations of five of such species, including pooled-plant samples (the “validation” phase). We then discuss the prospects and improvement avenues for genetic diversity assessment in grassland plants based on multispecies amplicon sequencing.

## Materials and Methods

The development of multispecies amplicons for genetic diversity assessments in grassland plant species followed a two-phase approach. In the “discovery phase”, the genetic diversity of 611 conserved loci was assessed in 16 forage species with targeted sequencing. Then, in the “validation phase”, selected multispecies amplicons were sequenced in test populations of five species (Figure 1).

**Figure 1.**
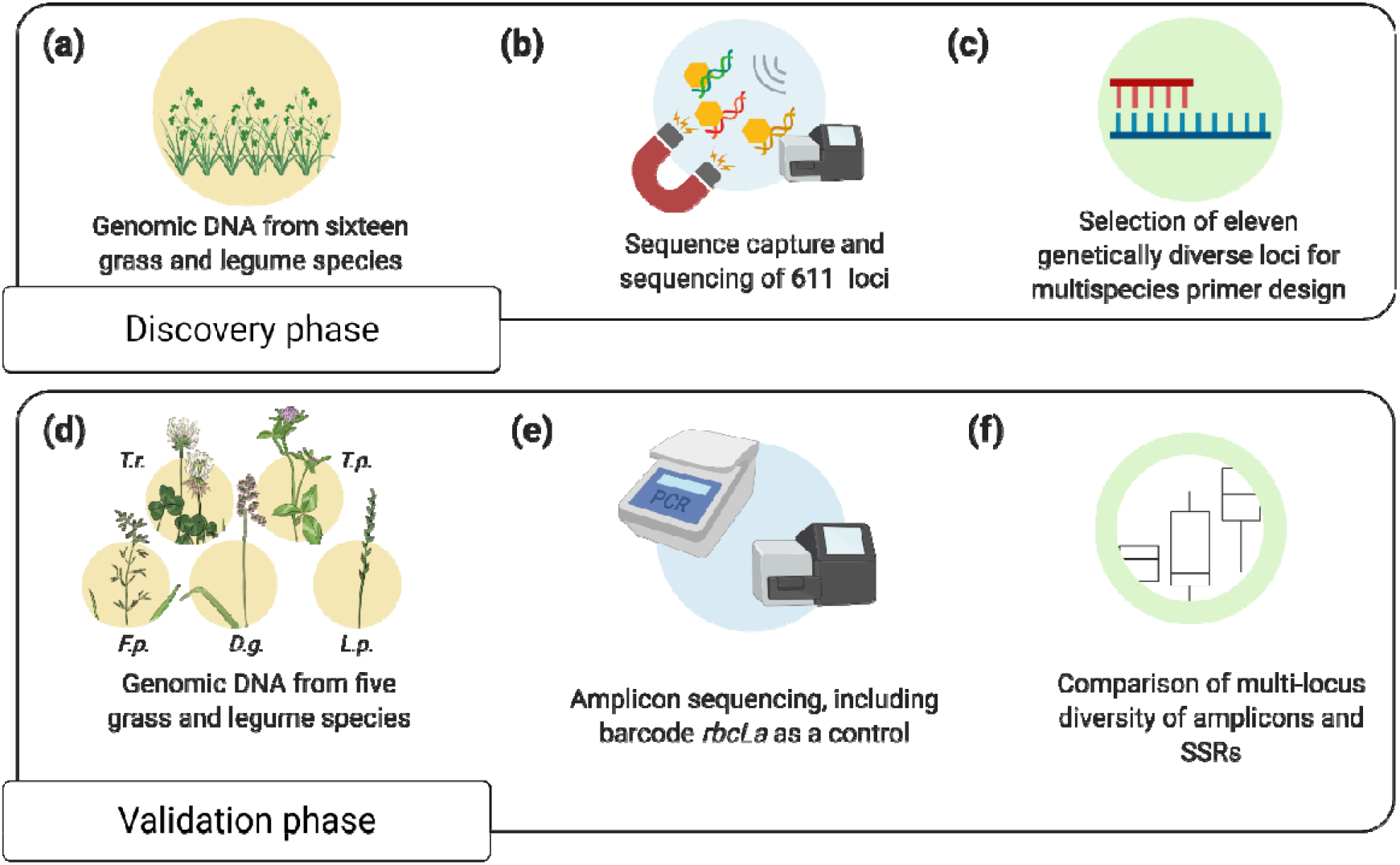
Graphical summary of the steps leading to the identification of multispecies amplicons that were used for genetic diversity assessments. **(a)** Genomic DNA was extracted from 16 grass and legume species (five plants from different cultivars per species). Each species was represented by five single-plant samples and one pooled-plant sample. The pooled plant sample of each species contained the same five plants used for single-plant samples. **(b)** Illumina dual-indexed libraries were prepared (~550 bp fragment size) and sequenced. Libraries were then used as targets for sequence capture. The target loci were 611 single-copy, orthologous genes and ultra-conserved like elements. **(c)** Eleven loci were selected for multispecies primer design, aiming for amplicons of ~500 bp. **(d)** Genomic DNA was extracted from sixteen plants from five grass and legume species (16 plants from three cultivars per species). Each species was represented by sixteen single-plant DNA samples and three pooled-plant DNA samples. In each species, pooled-plant samples included the same plants used for the single-plant samples. **(e)** Multispecies amplicons were amplified all samples, including the DNA barcode *rbcLa* as a reference for a low-diversity locus. Amplicons were pooled equimolarly and dual-indexed libraries and sequenced on the Illumina MiSeq platform. Diversity estimations took place as described above. **(f)** The sixteen single-plant DNA samples were genotyped using eigth SSRs. The resulting SSR multi-locus genotypes were compared to amplicon sequencing-based multi-locus genotypes, in terms of their multi-locus genotype count and pairwise distances. Figure created with https://BioRender.com. Art by Danira León https://www.behance.net/leondanira1c28.

### Discovery phase

#### Bait design and synthesis

A set of 734 putatively single-copy, orthologous genes (SCOGs) was identified with OrthoMCL (L. Li, Stoeckert Jr., & Roos, 2003) in selected reference genomes including *Arabidopsis thaliana* (L.), Heynh. (GCA_000001735.1; Lamesch et al., 2012), *Brachypodium distachyon* (L.) P.Beauv. (GCA_000005505.4; Vogel et al., 2010), *Glycine max* (L.) Merr., 1917(GCA_000004515.4; Schmutz et al., 2010), *Solanum lycopersicum* L. (GCA_000188115.3; Sato et al., 2012), *Theobroma cacao* (L.) (GCA_000403535.1; Motamayor et al., 2013), *Trifolium pratense* L. (GCA_900079335.1; De Vega et al., 2015)and *Vitis vinifera* L. (GCA_000003745.2; Jaillon et al., 2007). The reference genomes were downloaded from EnsemblPlants (https://plants.ensembl.org/index.html). An additional genome assembly of *Lolium perenne* L. (GCA_001735685.1) was obtained from NCBI (https://www.ncbi.nlm.nih.gov). Bait design was performed using BaitFisher v1.2.7 (Mayer et al., 2016) and non-continuous multiple sequence alignments of the 734 SCOGs as input. The annotation of *A. thaliana* (GCA_000001735.1) was used to account for intron-exon boundaries.

Additionally, 1,277 Ultra Conserved-Like Elements (ULEs) were identified with Phyluce v1.4 (Faircloth, 2016). For this purpose, the same set of genomes mentioned above was complemented with *Aegilops tauschii* Coss. (GCA_002575655.1; Luo et al., 2017), *Leersia perrieri* (A.Camus) Launer (GCA_000325765.3), *Lotus japonicus* (Regel) K.Larsen, 1955 (GCA_000181115.2; Sato et al., 2008), *Medicago truncatula* Gaertn. (GCA_000219495.2; Tang et al., 2014), *Oryza sativa* L. subsp. *japonica* (GCA_001433935.1; Kawahara et al., 2013) and *Phaseolus vulgaris* L., 1753 (GCA_000499845.1; Schmutz et al., 2014). The Phyluce UCE-identification pipeline was performed thrice using one of the following guiding genomes in each iteration: *A. thaliana* (GCA_000001735.1), *B. distachyon* (GCA_000005505.4) and *M. truncatula* (GCA_000219495.2). Phyluce v1.4 was then used to find candidate bait sequences.

To control which genomic regions were targeted, the bait sequences were mapped to three annotated reference genomes: *A. thaliana* (GCA_000001735.1), *B. distachyon* (GCA_000005505.4) and *M. truncatula* (GCA_000219495.2). Bait sequences were used to synthesize a custom myBaits® kit (Arbor Biosciences, MI, USA) from now on referred to as the “FORAGE-611” baits.

#### Plant DNA extraction

Seeds of 16 forage species were germinated on filter paper and their seedlings were transferred into pot trays (77 wells, 50 cm × 32 cm, with compost as substrate). The 16 species were: *Alopecurus pratensis* L., *Arrhenaterum elatius* L., *Cynosurus cristatus* L., *Dactylis glomerata* L., *Festuca pratensis* Huds., F. *rubra* L., *L. perenne, Lolium multiflorum* Lam., *Lotus corniculatus* L., *Medicago sativa* L., *Onobrychis viciifolia* Scop., *Phleum pratense* L., *Poa pratensis* L., *T. pratense, Trifolium repens* L. and *Trisetum flavescens* L. After 3-5 weeks, five single plants from different cultivars per species were sampled (Supplementary Table 1). For grasses, samples consisted of three leaf fragments of ~1 cm; for legumes, samples consisted of three young leaflets were harvested per plant. The plant material was freeze-dried for 48 h and pulverized in a QIAGEN TissueLyser II (QIAGEN, Hilden, Germany). DNA was extracted using the NucleoSpin® II kit (Macherey-Nagel, Düren, Germany) and its integrity visually inspected by agarose gel electrophoresis (1% w/v). DNA purity and concentration were determined with a NanoDrop™ spectrophotometer (ThermoFisher Scientific, Waltham, MA, USA).

#### Sequence capture and DNA sequencing

Dual-indexed libraries were constructed using the NEBNext® Ultra™ II DNA Library Prep Kit for Illumina (New England Biolabs, UK). Each species was represented by six libraries: five single-plant libraries and one pooled DNA library, for a total of 96 libraries. The pooled-plant libraries consisted of equimolarly pooled DNA from the 5 single-plants. The average library insert size was ~550 bp.

After indexing, the libraries were divided in four pools of 6× libraries, four pools of 8× libraries and four pools of 10× libraries. Each pool was hybridized to the FORAGE-611 baits. The target DNA fragments were enriched following manufacturer instructions. Fragments were sequenced using the Illumina MiSeq v3 Reagent Kit (2 x 300 bp, 600 cycles). The raw pair-ended reads were merged using BBMerge (Brian Bushnell, Rood, & Singer, 2017) with default parameters. Adapter removal and quality filtering was done using fastp (Chen et al., 2018) with default parameters on the merged reads.

#### Sequencing quality control

The pooled DNA libraries were used to construct pseudo-reference assemblies (pseudo-RAs) using SPAdes (Bankevich et al., 2012). One pseudo-RA was assembled for each species. The sequences of the FORAGE-611 baits were mapped to the pseudo-RAs using BBMap (B. Bushnell, 2014) with default parameters. Bait-mapping coordinates were determined with SAMtools (H. Li, 2011; H. Li et al., 2009) and BEDtools (Quinlan & Hall, 2010). Target locus centers were defined as the middle-point between the 3’-most and 5’-most coordinate of the mapped bait sequences that belong to the same locus. A target locus region was defined as the sequences spanning 500 bp up- and downstream from each locus center. In the cases where baits mapped to more than one pseudo-RA contig, only the largest contig was kept for further analysis.

The 80 single-plant libraries were mapped to their corresponding pseudo-RAs with Bbmap. Reads mapping within target locus regions were considered as “on-target reads”. Duplicate reads were handled with Picard MarkDuplicates (Broad Institute, 2019), marking them for variant calling and removing them for k-mer richness calculations.

A locus with >5 mapped reads was labelled as a quality-controlled locus (QCL). Libraries with >100 QCL and >1,000 total on-target reads were labelled as quality-controlled libraries (QC-libs). Only QC-libs and on-target reads were considered for further analysis. Read-mapping statistics were determined using BEDtools.

#### Diversity metrics

Four diversity metrics were used to rank the 611targeted loci within each species: nucleotide diversity (π), SNP-based nucleotide diversity, SNP density and normalized k-mer richness. Each metric was calculated per locus within each species.

To calculate SNP-based within-species gene diversity, variant calling was done species-by-species using BCFtools mpileup (H. Li, 2011; H. Li et al., 2009) on the BAM files from species with >3 QC-libs. The resulting variant call files (VCFs) were filtered using BCFtools for biallelic SNPs with a minimum quality of 20, a minimum allele frequency of 0.1 and a minimum read depth per sample of 5. Estimations of π were calculated per site using VCFtools, treating all samples as diploid (Danecek et al., 2011). Locus-level estimations were then summed and divided by locus length, which in turn were based on the scaffold lengths in pseudo-RA and had a maximum value of 1 kbp per locus. SNP count and SNP densities were calculated using the same custom R script based on genotype tables produced with VCFtools.

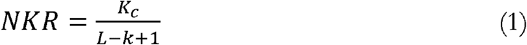

To calculate normalized *k*-mer richness (NKR), de-replicated reads were extracted from de-replicated BAM files using SAMtools bam2fq, creating separate FASTQ files for each locus within each library. *K*-mers of *k* = 25 in each FASTQ file were determined using jellyfish (Marçais & Kingsford, 2011). *K*-mer dumps were filtered by discarding *k*-mers with less than 5x sequencing depth. Locus-specific NKR was calculated at the species level by concatenating the filtered *k*-mer dumps, counting unique *k*-mers within each concatenated dump and then dividing the value of such unique *k*-mer count by maximum expected *k*-mer count per locus (L – *k* + 1). This is summarized in equation 1, where *K*_*c*_ indicates the count of unique *k*-mers, *k* indicates *k*-mer length and L indicates locus length.

Total diversity metrics were calculated for each species. Total NKR was calculated by dividing the sum of all unique *k*-mers in all loci by the total expected *k*-mers in all loci ((L_*total*_ – *k* + 1, where L_*total*_ is the sum of the lengths of all loci in base pairs units). Total SNP count is the sum of all SNPs found in all loci and the SNP density per kbp was calculated following the formula: *SNP*_*total*_ × *1,000*/L_*total*_., where *SNP*_*total*_ is the total SNP count. Total π was calculated using formula 2, where *π*_*i*_ is the nucleotide diversity of the *i*-th locus, L_*i*_ is the length of the *i*-th locus and *a* is the total number of loci.

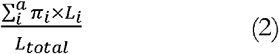

#### Primer design

Eleven loci out of the 611 initially captured were selected for primer design. Selection criteria included having high median gene diversity and normalized *k*-mer richness, as well as being present in as many species as possible. Primers were designed using PriMux (Hysom et al., 2012) with a target amplicon size of 500 bp and primer size of 25 nt. Primers with successful *in silico* and *in vitro* multispecies PCRs were selected for further testing. Thirteen primer pairs were selected for synthesis (Microsynth AG, Balgach, Switzerland). Six primer pairs were grass-specific, six were legume-specific and one pair was found in both plant families. Of those, a final set of twelve primer pairs produced amplicons that were in distinct within each family.

### Validation phase

#### Plant DNA extraction

For single-plant DNA extractions, seeds of five forage species (*D. glomerata, F. pratensis, L. perenne, T. pratense* L. and *T. repens*) were germinated and used for DNA extraction after 3 to 4 weeks of growth as described above. Each species was represented by 16 individual plants from three different cultivars (Supplementary Table 4). For pooled-plant DNA extractions, ground leaf material from the same 16 plants per species was pooled at equal proportions (50 mg per plant), mixed and 20 mg of that mixture were used for DNA extraction. DNA was diluted to 5 ng/μL.

#### Amplicon sequencing

Selected amplicons (Table 1) were amplified in 25 μL PCR reactions containing 15 ng of template DNA, 1x flexi buffer (Promega, Madison, WI, USA), 2 mM MgCl_2_, 200 μM dNTPs, each primer at 0.4 μM and 0.75 units of GoTaq® G2 Flexi DNA Polymerase (Promega, Madison, WI, USA).PCR conditions were 5 min at 94°C followed by 25 cycles of 40 s at 94°C, 1 min at a primer-specific melting temperature (Tm; Table 1) and 40 s at 72°C, followed by a final extension cycle of 10 min at 72°C. The amplicons for each sample were then equimolarly pooled.

**Table 1.**
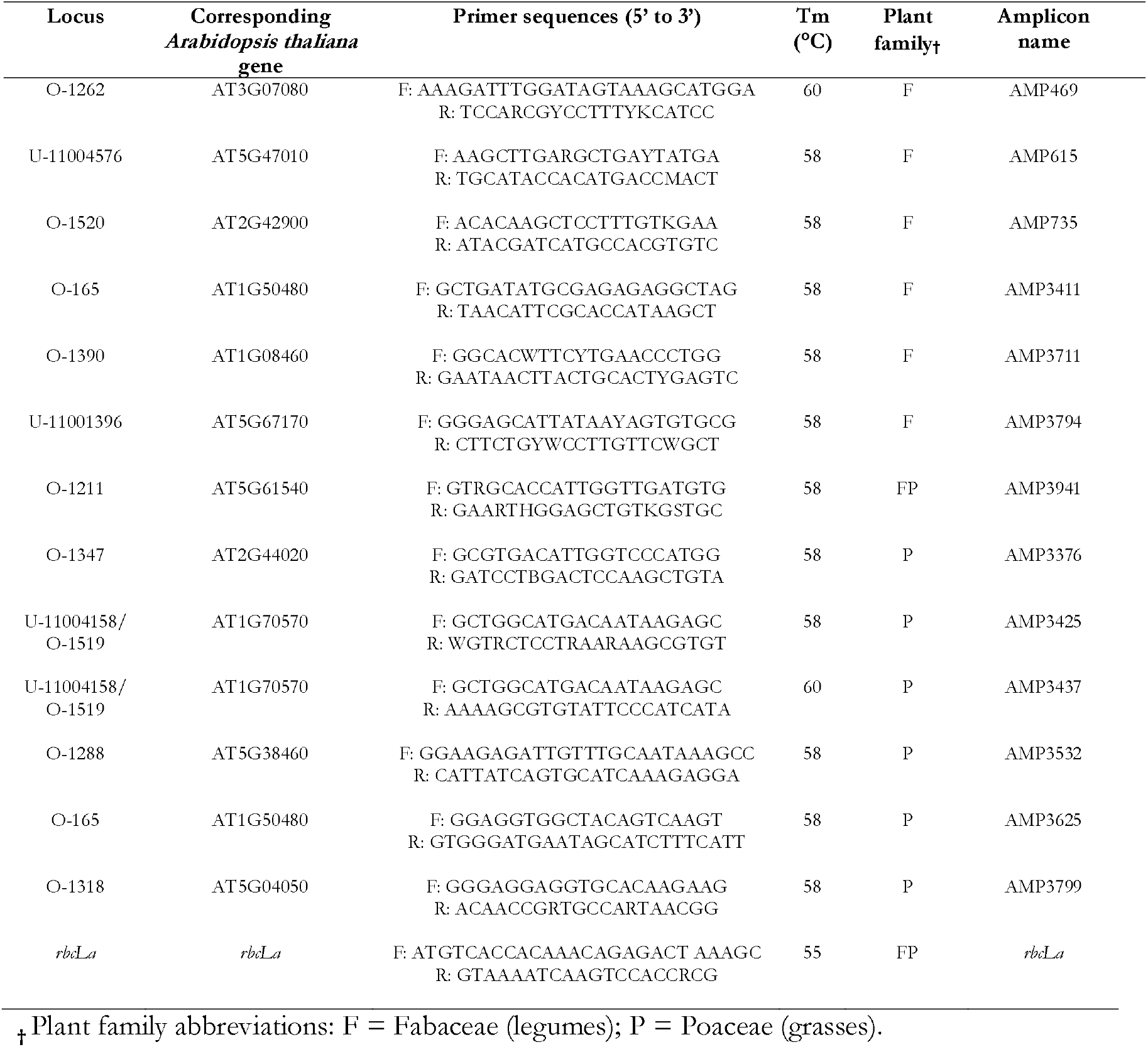
Loci selected for amplicon sequencing and their corresponding multispecies primer pairs sequences.

Dual-indexed libraries were constructed using the NEBNext® Ultra™ II DNA Library Prep Kit for Illumina (New England Biolabs, UK). In total, 80 single-plant libraries (16 per species) and 15 pooled-plant libraries (three per species) were prepared. Dual-indexed libraries were equimolarly pooled and sequenced using the Illumina MiSeq v3 Reagent Kit (2 x 300 bp reads; 600 cycles). Merging of raw pair-ended reads, adapter removal and quality filtering were done as described above. Additionally, to remove possible remnants of primer sequences, reads were trimmed 25 bp at each end using cutadapt (Martin, 2011).

#### Diversity metrics

To calculate SNP-based diversity metrics, quality-controlled reads were mapped to pseudo-reference assemblies, which consisted only on the sequences of the selected loci taken from the sequence capture data of the discovery phase. The pseudo-reference assemblies also contained the sequences of DNA barcode *rbcLa* corresponding to each species, which were obtained from the Barcoding Of Life Datasystems (Ratnasingham & Hebert, 2007), project SWFRG (Loera-Sánchez, Studer, & Kölliker, 2020). For single-plant libraries, variant calling and SNP-based diversity metrics were calculated as described above. Variant calling was performed simultaneously on the BAM files of the sixteen single-plant libraries of each species. In the case of pooled-plant libraries, amplicon-wise π and SNP counts were calculated using the “Variance-at-position.pl” script from PoPoolation v1.2.2 (Kofler et al., 2011) with a minimum covered fraction of 0.5,. a pool size of 32 (i.e., the equivalent of 16 diploid plants), a minimum site quality of 20, a minimum coverage of 10, and a minimum allele count of 5. The PoPoolation analysis was run individually for each BAM file of the pooled-plant libraries.

SNP haplotypes were reconstructed by concatenating each SNP position from each sequencing read for both single- and pooled-plant samples. For this, BAM files containing the read mappings to the amplicon pseudo-reference assemblies were parsed to extract reads and SNP positions using a custom Python script. The SNP positions came from the mappings of the single-plant libraries. Determining the final haplotype-based genotype required filtering out spurious haplotypes with low read counts, which likely are due to PCR or sequencing errors. Briefly, for each amplicon, the haplotype with the maximum read count was determined. Such maximum read count was used to normalize read counts for the rest of the haplotypes. Normalized haplotype allele counts thus ranged from zero to one. Only haplotypes with normalized counts higher than a specific cutoff value (0.9 for diploids, 0.3 for tetraploids and 0.2 for pools) were retained. For single-species samples, haplotypes were further visually inspected, and the following was corrected: a) haplotypes with missing values were removed, b) haplotypes with tri- or tetra-allelic SNPs were removed and c) co-occurring haplotypes with only one different SNP allele call were consolidated into one haplotype. No visual inspection was conducted for pooled-plant samples, but only haplotypes that were present in single-species samples were retained.

Haplotype-based multi-locus genotypes, pairwise Prevosti’s distances, allele count and expected heterozigosities were calculated using the ‘poppr’ R package (Kamvar, Tabima, & Gr◻unwald, 2014).

To calculate normalized *k*-mer richness, reads were extracted from de-replicated BAM files using SAMtools bam2fq, creating separate FASTQ files for each amplicon within each library. Each FASTQ file was then sub-sampled to contain 100 reads using Seqtk v1.3 (*Seqtk-1.3*, 2018). *K*-mers of *k* = 25 in each sub-sampled FASTQ file were determined with jellyfish (Marçais & Kingsford, 2011). The resulting *k*-mer dumps were filtered by discarding *k*-mers with <25x sequencing depth. Amplicon-specific NKR was calculated at the species level as described above.

Total diversity per species was calculated using exclusively single-plant libraries. Calculations were conducted as described in the discovery phase. The count of SNP-based haplotypes (H.Ac) for all amplicons was also calculated for each species. Genetic diversity metrics for each sample were compared to the diversity of the DNA barcode *rbcLa*, which is known to have a very low within-species diversity.

#### SSR and multi-locus analysis

Single-plant DNA samples were also used as templates for SSR analyses. In total, eight species-specific SSR primer pairs were used. SSR primers and PCR conditions are described in Supplementary Table 5. Genotype tables were analyzed using the R package ‘poppr’ (Kamvar et al., 2014). Summary statistics per locus are shown in Supplementary Table 6.

The five amplicons with the highest coverage per species were selected for multi-locus analysis (AMP469 was excluded for *T. repens* as it is potentially duplicated in this species; Table 8, column 3). Only amplicons from single-plant libraries were analyzed. Accordingly, the five most diverse SSRs per species were selected for multi-locus analysis (again, four for *T. repens*; Table 8, column 4).

Multi-locus genotypes were produced for SSRs (from now on “SSR-MLGs”) and for amplicon-based haplotypes (from now on “HAP-MLGs”) using the R package ‘poppr’ (Kamvar et al., 2014). Pairwise Prevosti’s distance was calculated for SSR- and HAP-MLGs using the R package ‘poppr’ (Kamvar et al., 2014). Plants with >5% missing loci, either amplicons or SSRs, were removed from this analysis.

In addition, multi-locus *k*-mer distances were calculated using sourmash (Titus Brown & Irber, 2016). For this, the normalized FASTQ files described above were merged into multi-amplicon FASTQ files according to library name. Only FASTQ files corresponding to single-plant libraries and to the amplicons selected for amplicon-MLGs were merged. Multi-amplicon FASTQ files were used to calculate *k*-mer signatures using sourmash sketch (Titus Brown & Irber, 2016). Finally, pairwise similarities among samples of the same species were calculated using sourmash compare (Titus Brown & Irber, 2016). Distances were calculated from similarities by applying the formula: *d* = 1 – *s*, were *d* is distance and *s* is simalirity. The ratios among the three kinds of distances were calculated for each pair of plants and summarized in Table 8.

## Results

### Discovery phase

#### Target loci selection and bait design

A total of 611 loci (265 ULEs and 346 orthologous genes) were selected, for which 12,253, 100-nucleotide long baits were designed (5,526 targeting ULEs and 6,727 orthologous genes). In total, 602 loci mapped to a gene model in any of the three reference genomes used as controls (*A. thaliana, B. distachyon and M. truncatula*). Of those loci, 365 were present in all three reference genomes, 142 were present in only in *A. thaliana* and *B. distachyon*, 31 in *A. thaliana* and *M. truncatula*, 31 in *B. distachyon* and *M. truncatula*, ten only in *A. thaliana*, ten in *B. distachyon* and thirteen in *M. truncatula* (Supplementary Table 2 and Supplementary Figure 1).

In any of the three reference genomes, a total of 85 genes contained two or more target loci and, conversely, 23 loci mapped to two or more genes (Supplementary Table 3).

#### Sequence capture and sequencing

Around 14.5 million raw read pairs were obtained after sequence capture, 10.3 million reads for grasses and 4.2 million for legumes (Table 2). At the species level, total raw output ranged from 512,233 read pairs for *T. flavescens* to ~1.5 million read pairs for *L. corniculatus*. The N50 of pseudo-RAs generated from the pooled-plant samples ranged from 590 bp (*T. pratense*) to 4,905 bp (*A. pratensis*).

Approximately two million read pairs were successfully quality-controlled, merged and mapped to a target locus. This constitutes a total capture efficiency of 13.45% of the total raw reads. Within grasses and legumes, capture efficiencies were 4.78% (495,858 quality-controlled, merged reads) and 35% (~1.5 million quality-controlled, merged reads), respectively. At the species level, capture efficiency varied from 1.58% for *A. elatius* (18,735 quality-controlled, merged reads) to 55.82% for *T. pratense* (537,289 quality-controlled, merged reads). Read duplication in quality-controlled, merged reads ranged from 12.75% in *F. rubra* to 38.59% in *T. pratense*.

For further analysis, only quality-controlled loci (QCL: loci with five or more quality-controlled, merged reads within a library) and quality-controlled libraries (QC-libs: libraries with >100 QCL and >1,000 quality-controlled, merged reads) were considered. Read counts were always calculated as quality-controlled, merged reads. In total, 60 QC-libs with a median of 438 QCL per library (interquartile range, IQR = 355-532; Table 2) were obtained. Twenty-three loci were not captured in any library. Each QCL had a median of *37* reads (IQR = 17 - 84). For grasses, there were 37 QC-libs with a median of 377 QCL per library (IQR = 313 - 410); median reads per grass QCL was 21, (IQR = 12 - 36). For legumes, there were 23 QC-libs with a median of 548 QCL (IQR = 520 - 552); median reads per legume QCL was 84 (IQR = 48 - 136). At the species level, QC-lib varied from one for *T. flavecens* to a maximum of five obtained for *C. cristatus, D. glomerata, L. corniculatus, O. viciifolia and T. pratense*. Median read count per QCL varied from 13 in *L. perenne* (IQR = 9 - 20) to 149 in *T. pratense* (IQR = 97 - 227). Median QCL count varied from 213 for *A. elatius* (IQR = 190 - 233) to 579 for *T. repens* (IQR = 574 - 584).

#### Genetic diversity estimations

Diversity statistics were calculated for each locus within each species. Overall, normalized k-mer richness (NKR) values ranged from zero (locus ortho-1479 in *T. flavescens*) to 13.04 (locus ortho-147 in *T. repens*) with a median of 1.06 (IQR = 0.82 - 1.35). SNP count ranged from one (326 locus-species pairs) to 86 (loci ortho-1288/uce-11005138 in *Phleum pratense*) with a median of 9 (IQR = 4 - 15). SNP densities ranged from 1 (36 locus-species pairs) to 99.83 (ortho-1178 in *T. pratense*) with a median of 10.79 (IQR = 5 - 19.27). Nucleotide diversity (π) ranged from 2.5×10^−4^ (ortho-1106 and uce-12001183 in *M. sativa*) to 5.17×10^−2^ (ortho-1186/uce-11003333 in *F. pratensis*) with a median of 4.99×10^−3^ (IQR = 2.33×10^−3^ - 9.08×10^−3^).

In general, NKR and π showed a very low correlation (*r*^2^ = 0.08, Fig. 1a). At the species-level, *r*^2^ values ranged from 0.1 for *P. pratensis* to 0.47 in *L. multiflorum*, while for most species *r*^2^ ≥ 0.20 (Fig. 1b).

Locus selection was based on such median diversity values across species, as well as in species count (i.e., the number of species were a locus was found) and its suitability for multispecies primer design. Across species, median NKR values ranged from 0.45 (ortho-350) to 8.78 (ortho-106). Median SNP count ranged from 1 SNP (ortho-1474 and ortho-1498) to 43 SNPs (uce-11005074). Median SNP densities ranged from 1.01 (ortho-1474) to 49.7 (ortho-1198). Median π ranged from 5.3×10^−4^ (ortho-1498) to 2.1×10^−2^ (uce-11005074). Median diversity metrics across species for the selected loci are shown in Table 4. A Wilcoxon test showed significant differences (P-value < 0.001) between selected and non-selected loci based on the median of medians of NKR, SNP count, SNP density per kbp and π (Fig. 1c).

**Table 3.** Total diversity statistics per species for captured loci.

| Species name | NKR | SNPs | SNPs/kbp | $\pi$ | $N_t$ |
| --- | --- | --- | --- | --- | --- |
| *Alopecurus pratensis* | 1.18 | 3,819 | 19.66 | $9.50 \times 10^{-3}$ | 234 |
| *Arrhenatherum elatius* | 0.78 | 287 | 5.29 | $2.66 \times 10^{-3}$ | 72 |
| *Cynosurus cristatus* | 1.03 | 1,040 | 7.3 | $3.35 \times 10^{-3}$ | 173 |
| *Dactylis glomerata* | 1.36 | 3,022 | 15.91 | $7.84 \times 10^{-3}$ | 229 |
| *Festuca pratensis* | 1.11 | 3,525 | 26.81 | $1.73 \times 10^{-2}$ | 188 |
| *Festuca rubra* | 0.76 | 3,208 | 14.77 | $7.15 \times 10^{-3}$ | 281 |
| *Lolium multiflorum* | 0.85 | 3,009 | 17.08 | $8.38 \times 10^{-3}$ | 218 |
| *Lolium perenne* | 0.65 | 1,479 | 8.55 | $5.05 \times 10^{-3}$ | 211 |
| *Lotus corniculatus* | 1.29 | 4,339 | 9.68 | $4.25 \times 10^{-3}$ | 477 |
| *Medicago sativa* | 1.20 | 5,615 | 12.47 | $5.38 \times 10^{-3}$ | 507 |
| *Onobrychis viciifolia* | 1.17 | 3,259 | 8.84 | $4.03 \times 10^{-3}$ | 416 |
| *Phleum pratense* | 1.13 | 6,094 | 20.12 | $8.67 \times 10^{-3}$ | 354 |
| *Poa pratensis* | 1.24 | 5,723 | 17.4 | $9.28 \times 10^{-3}$ | 415 |
| *Trifolium pratense* | 1.31 | 2,766 | 6.36 | $2.76 \times 10^{-3}$ | 469 |
| *Trifolium repens* | 1.55 | 8,463 | 20.17 | $9.41 \times 10^{-3}$ | 536 |
| *Trisetum flavescens* | 0.53 | 696 | 4.88 | $4.92 \times 10^{-3}$ | 179 |

† $N_t$ : Total quality-controlled loci captured per species.

**Table 4.** Median genetic diversity statistics of the selected loci.

| Locus | NKR | SNPs | SNPs/kbp | $\pi_{\ddagger}$ | $N$ |
| --- | --- | --- | --- | --- | --- |
| ortho-1390 | 1.00 (0.84 - 1.36) | 9.50 (5.50 - 18.00) | 16.00 (6.66 - 27.32) | $8.09 \times 10^{-3}$ ($2.45 \times 10^{-3}$ - $1.21 \times 10^{-2}$) | 15 |
| ortho-1347 | 1.20 (0.79 - 1.35) | 22.00 (6.00 - 31.00) | 22.00 (6.83 - 34.66) | $8.65 \times 10^{-3}$ ($4.15 \times 10^{-3}$ - $1.09 \times 10^{-2}$) | 15 |
| ortho-1318 | 1.40 (1.06 - 1.60) | 16.50 (8.50 - 35.00) | 26.46 (9.80 - 39.14) | $9.68 \times 10^{-3}$ ($4.29 \times 10^{-3}$ - $1.95 \times 10^{-2}$) | 14 |
| ortho-1262 | 1.42 (1.13 - 1.66) | 16.50 (13.62 - 22.25) | 28.13 (19.12 - 31.58) | $9.94 \times 10^{-3}$ ($7.27 \times 10^{-3}$ - $1.50 \times 10^{-2}$) | 16 |
| ortho-1211 | 1.43 (0.93 - 2.13) | 14.00 (9.50 - 21.00) | 19.71 (10.70 - 29.00) | $6.36 \times 10^{-3}$ ($5.59 \times 10^{-3}$ - $8.63 \times 10^{-3}$) | 16 |
| uce-11004576 | 1.49 (1.12 - 1.82) | 9.00 (2.50 - 14.50) | 9.55 (3.38 - 26.15) | $3.42 \times 10^{-3}$ ($1.33 \times 10^{-3}$ - $8.68 \times 10^{-3}$) | 14 |
| uce-11004158 | 1.68 (1.20 - 2.10) | 24.00 (11.50 - 31.00) | 27.06 (18.69 - 38.19) | $1.03 \times 10^{-2}$ ($8.50 \times 10^{-3}$ - $1.75 \times 10^{-2}$) | 15 |
| ortho-1288 | 1.74 (1.28 - 1.97) | 36.00 (14.25 - 62.75) | 39.82 (15.76 - 68.32) | $2.10 \times 10^{-2}$ ($9.09 \times 10^{-3}$ - $3.49 \times 10^{-2}$) | 12 |
| ortho-165 | 1.75 (1.52 - 2.24) | 15.00 (6.00 - 24.75) | 18.97 (7.89 - 28.09) | $6.23 \times 10^{-3}$ ($2.88 \times 10^{-3}$ - $1.08 \times 10^{-2}$) | 14 |
| ortho-1520 | 1.91 (1.68 - 2.07) | 27.00 (18.00 - 33.00) | 29.22 (19.72 - 39.14) | $1.26 \times 10^{-2}$ ($8.34 \times 10^{-3}$ - $1.73 \times 10^{-2}$) | 3 |
| uce-11001396 | 2.62 (1.33 - 2.77) | 7.67 (6.00 - 13.00) | 14.59 (11.00 - 35.00) | $5.85 \times 10^{-3}$ ($3.64 \times 10^{-3}$ - $8.53 \times 10^{-3}$) | 13 |

$\dagger \pi$ : nucleotide diversity per 1 kbp; $\ddagger N$ : number of species where a locus was found.

### Validation phase

#### Multispecies amplicon sequencing and genetic diversity analysis

Amplicon sequencing in test populations (n = 16) of *D. glomerata, F. pratensis, L. perenne, T. pratense* and *T. repens* produced 12.4 million raw read pairs of which 4.3 million reads (34.81%) were successfully quality-controlled and merged. Raw output was 6.5 million reads for grasses, with 1.8 million quality-controlled, merged reads (26.77% of raw output for this plant family). For legumes, raw output was ~5.9 million reads, with 2.5 million quality controlled, merged reads (43.72%). At the species level, raw output was 1.99, 2.25, 2.30, 3.03 and 2.86 million reads for *D. glomerata, F. pratensis, L. perenne, T. pratense* and *T. repens*, respectively. In turn, quality-controlled, merged read counts were 0.57 (28.79% of raw output for this plant species), 0.64 (28.34%), 0.54 (23.49%), 1.51 (49.67%) and 1.07 (37.42%) million reads for *D. glomerata, F. pratensis, L. perenne, T. pratense* and *T. repens*, respectively. Duplicate rate was always >90% in all groups (total, plant families and species).

At the locus level, quality-controlled amplicons (i.e., amplicons with ≥5 mapped reads) reported a median of >1,000 reads per library. Median count of quality-controlled amplicons was eight in *L. perenne* and *T. pratense* libraries, and seven in the rest of the species. The DNA barcode *psbK-psbI* could not be recovered from legume samples, so it was excluded from analysis. Furthermore, amplicon AMP3794 had zero coverage in *T. repens* libraries. In addition, amplicon AMP3437 had zero coverage in *D. glomerata* and *F. pratense* libraries. The latter case is because AMP3437 targets the same locus as AMP3425 so, from here on, AMP3437 reads were treated as AMP3425 reads.

In single-plant libraries and excluding DNA barcode *rbcLa*, total NKR values ranged from 1 (*F. pratensis*) to 2.05 (*T. repens*). Total SNP counts ranged from 27 (*F. pratensis*) to 62 (*D. glomerata*). Total π ranged from 5.19×10^−3^ to 1.32×10^−2^. Median SNP-based haplotype counts ranged from 19 (*F. pratensis*) to 43 (*L. perenne*).

In single-plant libraries and excluding DNA barcode rbcLa, NKR ranged from zero (AMP3625 in *F. pratensis*) to 2.94 (AMP3799 in *D. glomerata*) with a median of 1.53 (IQR = 1.02-1.97). SNP count ranged from zero (nine species-locus pairs) to 28 (AMP3799 in *D. glomerata*) with a median of 3 (IQR = 0-11). SNP densities ranged from zero (nine species-locus pairs) to 50.36 SNPs/kbp (AMP3799 in *D. glomerata*) with a median of 6.91(IQR = 0-20.14). π ranged from zero (nine species-locus pairs) to 3.83×10^−2^ (AMP3799 in *D. glomerata*) with a median of 4.62×10^−3^ (IQR = 0-1.52×10^−2^). SNP-based haplotype allele count (H.Ac) ranged from one (11 cases) to 25 (AMP3799 in *D. glomerata*) with a median of 3 (IQR = 1 - 7).

In pooled-plant libraries, NKR values ranged from zero (AMP3625 in *F. pratensis* pool 2) to 3.62 (AMP3799 in *D. glomerata* pool 1) with a median of 1.62 (IQR = 1.01 - 2.21). SNP counts ranged from zero (38 species-locus pairs) to 54 (AMP3799 in *D. glomerata* pools 1) with a median of 2 (IQR = 0 - 19). SNP density ranged from zero (38 species-locus pairs) to 101.82 SNPs/kbp (AMP3799 in *D. glomerata* pool 3) with a median of 3.77 (IQR = 0 - 35.83). π ranged from 0 (38 species-locus pairs) to 6.68×10^−2^ (AMP3425 in *L. perenne* pool 1) with a median of 3.13×10^−3^ (IQR = 0 - 1.51×10^−2^). H.Ac ranged from one (41 species-locus pairs) to 22 (AMP3799 in *D. glomerata* pool 1) with a median of 2 (IQR = 1 - 5).

The concordance between diversity metrics calculated for each species-amplicon pair from single-plant libraries and pooled-plant libraries was good. Overall, the coefficients of variation of all diversity metrics for each amplicon and species had a median of 13.38% (IQR = 3.35% - 27.58%). The coefficient of variation (CV) of NKR estimates ranged from 0.11% (AMP3532 in *F. pratensis* and *rbcLa* in *T. pratense*) to 86.67% (AMP3625 in *F. pratensis*) with a median of 8.1% (IQR = 2.43% - 16.68%). The range of the CV of SNP counts ranged from 0% (16 species-amplicon pairs) to 88.79% (AMP3425 in *L. perenne*) with a median of 12.87% (IQR = 0% - 27.75%). In the case of SNP densities, CV ranged from 0% (12 species-amplicon pairs) to 90.60% (AMP3425 in *L. perenne*) with a median of 10.10% (IQR = 0% - 28.65%). The CV of π estimates ranged from 0% (twelve species-amplicon pairs) to 114.47% (AMP3425 in *L. perenne*) with a median of 15.43% (IQR = 0% - 26.06%). Finally, the CV of SNP-based haplotype counts ranged from 0% (22 species-amplicon pairs) to 119.02% (AMP3425 in *L. perenne*) with a median of 0% (IQR = 0% - 23.14%).

At the amplicon level and across species, median CV of NKR estimates ranged from 1.36% (AMP3532) to 56.70% (AMP3625). For SNP counts, median CV ranged from 0% (AMP615, AMP3411, AMP3941 and AMP3794) to 88.79% (AMP3425). For SNP densities, median CV ranged from 0% (*rbcLa* and AMP3941) to 90.60% (AMP3425). For π estimates, median CV ranged from 0% (*rbcLa* and AMP3941) to 114.47% (AMP3425). For SNP-based haplotype counts, median CV ranged from 0% (*rbcLa*, AMP3941, AMP615, AMP3411, AMP3625, AMP3794 and AMP3425) to 28.57% (AMP3376).

Across species, the amplicons were significantly more diverse than the DNA barcode *rbcLa* for at least one diversity metric (Fig. 3a). However, amplicon AMP469 in *T. repens* produced SNP-based haplotypes that do not match the ploidy of this species (i.e., tetraploid). This amplicon was not considered for overall diversity calculations (Table 6) nor for comparisons with SSR-based diversity metrics.

**Table 5.** Amplicon sequencing output summary.

| Family | Species name | Raw read pairs | % Merged<br>on-target<sup>†</sup> | %Duplicates<br>in merged | Reads per amplicon<sup>‡</sup> | Amplicons<br>per library<sup>‡</sup> | Library<br>count |
| --- | --- | --- | --- | --- | --- | --- | --- |
| Fabaceae | *Trifolium pratense* | 3,034,018 | 49.67 | 96.32 | 6,684 (4,119 - 15,068) | 8 | 19 |
|  | *Trifolium repens* | 2,864,276 | 37.42 | 95.99 | 5,947 (2,935 - 11,129) | 7 | 19 |
| Poaceae | *Dactylis glomerata* | 1,987,368 | 28.79 | 91.34 | 2,652 (1,254 - 5,476) | 7 | 19 |
|  | *Festuca pratensis* | 2,250,146 | 28.34 | 92.63 | 2,208 (945 - 6,793) | 7 | 19 |
|  | *Lolium perenne* | 2,306,113 | 23.49 | 91.69 | 1,273 (577 - 4,876) | 8 (7,8) | 19 |
| Fabaceae | All | 5,898,294 | 43.72 | 96.18 | 6,446 (3,336 - 13,948) | 7 (7-8) | 38 |
| Poaceae | All | 6,543,627 | 26.77 | 91.92 | 1,901 (868 - 5,870) | 7 | 57 |
| Total |  | 12,441,921 | 34.81 | 94.46 | 4,149 (1,174 - 8,411) | 7 (7-8) | 95 |

<sup>†</sup> Merged, on-target reads are reads that were quality-controlled and successfully merged and mapped to a reference amplicon; percentage is in relation to raw read pairs. <sup>‡</sup> Median and interquartile range calculated considering only quality-controlled amplicons (e.g., amplicons with five or more mapped reads). Interquartile range shown for groups that have variation.

**Table 6.** Genetic diversity per species based on the multispecies amplicons in single-plant libraries.

| Species name | NKR | SNPs | SNPs/kbp | $\pi$ | H.Ac | Total length (bp) † | N |
| --- | --- | --- | --- | --- | --- | --- | --- |
| *Dactylis glomerata* | 1.89 | 62 | 18.01 | $1.29 \times 10^{-2}$ | 40 | 3,442 | 6 |
| *Festuca pratensis* | 1.00 | 27 | 7.75 | $5.19 \times 10^{-3}$ | 19 | 3,483 | 6 |
| *Lolium perenne* | 1.41 | 44 | 12.61 | $8.49 \times 10^{-3}$ | 43 | 3,490 | 6 |
| *Trifolium pratense* | 1.42 | 36 | 10.28 | $7.63 \times 10^{-3}$ | 36 | 3,503 | 7 |
| *Trifolium repens* | 2.01 | 23 | 11.56 | $1.05 \times 10^{-2}$ | 13 | 1,989 | 5 |
| **Mean** | 1.55 | 38.40 | 12.04 | $8.94 \times 10^{-3}$ | 30.20 | 3,181.40 | 6.00 |

† Sum of the lengths of all amplicons in each species expressed in base pairs (bp). All metrics exclude the DNA barcode *rbcLa* and amplicon AMP469 for *Trifolium repens*.

Sequence-based diversity metrics showed a weak positive correlation to each other (0.25 ≤ *r*^2^ ≤ 0.32, Figure 2b, black). Excluding amplicons without SNP calls marginally improved correlations (0.3 ≤ *r*^2^ ≤ 0.45, Figure 2b, blue). Excluding amplicons without SNP calls and considering only single-plant libraries resulted in considerably higher correlations (0.51 ≤ *r*^2^ ≤ 0.69, Figure 2b, red).

**Figure 2.**
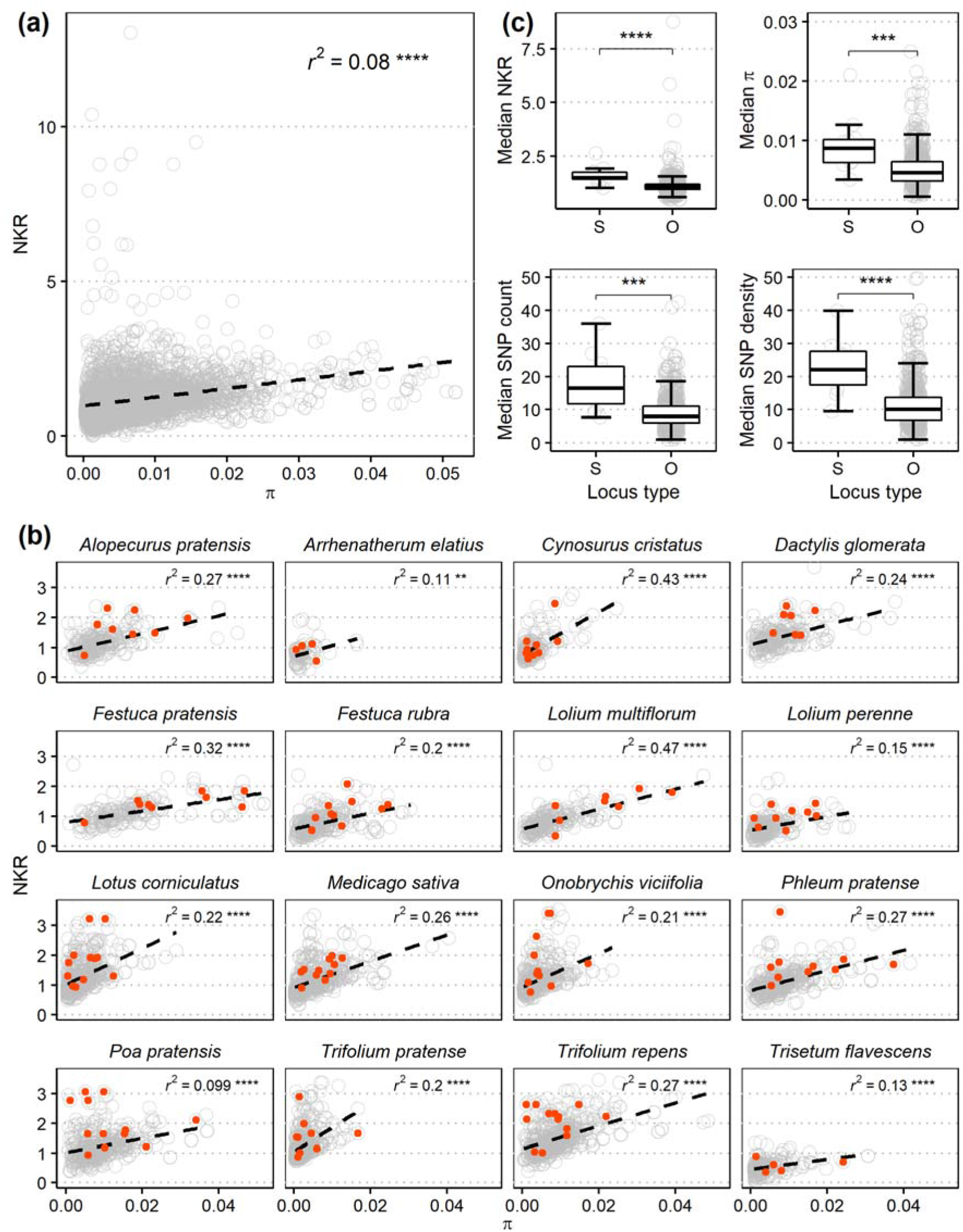
Diversity statistics of the captured loci. **(a)** Nucleotide diversity per kbp (π) vs normalized *k*-mer richness (NKR) for all captured loci in the 60 quality-controlled libraries. Linear regression shown in black dotted line. Regression *r*^2^ value and P-value significance level shown in upper right corner. **(b)** π per kbp vs NKR for all captured loci divided by species. Red dots indicate selected loci. Linear regression shown in black dotted line. Regression *r*^2^ value and P-value significance level shown in bottom right. **(c)** A Wilcoxon test shows that median diversity metrics across the sixteen species are higher in selected loci (“S” label, n = 10) than in other captured loci (“O” label, n = 578). Significance levels for P-values: 0.0001 = ****, 0.001 = ***, 0.01 = **, 0.05 = *, ns = non-significant. The upper and bottom ends of boxplots indicate the 3^rd^ and 1^st^ quartiles, respectively. The black line inside of boxplots indicates the median value. The upper and lower whiskers in boxplots indicate the maximum (3^rd^ quartile + 1.5 times the interquartile range) and minimum (1^st^ quartile - 1.5 times the interquartile range) values, respectively.

**Figure 3.**
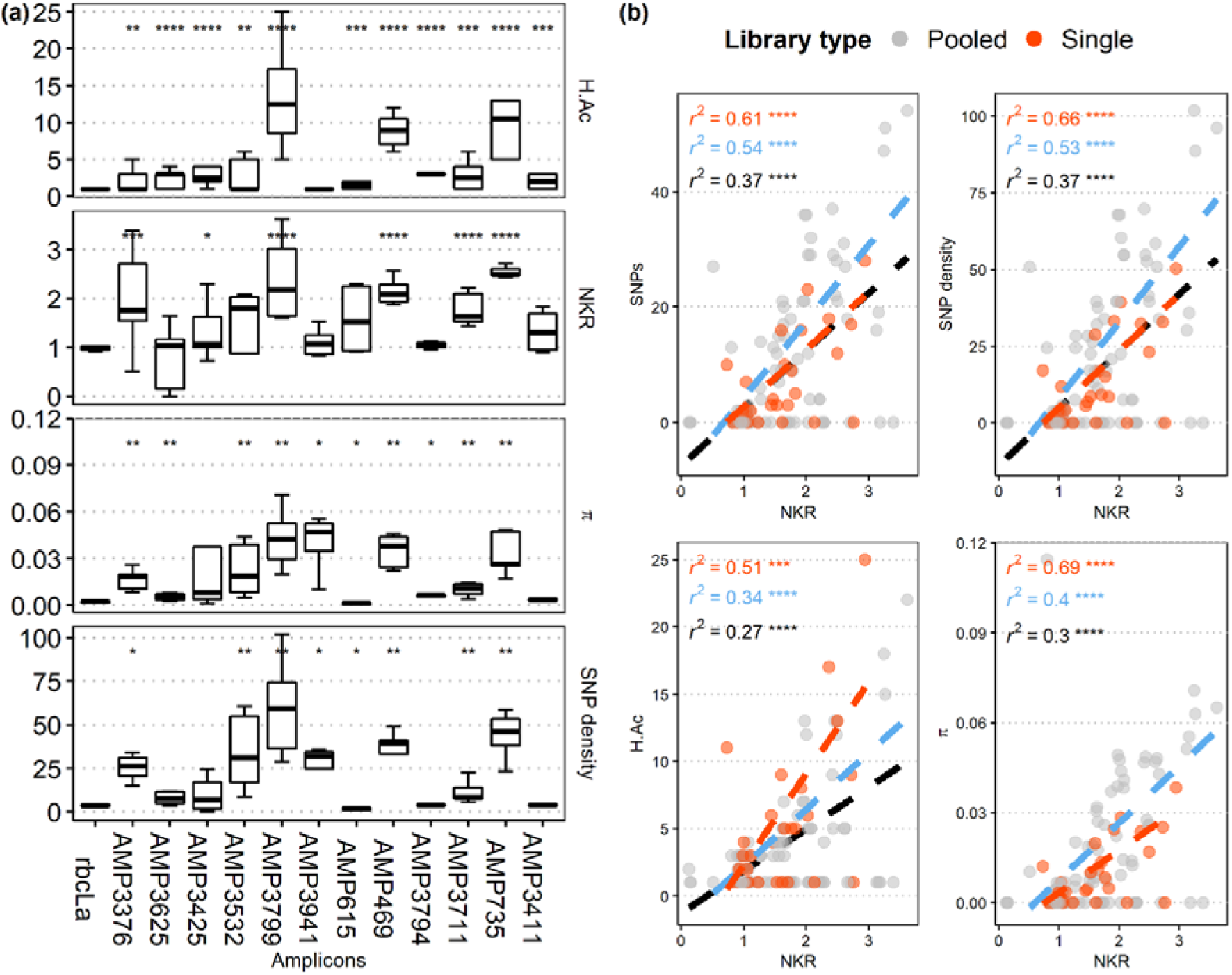
Sequence-based genetic diversity in groups of 16 plants per species. **(a)** SNP-based haplotype allele count (H.Ac), normalized *k*-mer richness (NKR), nucleotide diversity (π) and SNP density (SNPs/kbp) at each locus in groups of sixteen plants per species. Amplicons without SNP calls were assumed to have only one haplotype allele in all sixteen plants. Wilcoxon test’s significance levels are in relation to *rbcLa*. **(b)** Regression of diversity metrics per amplicon. Red dots show metrics from libraries generated from single-plant sequencing libraries; gray dots show metrics for libraries generated from pooled-plant sequencing libraries. Each dot represents one amplicon in a single- or pooled-plant libraries. The r2 value and *P*-value for all amplicons are shown in black; in blue, the same is shown only for amplicons with SNP calls; in red, only for amplicons from single-plant libraries and with SNP calls. Significance levels for *P*-values: 0.0001 = ****, 0.001 = ***, 0.01 = **, 0.05 = *, ns = non-significant. The upper and bottom ends of boxplots indicate the 3^rd^ and 1^st^ quartiles, respectively. The black line inside of boxplots indicates the median value. The upper and lower whiskers in boxplots indicate the maximum (3^rd^ quartile + 1.5 times the interquartile range) and minimum (1^st^ quartile - 1.5 times the interquartile range) values, respectively.

### Multi-locus analysis

*K*-mer based distances had an overall median of 0.29 (IQR = 0.21 - 0.38). At the species level, median *k*-mer-based distances were 0.34 (IQR = 0.28 - 0.39) for *D. glomerata*, 0.20 (IQR = 0.15 - 0.43) for *F. pratensis*, 0.31 (IQR = 0.22 - 0.40) for *L. perenne*, 0.28 (IQR = 0.19 - 0.39) for *T. pratense*, and 0.24 (IQR = 0.21 - 0.30).

Prevosti’s pairwise distances based on HAP-MLGs had an overall median of 0.25 (IQR = 0.12 - 0.32). At the species level, median distances based on HAP-MLGs were 0.28 (IQR = 0.25 - 0.32) for *D. glomerata*, 0.25 (IQR =0.16 - 0.30) for *F. pratensis*, 0.28 (IQR =0.22 - 0.32) for *L. perenne*, 0.45 (IQR =0.40 - 0.50) for *T. pratense* and 0.09 (0.06 - 0.12) for *T. repens*.

Prevosti’s pairwise distances based on SSR-MLGs had an overall median of 0.60 (IQR = 0.50 - 0.70). At the species level, median distances based on SSR-MLGs were 0.55 (IQR = 0.50 - 0.65) for *D. glomerata*, 0.50 (IQR =0.40 - 0.60) for *F. pratensis*, 0.50 (IQR = 0.40 - 0.55) for *L. perenne*, 0.90 (IQR = 0.83 - 1.00) for *T. pratense* and 0.62 (IQR =0.56 - 0.69) for *T. repens*.

Distances based on SSR-MLGs were larger than both *k*-mer- and HAP-MLGs distances in all species (Fig. 4). In turn, *k*-mer-based distances were significantly larger than HAP-MLGs distances in all species except *F. pratensis* (no significant differences between *k*-mer- and haplotype-based distances, Fig. 4) and *T. pratense* (haplotype-based distances were larger than *k*-mer based, Fig. 4).

**Figure 4.**
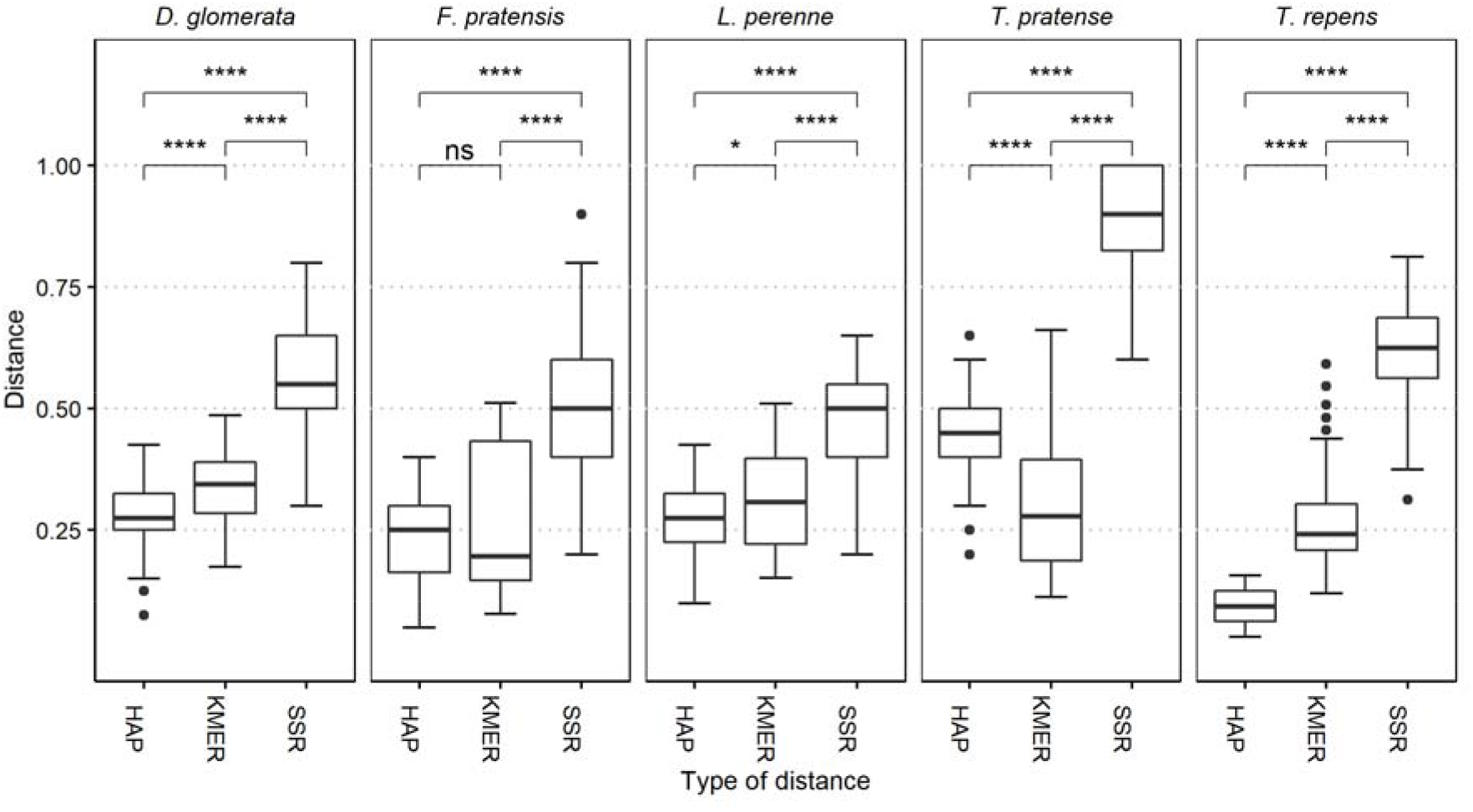
Comparison of pairwise distances derived from single-plant samples using sequence- and SSR-based genetic diversity statistics. Prevosti’s distance was used for amplicon haplotype-based multi-locus genotypes (HAP-MLGs) and SSR-based MLGs (SSR-MLGs). Sourmash-derived distance was used for *k*-mers. Five amplicons per species (four, in case of *T. repens*) were used to calculate distance matrices for HAP-MLGs and *k*-mers. The five most polymorphic SSRs per species were used to calculate the SSR-MLGs distance matrix. Samples with >5% missing loci were removed. Wilcoxon test’s significance levels are shown for all possible comparisons. Significance levels for P-values: 0.0001 = ****, 0.001 = ***, 0.01 = **, 0.05 = *, ns = non-significant. The upper and bottom ends of boxplots indicate the 3^rd^ and 1^st^ quartiles, respectively. The black line inside of boxplots indicates the median value. The upper and lower whiskers in boxplots indicate the maximum (3^rd^ quartile + 1.5 times the interquartile range) and minimum (1^st^ quartile - 1.5 times the interquartile range) values, respectively. Dots on top and at the bottom of boxplots indicate outliers.

In grasses, *k*-mer- and haplotype-based distances are very similar to each other (i.e., their ratio is close to 1; Table 8), whereas in legumes, they are more dissimilar (i.e., their ratio deviates from 1; Table 8).

## Discussion

We observed capture efficiencies (i.e., the proportion of the total sequencing read output that maps on the target loci; Table 2) ranging from 1.58% to 55.82% per species, a range that is in line with previous reports of multispecies sequence capture assays. For example, a study of 25 legume species reported capture efficiencies ranging from 17% to 48% per species (Vatanparast et al., 2018). Another study that analyzed 42 angiosperm species reported a wider range of capture efficiencies per species: 5% to 68.1% (Johnson et al., 2019). Grass and legume species had different capture efficiencies, which is likely due to grasses having larger average genome sizes (4,838.5 Mbp) than legumes (958.4 Mbp; Supplementary Table 7). As libraries with similar concentrations were pooled together disregarding their species in each sequence capture reaction, this could have resulted in a lower effective concentration of target loci for grass samples in the discovery phase of this study. This is supported by a negative linear correlation observed between genome size and capture efficiency (*r*^2^ = 0.58, *P*-value < 0.001). On average and depending on the species, 34.9% to 94.8% of the 611 target loci per species were recovered (Table 2). A similarly high, species-dependent variability of captured loci has been observed in other multispecies studies, from 54% to 98% (Vatanparast et al., 2018) or even 2% to 98% (Johnson et al., 2019).

We expected our estimates of genetic diversity based on the sequence capture assay to be lower than genome-wide genetic diversity estimates of forage crop species. This is because the large proportion of coding sequences and ultra-conserved-like elements in our target loci, which were included to increase the chances of finding multispecies priming sites around polymorphic sequences. SNP density per species (Table 3) were lower than a genome-wide estimate in *L. multiflorum* (31 SNPs/kbp; Knorst et al., 2019) and than an estimate based on genic regions in *L. perenne* (35.6 SNPs/kbp; Ruttink et al., 2015). The SNP densities we found in our sequence capture data (Table 3) are closer to the 17 SNPs/kbp reported for a set of plastomes from a comprehensive sampling of *D. glomerata* from western Europe (Hodkinson et al., 2019).

Three main criteria to choose candidate loci for the validation phase were followed: a) a high NKR value, b) high SNP-based diversity (i.e., high π and SNP density) and c) suitability for multispecies primers with predicted amplicon sizes of 450 ± 50 bp. Such an amplicon length was chosen so the longest possible haplotypes could be generated after merging Illumina MiSeq reads.

NKR and SNP-based diversity measures are two complementary ways to assess diversity. NKR is the count of unique *k*-mer from reads mapping to a locus divided by the locus length. Such measure accounts for insertion/deletion (indel) variants that go undetected by standard SNP calling. As a tool to detect intraspecific variation, *k*-mer richness (or *k*-mer count) analysis has been applied in barley sequences to detect apparent heterozygous mappings, i.e., clusters of divergent sequencing reads that map to genomic loci that are closely related (Pérez-Cantalapiedra et al., 2018). *K*-mer richness analysis has also been applied to infer haplotype diversity in viral populations (Malhotra et al., 2013).

We observed a low to moderate positive correlation between NKR and π per kbp when analyzing each species individually (Fig. 2b), but no significant correlation was present in ungrouped data (Fig. 2a). This is likely because the pseudo-reference assemblies underlying our SNP and *k*-mer analyses were generated on a species-by-species basis. In addition, insertion/deletions (i.e., indel) size variability was high (Supplementary Figure 10), causing poor correlations SNP-based metrics and NKR.

The sequence capture approach of the discovery phase of this work was developed as a steppingstone to find suitable loci for multispecies amplicon sequencing; however, it can *per* se be useful for other applications that justify its relatively high cost. The FORAGE-611 bait set could help elucidating biogeographic patterns in grasses and legume populations. It could also help to resolve phylogenetic relationships in closely related clades, such as the *Festuca-Lolium* complex (Cheng et al., 2016), which comprises some of the most widely cultivated forage grasses in temperate climates It can also be a helpful tool to study intergeneric hybridization and allopolyploidization, a prevalent feature in grasses (Tkach et al., 2020).

In the validation phase of this study, we focused on eleven polymorphic loci and the DNA barcode *rbcLa* (Table 1), which were amplified and sequenced from single- and pooled-plant samples of *D. glomerata, F. pratensis, L. perenne, T. pratense* and *T. repens*. We initially designed a set of >30 candidate multispecies primer pairs targeting >20 loci, but we only picked those with high efficiency in test PCRs (results not shown). More working multispecies primer pairs targeting more loci could be generated by varying amplicon size and by further primer optimization. However, this was out of the scope of this proof-of-concept study.

As expected from amplicon sequencing data, sequencing coverage per locus was very high (>1,000x). Most of the selected amplicons were polymorphic, which made them useful for genetic diversity assessment (Fig. 3a and Table 5). In general, genetic diversity in single- and pooled-plant libraries were mostly similar, with some variation due to the diversity metric and the amplicon.

Among diversity metrics, NKR had the lowest variation due to sample pooling. This highlights the robustness of alignment-free methods for genetic diversity studies. SNP-based metrics (i.e., SNP count, SNP density and π) had higher variations, likely due to the differences between the SNP calling software. PoPoolation produced more SNP calls in the pooled-plant libraries compared to the SNP calls of BCFtools mpileup in their matching single-plant libraries. This is to be expected because PoPoolation false SNP discovery rates are known to increase with depth (Raineri et al., 2012). In the case of SNP-based haplotypes, the low variation haplotype counts stems from the way such haplotypes were generated in pooled-samples (i.e., only the haplotypes that were present in the single-plant libraries were considered).

Among amplicons, those with low variability due to sample pooling (median CV < 10%) were rbcLa, AMP3411, AMP615, AMP3941 and AMP3794, which are also among the least diverse amplicons (Table 7). In the rest of the amplicons, the moderate levels of variability of diversity metrics (median CV between 12.54% to 33.19%) is also likely a result of differences in SNP calling software. However, correlations among amplicon diversity metrics were better than those observed in the discovery phase of this study (Fig. 3b). This is likely because indels in our multispecies amplicons had less size variability than the 611 loci captured in the discovery phase (Supplementary Figure 3), which in turn reduced the discrepancy between NKR and SNP-based metrics.

**Table 7.**
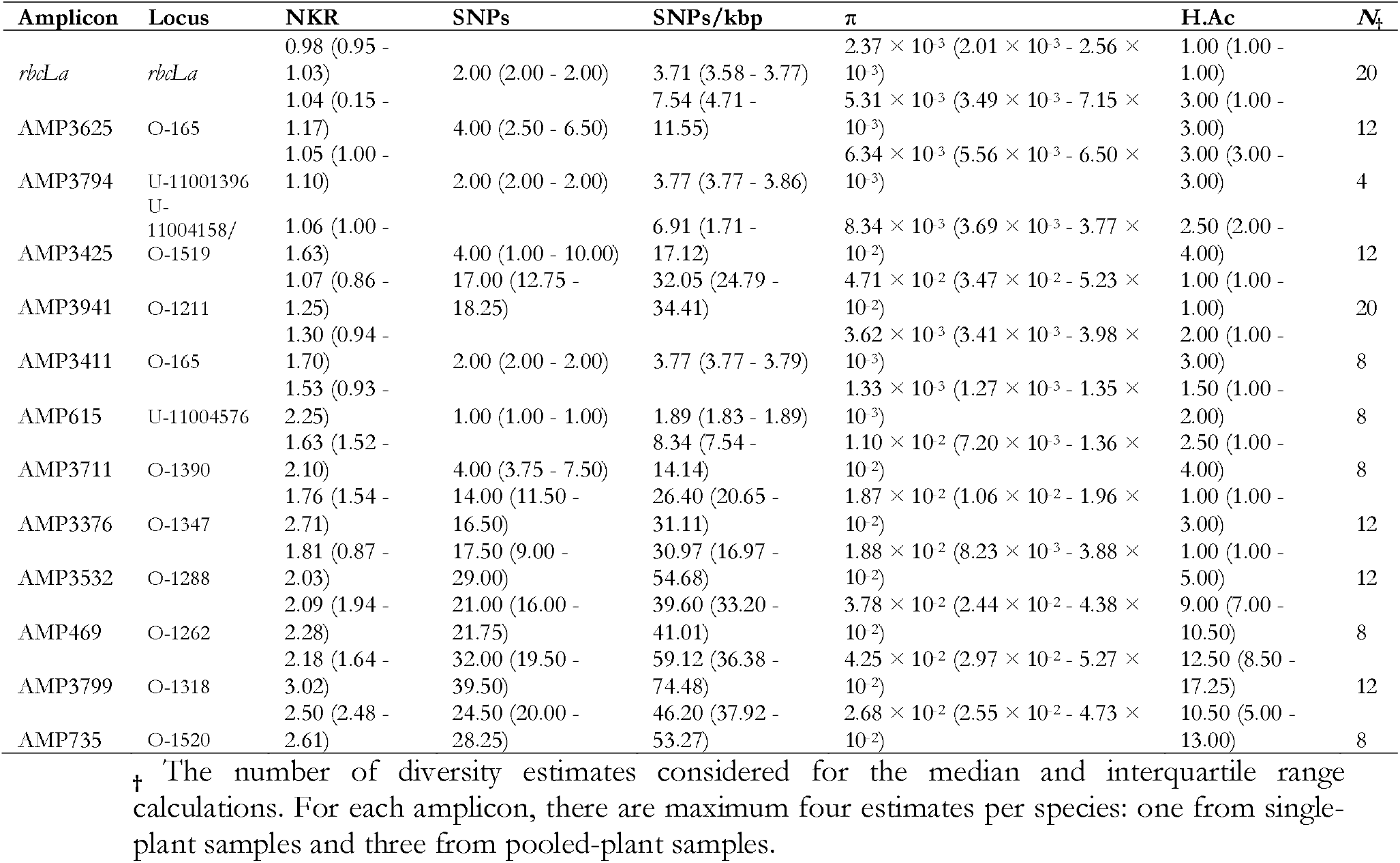
Median genetic diversity statistics per amplicon including single- and pooled- plant samples for the multispecies amplicons and DNA barcode *rbcLa*.

**Table 7.**
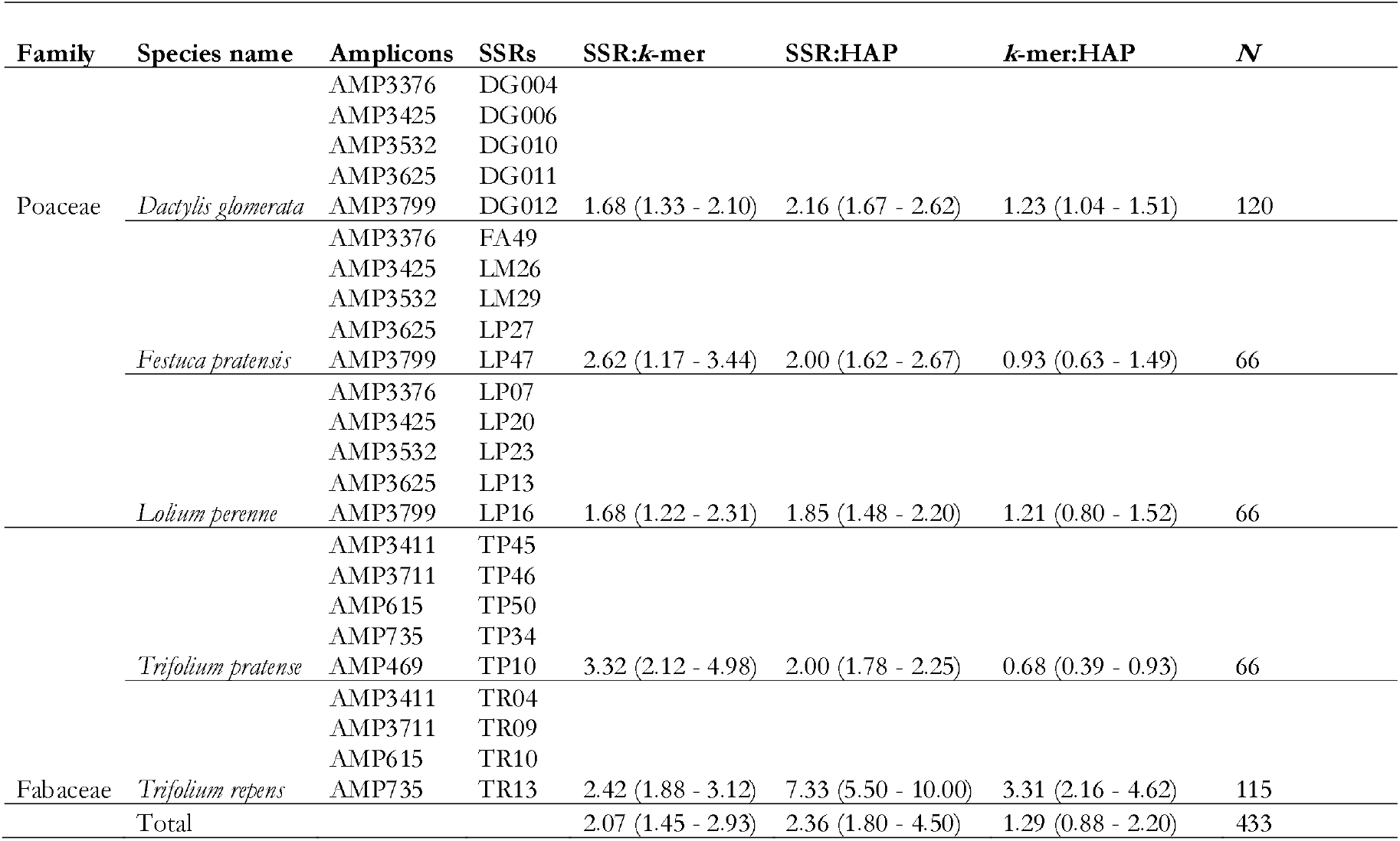
Median ratios between SSR- and amplicon sequencing-based multi-locus pairwise distances.

The average genetic diversity per species (mean π = 8.94×10^−3^; mean SNP density = 12.04 SNPs/kbp; Table 6) was higher than the diversity reported in another targeted sequencing study, which analyzed nine genes putatively involved in flowering (overall π = 7.90×10^−3^; overall SNP density = 7.87 SNPs/kbp; Fiil et al., 2011). Such study was carried out using twenty *L. perenne* genotypes from different European sources (i.e., the “*Lolium* Test Set” or LTS). In contrast, the average genetic diversity per species (Table 6) was lower than those reported for the LTS in eleven pathogen resistance genes (π = 3.14×10^−2^; total SNP density ≈ 100 SNPs/kbp; Xing et al., 2007), which are subject to constant selection pressure and co-evolving alongside pathogens and, therefore, are highly variable. The moderate levels of genetic diversity of our multispecies amplicons compared to resistance genes is likely because our target loci are not involved in a common biological process under evolutionary pressure. Our target loci were picked based on their median within-species diversity across multiple forage species and based on their suitability for multispecies amplification.

To benchmark the diversity estimations from multispecies amplicons, pairwise genetic distances based on k-mers and SNPs were compared to pairwise distances produced with SSRs, a highly polymorphic and multi-allelic marker system. Multispecies amplicons underestimated true pairwise genetic distances by ~50% (Fig. 4 and Table 7). This indicates that more multispecies amplicons are needed to improve their discriminatory power. Theoretically, the discriminatory power of a few tens of SSRs is equivalent to roughly a hundred neutral SNPs (Kalinowski, 2002). However, empirical evidence from an outbreeding plant species shows that the number of SNPs needed to approximate genome-wide diversity estimations is higher than theoretical predictions (Fischer et al., 2017). No correlation was observed between multispecies amplicon sequencing- and SSR-based pairwise distances (data not shown), indicating that multispecies amplicons and SSRs interrogate different parts of each species’ genome, which results in different genetic distance estimations for any given pair of genotypes. This highlights the large diversity present in the genomes of outbreeding forage species.

In conclusion, we showed that multispecies amplicon sequencing is useful for genetic diversity assessments in outbreeding forage crop species. The approach uses inexpensive, unmodified primers and other conventional PCR reagents. By designing multispecies primers, amplicons are transferable across many species of the Poaceae and Fabaceae families, which can further diminish the costs and labor needed for large-scale, multispecies assessments. Compared to other approaches like GBS and RAD-seq, the output of amplicon sequencing is highly enriched in informative data, making it more resource efficient. Furthermore, amplicon sequencing data greatly simplify bioinformatic analyses (M. Sato et al., 2019). We also showed that genetic diversity estimates based on amplicon sequencing of pooled-plant material are similar to those of single-plant material, which further reduces costs per sample. Since PCR is a very robust and versatile method, improvements to the genetic diversity estimates based on multispecies amplicon can be achieved by increasing the number of amplicons analyzed, by varying amplicon sizes, by varying PCR conditions (e.g., multiplex PCR can reduce the processing time needed for each amplicon; Veeckman et al., 2019), or by adapting sequencing approaches (e.g., pool sequencing can be used to characterize larger plant populations).

The tools presented in this study provide the basis for cost-effective, multispecies genetic diversity assessments in grassland plants. In our view, these results represent a promising starting point for further improvements and adaptations. As awareness increases around the ecological significance of plant genetic diversity and as widespread monitoring of genetic diversity gains traction (Hoban et al., 2020; Laikre et al., 2020; Pärli et al., 2021), such multispecies approaches can be valuable additions to the toolset of genetic diversity monitoring efforts.

## Authors’ contributions

R.K. and B.S. conceived the study and provided insights on experimental design and data analysis. M.L.-S. did the laboratory and bioinformatic analyses. All authors read and approved the final manuscript.

## Acknowledgements

We thank Franco Widmer and Christoph Grieder from Agroscope for giving us access to greenhouses and laboratory space. Steven Yates (ETH Zurich) provided the list of orthologous, single-copy genes used for bait design. Thanks to Damian Käch (ETH Zurich), who performed the SSR genotyping of this study as part of his Bachelor thesis. Library preparations and sequencing were performed at the Genetic Diversity Center of ETH Zurich. We thank M. Hardegger and C. Kägi (Swiss Federal Office of Agriculture) for their valuable advice regarding the design of the study. This work was funded by the Swiss Federal Office of Agriculture.

