## Supplementary tables and figures for "A multispecies amplicon sequencing approach for genetic diversity assessment in grassland plant species"

### Discovery phase

#### **Table 1** Plant material used for sequence capture

| Dual-index library | Species | Cultivar(s) | Single or pooled plants |
|---|---|---|---|
| Ap1 | *Alopecurus pratensis* L. | 'Alko' (Saatszucht Steinach, DE) | Single |
| Ap2 | *Alopecurus pratensis* L. | 'Alko' (Saatszucht Steinach, DE) | Single |
| Ap3 | *Alopecurus pratensis* L. | 'Alko' (Saatszucht Steinach, DE) | Single |
| Ap4 | *Alopecurus pratensis* L. | 'Alopex' (DSP/Agroscope, CH) | Single |
| Ap5 | *Alopecurus pratensis* L. | 'Alopex' (DSP/Agroscope, CH) | Single |
| Ap6 | *Alopecurus pratensis* L. | | Pooled (plants 1 - 5) |
| Ae1 | *Arrhenaterum elatius* L. | 'Arone' (Saatszucht Steinach, DE) | Single |
| Ae2 | *Arrhenaterum elatius* L. | 'Arone' (Saatszucht Steinach, DE) | Single |
| Ae3 | *Arrhenaterum elatius* L. | 'Arone' (Saatszucht Steinach, DE) | Single |
| Ae4 | *Arrhenaterum elatius* L. | 'Median' (DLF Životice, CZ) | Single |
| Ae5 | *Arrhenaterum elatius* L. | 'Median' (DLF Životice, CZ) | Single |
| Ae6 | *Arrhenaterum elatius* L. | | Pooled (plants 1 - 5) |
| Cc1 | *Cynosurus cristatus* L. | 'Lena' (HBLFA, AT) | Single |
| Cc2 | *Cynosurus cristatus* L. | 'Lena' (HBLFA, AT) | Single |
| Cc3 | *Cynosurus cristatus* L. | 'Rožnovská' (OSEVA PRO, CZ) | Single |
| Cc4 | *Cynosurus cristatus* L. | 'Rožnovská' (OSEVA PRO, CZ) | Single |
| Cc5 | *Cynosurus cristatus* L. | 'Cresta' (DSP/Agroscope, CH) | Single |
| Cc6 | *Cynosurus cristatus* L. | | Pooled (plants 1 - 5) |
| Dg1 | *Dactylis glomerata* L. | 'Barexcel' (Barenbrug, NL) | Single |
| Dg2 | *Dactylis glomerata* L. | 'Barexcel' (Barenbrug, NL) | Single |
| Dg3 | *Dactylis glomerata* L. | 'Reda' (DSP/Agroscope, CH) | Single |
| Dg4 | *Dactylis glomerata* L. | 'Reda' (DSP/Agroscope, CH) | Single |
| Dg5 | *Dactylis glomerata* L. | 'Brennus' (R2n, FR) | Single |
| Dg6 | *Dactylis glomerata* L. | | Pooled (plants 1 - 5) |
| Fp1 | *Festuca pratensis* Huds. | 'Pradel' (DSP/Agroscope, CH) | Single |
| Fp2 | *Festuca pratensis* Huds. | 'Pradel' (DSP/Agroscope, CH) | Single |
| Fp3 | *Festuca pratensis* Huds. | 'Cosmolit' (Saatszucht Steinach, DE) | Single |
| Fp4 | *Festuca pratensis* Huds. | 'Cosmolit' (Saatszucht Steinach, DE) | Single |
| Fp5 | *Festuca pratensis* Huds. | 'Cosmolit' (Saatszucht Steinach, DE) | Single |
| Fp6 | *Festuca pratensis* Huds. | | Pooled (plants 1 - 5) |
| Fr1 | *Festuca rubra* L. | 'Echo' (DLF-Trifolium, DK) | Single |
| Fr2 | *Festuca rubra* L. | 'Echo' (DLF-Trifolium, DK) | Single |
| Fr3 | *Festuca rubra* L. | 'Pran Solas' (Schweizer, CH) | Single |
| Fr4 | *Festuca rubra* L. | 'Pran Solas' (Schweizer, CH) | Single |
| Fr5 | *Festuca rubra* L. | 'Roland' (Saatszucht Steinach, DE) | Single |
| Fr6 | *Festuca rubra* L. | | Pooled (plants 1 - 5) |
| Lp1 | *Lolium perenne* L. | 'Arara' (DSP/Agroscope, CH) | Single |
| Lp2 | *Lolium perenne* L. | 'Arara' (DSP/Agroscope, CH) | Single |
| Lp3 | *Lolium perenne* L. | 'Artesia' (DSP/Agroscope, CH) | Single |
| Lp4 | *Lolium perenne* L. | 'Artesia' (DSP/Agroscope, CH) | Single |
| Lp5 | *Lolium perenne* L. | 'Arvella' (DSP/Agroscope, CH) | Single |
| Lp6 | *Lolium perenne* L. | | Pooled (plants 1 - 5) |
| Lm1 | *Lolium multiflorum* Lam. | 'Axis' (DSP/Agroscope, CH) | Single |
| Lm2 | *Lolium multiflorum* Lam. | 'Axis' (DSP/Agroscope, CH) | Single |
| Lm3 | *Lolium multiflorum* Lam. | 'Zebra' (DSP/Agroscope, CH) | Single |
| Lm4 | *Lolium multiflorum* Lam. | 'Zebra' (DSP/Agroscope, CH) | Single |
| Lm5 | *Lolium multiflorum* Lam. | 'Caribu' (DSP/Agroscope, CH) | Single |
| Lm6 | *Lolium multiflorum* Lam. | | Pooled (plants 1 - 5) |
| Lc1 | *Lotus corniculatus* L. | 'Polom' (CVRV, VÚRV, CZ) | Single |
| Lc2 | *Lotus corniculatus* L. | 'Polom' (CVRV, VÚRV, CZ) | Single |
| Lc3 | *Lotus corniculatus* L. | 'Lotar' (OSEVA UNI, SK) | Single |
| Lc4 | *Lotus corniculatus* L. | 'Lotar' (OSEVA UNI, SK) | Single |
| Lc5 | *Lotus corniculatus* L. | 'Lotar' (OSEVA UNI, SK) | Single |
| Lc6 | *Lotus corniculatus* L. | | Pooled (plants 1 - 5) |
| Ms1 | *Medicago sativa* L. | 'Sanditi' (Barenbrug, NL) | Single |
| Ms2 | *Medicago sativa* L. | 'Sanditi' (Barenbrug, NL) | Single |
| Ms3 | *Medicago sativa* L. | 'Catera' (Saatszucht Steinach, DE) | Single |
| Ms4 | *Medicago sativa* L. | 'Catera' (Saatszucht Steinach, DE) | Single |
| Ms5 | *Medicago sativa* L. | 'Catera' (Saatszucht Steinach, DE) | Single |
| Ms6 | *Medicago sativa* L. | | Pooled (plants 1 - 5) |
| Ov1 | *Onobrychis viciifolia* Scop. | 'Perdix' (DSP/Agroscope, CH) | Single |
| Ov2 | *Onobrychis viciifolia* Scop. | 'Perdix' (DSP/Agroscope, CH) | Single |
| Ov3 | *Onobrychis viciifolia* Scop. | 'Perly' (DSP/Agroscope, CH) | Single |
| Ov4 | *Onobrychis viciifolia* Scop. | 'Perly' (DSP/Agroscope, CH) | Single |
| Ov5 | *Onobrychis viciifolia* Scop. | 'Perdix' (DSP/Agroscope, CH) | Single |
| Ov6 | *Onobrychis viciifolia* Scop. | | Pooled (plants 1 - 5) |
| Ph1 | *Phleum pratense* L. | 'Tiller' (DLF-Trifolium, DK) | Single |
| Ph2 | *Phleum pratense* L. | 'Tiller' (DLF-Trifolium, DK) | Single |
| Ph3 | *Phleum pratense* L. | 'Toro' (CRA-FLC, IT) | Single |
| Ph4 | *Phleum pratense* L. | 'Toro' (CRA-FLC, IT) | Single |
| Ph5 | *Phleum pratense* L. | 'Anjo' (ILVO, BE) | Single |
| Ph6 | *Phleum pratense* L. | | Pooled (plants 1 - 5) |
| Pp1 | *Poa pratensis* L. | 'Likollo' (DSV, DE) | Single |
| Pp2 | *Poa pratensis* L. | 'Nixe' (Saatszucht Steinach, DE) | Single |
| Pp3 | *Poa pratensis* L. | 'Tommy' (DLF-Trifolium, DK) | Single |

| | | | |
|---|---|---|---|
| Pp4 | *Poa pratensis* L. | 'Tommy' (DLF-Trifolium, DK) | Single |
| Pp5 | *Poa pratensis* L. | 'Tommy' (DLF-Trifolium, DK) | Single |
| Pp6 | *Poa pratensis* L. | | Pooled (plants 1 - 5) |
| Tp1 | *Trifolium pratense* L. | 'Diplomat' (DSV, DE) | Single |
| Tp2 | *Trifolium pratense* L. | 'Pavo' (DSP/Agroscope, CH) | Single |
| Tp3 | *Trifolium pratense* L. | 'Pavo' (DSP/Agroscope, CH) | Single |
| Tp4 | *Trifolium pratense* L. | 'Bonus' (Selgen, CZ) | Single |
| Tp5 | *Trifolium pratense* L. | 'Bonus' (Selgen, CZ) | Single |
| Tp6 | *Trifolium pratense* L. | | Pooled (plants 1 - 5) |
| Tr1 | *Trifolium repens* L. | 'Beaumont' (CW 090; Barenbrug, NL) | Single |
| Tr2 | *Trifolium repens* L. | 'Beaumont' (CW 090; Barenbrug, NL) | Single |
| Tr3 | *Trifolium repens* L. | 'Hebe' (Svalöf-Weibull, SE) | Single |
| Tr4 | *Trifolium repens* L. | 'Hebe' (Svalöf-Weibull, SE) | Single |
| Tr5 | *Trifolium repens* L. | 'Bombus' (DSP/Agroscope, CH) | Single |
| Tr6 | *Trifolium repens* L. | | Pooled (plants 1 - 5) |
| Tf1 | *Trisetum flavescens* L. | 'Gunther' (HBLFA, AT) | Single |
| Tf2 | *Trisetum flavescens* L. | 'Gunther' (HBLFA, AT) | Single |
| Tf3 | *Trisetum flavescens* L. | 'Trisett51' (Saatszucht Steinach, DE) | Single |
| Tf4 | *Trisetum flavescens* L. | 'Trisett51' (Saatszucht Steinach, DE) | Single |
| Tf5 | *Trisetum flavescens* L. | 'Gunther' (HBLFA, AT) | Single |
| Tf6 | *Trisetum flavescens* L. | | Pooled (plants 1 - 5) |

**Table 2** Captured loci and their corresponding genes in the reference genomes of three model species

| | Locus | *A. thaliana* | *B. distachyon* | *M. truncatula* |
|---|---|---|---|---|
| 1 | **ortho-1003** | NA | NA | NA |
| 2 | **ortho-1004** | AT5G15802 | Bradi1g59990 | NA |
| 3 | **ortho-1005** | AT5G43650 | Bradi1g09177 | NA |
| 4 | **ortho-1006** | AT5G47090 | Bradi1g51340 | NA |
| 5 | **ortho-1007** | AT2G37240 | Bradi1g65880 | NA |
| 6 | **ortho-1008** | AT4G17380 | Bradi1g27377 | NA |
| 7 | **ortho-1009** | AT1G50000 | Bradi1g09870 | NA |
| 8 | **ortho-1011** | AT5G49030 | Bradi3g57220 | NA |
| 9 | **ortho-1014** | AT5G48440 | Bradi4g35087 | Medtr7g051240 |
| 10 | **ortho-1016** | AT5G21140 | Bradi4g43810 | NA |
| 11 | **ortho-1019** | AT1G10520 | Bradi1g44820 | Medtr5g040170 |
| 12 | **ortho-1019** | AT1G10520 | Bradi1g44820 | Medtr7g039450 |
| 13 | **ortho-1022** | AT3G58530 | Bradi3g10360 | NA |
| 14 | **ortho-1024** | AT3G26750 | Bradi3g02932 | NA |
| 15 | **ortho-1025** | AT2G31290 | Bradi1g08950 | NA |
| 16 | **ortho-1028** | AT2G20980 | Bradi4g36530 | NA |
| 17 | **ortho-1029** | AT3G08920 | Bradi1g65600 | Medtr4g015860 |
| 18 | **ortho-1030** | AT4G35980 | Bradi4g42850 | NA |
| 19 | **ortho-1031** | AT1G63270 | Bradi5g22580 | Medtr4g131330 |
| 20 | **ortho-1032** | AT1G33810 | Bradi3g13560 | NA |
| 21 | **ortho-1033** | AT1G48200 | Bradi1g50850 | Medtr1g073190 |
| 22 | **ortho-1035** | AT3G10850 | Bradi1g63410 | NA |
| 23 | **ortho-1037** | AT1G63855 | Bradi3g01870 | Medtr2g078490 |
| 24 | **ortho-1039** | AT5G45680 | NA | NA |
| 25 | **ortho-104** | NA | Bradi3g53987 | Medtr3g032530 |
| 26 | **ortho-1041** | AT2G36740 | Bradi3g12990 | NA |
| 27 | **ortho-1042** | AT4G09140 | Bradi2g61370 | Medtr3g005810 |
| 28 | **ortho-1043** | AT2G43235 | Bradi2g30480 | Medtr1g116540 |
| 29 | **ortho-1044** | AT5G03770 | Bradi2g55417 | NA |
| 30 | **ortho-1045** | AT4G04870 | Bradi2g52187 | NA |
| 31 | **ortho-1046** | AT5G03070 | Bradi2g00320 | NA |
| 32 | **ortho-1049** | AT3G08490 | Bradi1g69790 | Medtr3g033530 |
| 33 | **ortho-1052** | AT3G04650 | Bradi1g23880 | NA |
| 34 | **ortho-1053** | AT4G16510 | Bradi3g50910 | NA |
| 35 | **ortho-1054** | AT5G63700 | Bradi2g25017 | NA |
| 36 | **ortho-1055** | AT2G20360 | Bradi3g55340 | Medtr1g069455 |
| 37 | **ortho-1056** | AT3G48425 | Bradi4g39120 | Medtr3g075220 |
| 38 | **ortho-106** | AT4G14330 | Bradi3g54890 | Medtr1g075680 |
| 39 | **ortho-1060** | AT2G39725 | Bradi3g19714 | NA |
| 40 | **ortho-1061** | AT3G18630 | Bradi5g25880 | NA |
| 41 | **ortho-1063** | AT5G20220 | Bradi2g48140 | Medtr8g468600 |
| 42 | **ortho-1064** | AT5G02250 | Bradi2g37760 | Medtr4g021750 |
| 43 | **ortho-1066** | AT4G02790 | Bradi2g24370 | Medtr7g094890 |
| 44 | **ortho-1067** | AT5G51880 | Bradi1g77270 | Medtr1g101840 |
| 45 | **ortho-1069** | AT3G20060 | Bradi2g10220 | Medtr5g009020 |
| 46 | **ortho-1069** | AT1G50490 | Bradi2g10220 | Medtr5g009020 |
| 47 | **ortho-1071** | AT5G18390 | Bradi3g40880 | NA |
| 48 | **ortho-1072** | AT1G03750 | NA | NA |
| 49 | **ortho-1072** | AT1G03760 | NA | NA |
| 50 | **ortho-1073** | AT3G15180 | Bradi2g17000 | Medtr8g061280 |
| 51 | **ortho-1074** | AT1G73820 | Bradi4g41610 | NA |
| 52 | **ortho-1076** | AT1G71310 | Bradi2g56682 | NA |
| 53 | **ortho-1078** | AT4G16210 | Bradi1g64320 | Medtr4g122220 |
| 54 | **ortho-1079** | AT2G25280 | NA | NA |
| 55 | **ortho-1082** | AT2G35900 | Bradi2g51240 | NA |
| 56 | **ortho-1084** | AT3G12040 | Bradi3g57425 | Medtr7g093540 |
| 57 | **ortho-1085** | AT5G38530 | Bradi1g35600 | NA |
| 58 | **ortho-1086** | AT1G06240 | Bradi2g06920 | Medtr2g091140 |
| 59 | **ortho-1087** | AT3G01920 | Bradi3g41240 | Medtr6g045607 |
| 60 | **ortho-1089** | AT4G01270 | Bradi3g17130 | NA |
| 61 | **ortho-1092** | AT1G53530 | Bradi4g13337 | Medtr2g072450 |
| 62 | **ortho-1095** | AT5G05310 | Bradi4g25590 | Medtr1g007630 |
| 63 | **ortho-1097** | AT4G37020 | Bradi3g23267 | NA |
| 64 | **ortho-1098** | AT3G04890 | Bradi1g54380 | NA |
| 65 | **ortho-1099** | AT5G63100 | Bradi3g11150 | NA |
| 66 | **ortho-1100** | AT1G51110 | Bradi1g28310 | NA |
| 67 | **ortho-1102** | AT3G14580 | Bradi5g21920 | NA |
| 68 | **ortho-1103** | AT5G65000 | Bradi3g46580 | Medtr8g090230 |
| 69 | **ortho-1105** | AT5G17460 | Bradi2g01810 | Medtr5g086010 |
| 70 | **ortho-1105** | AT5G17460 | Bradi2g01810 | Medtr5g087440 |
| 71 | **ortho-1106** | AT1G66510 | NA | Medtr6g003980 |
| 72 | **ortho-1106** | AT1G66520 | NA | Medtr6g003980 |
| 73 | **ortho-1107** | AT1G52530 | NA | Medtr1g082730 |
| 74 | **ortho-1109** | AT3G55160 | Bradi3g15920 | Medtr6g011760 |
| 75 | **ortho-1110** | AT3G20440 | Bradi1g41970 | Medtr3g053220 |
| 76 | **ortho-1112** | AT1G18030 | Bradi4g27880 | Medtr6g081850 |
| 77 | **ortho-1113** | AT3G25120 | Bradi5g07180 | Medtr8g012725 |
| 78 | **ortho-1114** | AT2G40760 | Bradi2g44260 | Medtr5g067730 |
| 79 | **ortho-1115** | AT1G71500 | Bradi4g21270 | NA |
| 80 | **ortho-1116** | AT4G37200 | Bradi1g07470 | Medtr5g018580 |
| 81 | **ortho-1118** | AT3G51930 | Bradi2g60607 | NA |
| 82 | **ortho-1120** | AT1G08840 | Bradi5g19741 | NA |
| 83 | **ortho-1122** | AT5G14220 | Bradi5g14120 | NA |
| 84 | **ortho-1124** | AT4G18460 | Bradi2g08490 | Medtr2g036040 |
| 85 | **ortho-1127** | AT1G52590 | NA | Medtr2g103590 |
| 86 | **ortho-1130** | AT1G49870 | Bradi1g14877 | NA |
| 87 | **ortho-1132** | AT5G51170 | Bradi3g50320 | NA |
| 88 | **ortho-1133** | AT5G05200 | Bradi2g36950 | Medtr5g068050 |
| 89 | **ortho-1136** | AT2G30105 | Bradi3g27860 | Medtr3g465570 |
| 90 | **ortho-1138** | AT2G36370 | Bradi2g61920 | Medtr7g104170 |
| 91 | **ortho-1139** | AT2G31040 | Bradi2g46306 | Medtr4g079850 |
| 92 | **ortho-1140** | AT3G56310 | Bradi1g17730 | Medtr3g094160 |
| 93 | **ortho-1146** | AT5G07960 | Bradi1g08210 | Medtr3g024170 |
| 94 | **ortho-1146** | AT5G07960 | Bradi1g08210 | Medtr3g024190 |
| 95 | **ortho-1146** | AT5G07960 | Bradi1g08201 | Medtr3g024170 |
| 96 | **ortho-1146** | AT5G07960 | Bradi1g08201 | Medtr3g024190 |
| 97 | **ortho-1148** | AT2G18710 | Bradi3g19100 | Medtr2g460600 |
| 98 | **ortho-1150** | AT2G46910 | Bradi3g30740 | NA |
| 99 | **ortho-1151** | AT1G69523 | Bradi3g59480 | Medtr5g041780 |
| 100 | **ortho-1153** | AT3G20970 | NA | Medtr1g069215 |
| 101 | **ortho-1154** | AT5G48545 | Bradi4g39695 | NA |
| 102 | **ortho-1156** | AT5G57960 | Bradi4g14850 | Medtr1g022980 |
| 103 | **ortho-1158** | AT1G56050 | Bradi1g06772 | NA |
| 104 | **ortho-1159** | AT1G31190 | Bradi3g05226 | NA |
| 105 | **ortho-1161** | NA | Bradi1g11700 | Medtr7g117040 |
| 106 | **ortho-1163** | AT1G03250 | Bradi4g42940 | NA |
| 107 | **ortho-1170** | AT5G41960 | Bradi5g19590 | NA |
| 108 | **ortho-1173** | AT1G74530 | Bradi3g15890 | NA |
| 109 | **ortho-1174** | AT5G10730 | Bradi4g08550 | Medtr3g105660 |
| 110 | **ortho-1175** | AT3G20480 | Bradi1g59740 | Medtr3g052590 |
| 111 | **ortho-1178** | AT3G54970 | NA | Medtr1g103900 |
| 112 | **ortho-1185** | AT5G19850 | Bradi1g23675 | Medtr5g056200 |
| 113 | **ortho-1186** | AT3G25410 | Bradi3g43450 | Medtr7g053090 |
| 114 | **ortho-1187** | AT1G56180 | Bradi3g05210 | Medtr2g066060 |
| 115 | **ortho-1188** | AT3G24820 | Bradi5g24967 | NA |
| 116 | **ortho-119** | NA | Bradi4g02900 | NA |
| 117 | **ortho-1190** | AT3G15160 | Bradi5g22560 | Medtr4g081050 |
| 118 | **ortho-1191** | AT1G67660 | Bradi5g13880 | NA |
| 119 | **ortho-1192** | AT5G22050 | Bradi1g57100 | NA |
| 120 | **ortho-1194** | AT4G00060 | Bradi1g32300 | Medtr4g027425 |
| 121 | **ortho-1195** | AT1G72320 | Bradi4g25530 | NA |
| 122 | **ortho-1196** | AT2G05830 | Bradi4g22150 | NA |
| 123 | **ortho-1198** | AT5G39940 | Bradi4g00497 | Medtr6g011270 |
| 124 | **ortho-1199** | AT5G13680 | Bradi1g24690 | Medtr1g115300 |
| 125 | **ortho-1200** | AT2G38130 | Bradi2g32930 | Medtr7g084140 |
| 126 | **ortho-1201** | AT1G44920 | Bradi1g09940 | Medtr1g017450 |
| 127 | **ortho-1202** | AT4G33945 | Bradi3g58190 | Medtr1g034000 |
| 128 | **ortho-1203** | AT5G44740 | Bradi2g50340 | NA |
| 129 | **ortho-1203** | AT5G44740 | Bradi2g50350 | NA |
| 130 | **ortho-1204** | AT5G13510 | Bradi1g65900 | Medtr4g131540 |
| 131 | **ortho-1205** | AT5G54880 | Bradi1g15740 | Medtr5g092280 |
| 132 | **ortho-1206** | AT5G51130 | Bradi3g41780 | Medtr8g102950 |
| 133 | **ortho-1208** | AT1G54650 | Bradi5g05210 | NA |
| 134 | **ortho-1209** | AT2G21620 | Bradi3g52780 | Medtr3g116830 |
| 135 | **ortho-1210** | AT3G10220 | Bradi5g27560 | NA |
| 136 | **ortho-1211** | AT5G61540 | Bradi5g17580 | Medtr3g008180 |
| 137 | **ortho-1212** | AT5G15170 | Bradi1g25830 | Medtr7g050860 |
| 138 | **ortho-1213** | AT5G52970 | Bradi5g03590 | NA |
| 139 | **ortho-1214** | NA | Bradi2g31930 | NA |
| 140 | **ortho-1217** | AT3G23400 | Bradi4g14630 | Medtr1g075660 |
| 141 | **ortho-1218** | AT5G63910 | Bradi5g27617 | Medtr3g096675 |
| 142 | **ortho-1219** | AT5G02710 | Bradi1g61764 | Medtr0016s0020 |
| 143 | **ortho-1219** | AT5G02710 | Bradi1g61764 | Medtr1g067330 |
| 144 | **ortho-1220** | AT3G21350 | Bradi1g45820 | Medtr8g015230 |
| 145 | **ortho-1221** | AT1G71810 | Bradi5g23450 | NA |
| 146 | **ortho-1222** | AT5G67170 | Bradi5g25650 | Medtr4g094860 |
| 147 | **ortho-1225** | AT1G59600 | Bradi1g23140 | NA |
| 148 | **ortho-1226** | AT5G22110 | Bradi3g37840 | NA |
| 149 | **ortho-1228** | AT5G64010 | NA | NA |
| 150 | **ortho-1230** | AT5G57655 | Bradi1g18550 | Medtr4g128840 |
| 151 | **ortho-1231** | AT4G17760 | Bradi1g50020 | Medtr8g447310 |
| 152 | **ortho-1236** | AT3G23490 | Bradi3g28470 | NA |
| 153 | **ortho-1237** | AT2G39140 | Bradi1g74850 | NA |
| 154 | **ortho-1238** | AT5G38520 | Bradi2g25130 | Medtr1g076920 |
| 155 | **ortho-1239** | AT4G01040 | Bradi2g43770 | Medtr5g025480 |
| 156 | **ortho-1241** | AT2G25530 | Bradi2g52350 | NA |
| 157 | **ortho-1241** | AT3G15290 | Bradi2g52350 | NA |

| | | | | |
|---|---|---|---|---|
| 158 | ortho-1242 | AT1G59840 | Bradi5g24257 | Medtr5g037050 |
| 159 | ortho-1244 | AT1G16540 | Bradi1g32350 | Medtr4g030930 |
| 160 | ortho-1245 | AT4G15240 | Bradi3g14530 | NA |
| 161 | ortho-1246 | AT3G03560 | Bradi1g32732 | Medtr8g010360 |
| 162 | ortho-1248 | AT5G15390 | Bradi1g22607 | NA |
| 163 | ortho-1253 | AT2G39440 | Bradi3g05170 | NA |
| 164 | ortho-1258 | AT2G45990 | Bradi2g27310 | Medtr7g077870 |
| 165 | ortho-1259 | AT5G39830 | Bradi5g12370 | NA |
| 166 | ortho-1261 | AT4G33460 | Bradi2g51200 | NA |
| 167 | ortho-1262 | AT3G07080 | Bradi1g32927 | Medtr7g058520 |
| 168 | ortho-1263 | AT4G01860 | Bradi1g77780 | Medtr7g110260 |
| 169 | ortho-1265 | AT5G64830 | Bradi5g23500 | Medtr3g095020 |
| 170 | ortho-1267 | AT1G44414 | Bradi2g24230 | Medtr1g029990 |
| 171 | ortho-1269 | AT3G19800 | Bradi5g16200 | NA |
| 172 | ortho-1270 | AT3G50685 | NA | Medtr5g019920 |
| 173 | ortho-1271 | AT1G11880 | Bradi4g06960 | Medtr4g084070 |
| 174 | ortho-1272 | AT5G63000 | Bradi2g10170 | Medtr4g105570 |
| 175 | ortho-1273 | AT5G20990 | Bradi5g24930 | Medtr5g087490 |
| 176 | ortho-1274 | AT4G31600 | Bradi1g46960 | Medtr3g109070 |
| 177 | ortho-1278 | AT5G45380 | Bradi3g21640 | Medtr5g026640 |
| 178 | ortho-1279 | AT2G33385 | Bradi5g15480 | NA |
| 179 | ortho-1281 | AT2G04740 | Bradi1g42730 | NA |
| 180 | ortho-1282 | AT2G04845 | Bradi1g71575 | Medtr3g083200 |
| 181 | ortho-1283 | AT2G04842 | Bradi3g53530 | Medtr3g083180 |
| 182 | ortho-1286 | AT5G38660 | Bradi3g22030 | NA |
| 183 | ortho-1288 | AT5G38460 | Bradi1g63540 | NA |
| 184 | ortho-1289 | AT2G22530 | Bradi3g60040 | Medtr5g032210 |
| 185 | ortho-1290 | AT5G26040 | Bradi1g37510 | NA |
| 186 | ortho-1291 | AT2G47980 | NA | NA |
| 187 | ortho-1293 | AT4G14110 | Bradi5g10090 | NA |
| 188 | ortho-1297 | AT2G38680 | Bradi1g13590 | Medtr1g090380 |
| 189 | ortho-1299 | AT5G26230 | Bradi2g26620 | NA |
| 190 | ortho-1300 | AT2G44520 | NA | NA |
| 191 | ortho-1302 | AT4G23890 | Bradi3g21280 | Medtr7g076620 |
| 192 | ortho-1303 | AT1G14150 | Bradi3g46880 | Medtr5g041910 |
| 193 | ortho-1304 | AT4G16530 | Bradi3g30370 | NA |
| 194 | ortho-1306 | AT2G28600 | Bradi4g26640 | NA |
| 195 | ortho-1313 | AT2G43950 | Bradi3g48410 | NA |
| 196 | ortho-1315 | AT1G06560 | Bradi3g21700 | Medtr5g091110 |
| 197 | ortho-1316 | AT3G05410 | Bradi2g59950 | Medtr1g083110 |
| 198 | ortho-1318 | AT5G04050 | Bradi1g31400 | NA |
| 199 | ortho-1322 | AT1G08220 | Bradi2g34156 | NA |
| 200 | ortho-1322 | AT1G08220 | Bradi2g34159 | NA |
| 201 | ortho-1326 | AT5G65110 | Bradi4g14090 | NA |
| 202 | ortho-1328 | AT1G76130 | Bradi5g08800 | Medtr1g026170 |
| 203 | ortho-1329 | AT5G63440 | NA | NA |
| 204 | ortho-1331 | AT2G36885 | Bradi1g75900 | NA |
| 205 | ortho-1332 | AT5G24750 | Bradi3g07997 | NA |
| 206 | ortho-1333 | AT1G34380 | Bradi2g56537 | NA |
| 207 | ortho-1334 | AT3G46550 | Bradi2g54620 | NA |
| 208 | ortho-1335 | AT3G54690 | Bradi3g04440 | NA |
| 209 | ortho-1336 | AT3G59300 | Bradi2g57730 | Medtr5g090150 |
| 210 | ortho-1337 | AT2G03390 | Bradi2g50810 | NA |
| 211 | ortho-1338 | AT5G14100 | Bradi4g19047 | NA |
| 212 | ortho-1339 | AT1G25375 | Bradi2g02250 | Medtr1g110300 |
| 213 | ortho-1340 | AT5G60590 | Bradi1g71070 | Medtr4g459430 |
| 214 | ortho-1341 | AT3G52040 | Bradi4g23700 | Medtr3g034210 |
| 215 | ortho-1341 | AT3G52030 | Bradi4g23700 | Medtr3g034210 |
| 216 | ortho-1342 | AT3G04480 | Bradi3g54847 | Medtr7g111650 |
| 217 | ortho-1343 | AT5G17560 | Bradi1g42030 | NA |
| 218 | ortho-1344 | AT2G41760 | Bradi2g26900 | Medtr7g034355 |
| 219 | ortho-1346 | AT3G13180 | Bradi4g33067 | Medtr5g096020 |
| 220 | ortho-1347 | AT2G44020 | Bradi3g60150 | Medtr1g115415 |
| 221 | ortho-1348 | AT5G50290 | Bradi3g39960 | Medtr4g105550 |
| 222 | ortho-1348 | AT5G50290 | Bradi3g39950 | Medtr4g105550 |
| 223 | ortho-1349 | AT3G48900 | Bradi3g12930 | NA |
| 224 | ortho-1352 | AT2G39910 | Bradi1g33657 | NA |
| 225 | ortho-1353 | AT2G38025 | Bradi1g67110 | Medtr4g014750 |
| 226 | ortho-1357 | AT3G07670 | Bradi3g56450 | Medtr3g082690 |
| 227 | ortho-1359 | AT5G13520 | Bradi1g03860 | Medtr1g116230 |
| 228 | ortho-1360 | AT4G09620 | Bradi1g02400 | NA |
| 229 | ortho-1361 | AT4G03150 | NA | Medtr7g107680 |
| 230 | ortho-1361 | AT4G03140 | NA | Medtr7g107680 |
| 231 | ortho-1363 | AT2G34860 | Bradi3g37770 | NA |
| 232 | ortho-1364 | AT4G16570 | Bradi1g52270 | NA |
| 233 | ortho-1365 | AT5G46400 | Bradi3g35037 | NA |
| 234 | ortho-1366 | AT5G51540 | Bradi1g33210 | Medtr3g009160 |
| 235 | ortho-1367 | AT4G28740 | Bradi5g15130 | Medtr1g069240 |
| 236 | ortho-1369 | AT5G11840 | Bradi2g18310 | Medtr1g016360 |
| 237 | ortho-1372 | AT2G26780 | Bradi4g43517 | Medtr5g089000 |
| 238 | ortho-1375 | AT1G16650 | Bradi4g02070 | Medtr2g064450 |
| 239 | ortho-1380 | AT3G47610 | Bradi4g30160 | Medtr4g108590 |
| 240 | ortho-1381 | AT3G24030 | Bradi2g12822 | Medtr1g046510 |
| 241 | ortho-1385 | AT2G38920 | Bradi1g13500 | Medtr7g107060 |
| 242 | ortho-1390 | AT1G08460 | Bradi2g24020 | Medtr2g087270 |
| 243 | ortho-1394 | AT2G04039 | Bradi1g21780 | Medtr8g028100 |
| 244 | ortho-1396 | AT2G46200 | Bradi2g07530 | NA |
| 245 | ortho-1397 | AT1G67700 | Bradi3g47890 | NA |
| 246 | ortho-1399 | AT1G27050 | Bradi2g51650 | NA |
| 247 | ortho-1400 | AT5G04360 | Bradi5g00540 | Medtr1g090693 |
| 248 | ortho-1401 | AT1G17410 | Bradi3g46220 | Medtr8g021233 |
| 249 | ortho-1402 | AT4G02990 | NA | Medtr5g068860 |
| 250 | ortho-1404 | AT2G45520 | Bradi1g49890 | Medtr7g075640 |
| 251 | ortho-1405 | AT3G10060 | Bradi1g54970 | NA |
| 252 | ortho-1406 | AT3G01380 | Bradi3g47034 | NA |
| 253 | ortho-1408 | AT5G14140 | Bradi4g18830 | Medtr7g029090 |
| 254 | ortho-1410 | AT1G49380 | Bradi1g07340 | NA |
| 255 | ortho-1411 | AT1G31860 | Bradi2g10490 | Medtr7g084480 |
| 256 | ortho-1412 | AT3G44380 | Bradi3g42830 | NA |
| 257 | ortho-1413 | AT3G07720 | Bradi2g38270 | NA |
| 258 | ortho-1421 | AT2G21180 | Bradi1g68030 | Medtr4g075500 |
| 259 | ortho-1422 | AT4G39740 | Bradi4g29060 | NA |
| 260 | ortho-1423 | NA | Bradi3g56310 | NA |
| 261 | ortho-1424 | AT5G13760 | Bradi1g19300 | NA |
| 262 | ortho-1425 | AT3G25430 | Bradi2g42017 | NA |
| 263 | ortho-1426 | AT3G15140 | Bradi3g22650 | Medtr8g061030 |
| 264 | ortho-143 | NA | NA | NA |
| 265 | ortho-1431 | AT4G30950 | Bradi3g36920 | Medtr2g460790 |
| 266 | ortho-1433 | AT2G07170 | Bradi4g37910 | Medtr6g093150 |
| 267 | ortho-1434 | AT5G07330 | Bradi4g17200 | NA |
| 268 | ortho-1435 | AT3G22450 | Bradi3g29191 | Medtr5g006310 |
| 269 | ortho-1440 | AT1G12250 | Bradi3g50060 | Medtr5g012110 |
| 270 | ortho-1441 | AT4G01570 | Bradi1g11480 | NA |
| 271 | ortho-1443 | AT4G00030 | Bradi5g17740 | NA |
| 272 | ortho-1446 | AT3G01980 | Bradi1g01780 | Medtr1g007800 |
| 273 | ortho-145 | NA | NA | NA |
| 274 | ortho-1450 | AT2G03667 | Bradi4g03827 | Medtr4g083180 |
| 275 | ortho-1451 | AT5G16630 | Bradi3g36277 | NA |
| 276 | ortho-1453 | AT3G52390 | Bradi1g57200 | NA |
| 277 | ortho-1455 | AT1G77930 | Bradi1g20730 | Medtr3g073560 |
| 278 | ortho-1456 | AT4G16700 | Bradi1g78460 | Medtr5g012880 |
| 279 | ortho-1457 | AT5G08310 | Bradi2g31210 | Medtr1g114190 |
| 280 | ortho-1459 | AT1G12244 | Bradi3g33090 | Medtr4g027430 |
| 281 | ortho-1462 | AT1G28100 | NA | Medtr2g072940 |
| 282 | ortho-1462 | AT1G28110 | NA | Medtr2g072940 |
| 283 | ortho-1468 | AT5G12130 | Bradi2g37780 | NA |
| 284 | ortho-1469 | AT2G28560 | Bradi2g37817 | NA |
| 285 | ortho-147 | AT5G63960 | NA | Medtr3g096525 |
| 286 | ortho-1471 | AT4G15520 | Bradi2g36660 | Medtr4g118420 |
| 287 | ortho-1473 | AT1G53250 | Bradi2g14670 | NA |
| 288 | ortho-1474 | AT2G43110 | Bradi5g05700 | NA |
| 289 | ortho-1475 | AT3G24200 | Bradi1g02380 | Medtr7g051960 |
| 290 | ortho-1475 | AT3G24200 | Bradi1g02380 | Medtr2g060550 |
| 291 | ortho-1476 | AT3G18524 | Bradi1g15260 | Medtr4g111945 |
| 292 | ortho-1478 | AT4G13330 | Bradi5g15402 | NA |
| 293 | ortho-1479 | AT5G66005 | Bradi1g76380 | Medtr5g024200 |
| 294 | ortho-1480 | AT2G45270 | Bradi2g11360 | NA |
| 295 | ortho-1482 | AT1G11900 | NA | NA |
| 296 | ortho-1483 | AT1G14205 | Bradi1g73720 | NA |
| 297 | ortho-1484 | AT3G01800 | Bradi5g14160 | Medtr8g040070 |
| 298 | ortho-1486 | AT3G63390 | Bradi4g13770 | Medtr7g095740 |
| 299 | ortho-1487 | AT1G72090 | Bradi1g54570 | Medtr1g090690 |
| 300 | ortho-1487 | AT5G03940 | Bradi1g54570 | Medtr1g090690 |
| 301 | ortho-1489 | AT1G07130 | Bradi1g61610 | NA |
| 302 | ortho-1490 | NA | Bradi3g22960 | Medtr4g025130 |
| 303 | ortho-1493 | AT5G21060 | Bradi4g43920 | NA |
| 304 | ortho-1494 | AT3G48540 | Bradi2g50870 | NA |
| 305 | ortho-1496 | AT4G35987 | Bradi1g68320 | Medtr8g075480 |
| 306 | ortho-1497 | AT5G48340 | Bradi2g57597 | NA |
| 307 | ortho-1498 | AT2G14830 | Bradi4g06630 | NA |
| 308 | ortho-1499 | AT1G80770 | Bradi3g16440 | NA |
| 309 | ortho-1502 | AT5G10690 | Bradi4g30897 | Medtr1g009650 |
| 310 | ortho-1503 | AT3G11620 | Bradi2g61350 | Medtr7g105800 |
| 311 | ortho-1503 | AT3G11620 | Bradi2g61360 | Medtr7g105800 |
| 312 | ortho-1504 | AT4G38370 | Bradi5g03697 | NA |
| 313 | ortho-1505 | AT5G64250 | NA | NA |
| 314 | ortho-1506 | AT1G07040 | Bradi3g28120 | Medtr5g038460 |
| 315 | ortho-1508 | AT3G24730 | Bradi4g20250 | NA |
| 316 | ortho-1509 | AT1G10310 | Bradi3g45920 | Medtr4g070490 |
| 317 | ortho-1511 | AT1G79915 | Bradi1g30350 | NA |
| 318 | ortho-1512 | AT1G24095 | NA | Medtr5g035310 |
| 319 | ortho-1515 | AT4G14480 | Bradi3g38300 | NA |
| 320 | ortho-1518 | AT5G51200 | Bradi3g07490 | Medtr7g099880 |
| 321 | ortho-1519 | AT1G70570 | NA | Medtr8g069525 |
| 322 | ortho-1520 | AT2G42900 | Bradi2g15290 | NA |
| 323 | ortho-1521 | AT5G17570 | Bradi5g19070 | NA |
| 324 | ortho-1523 | AT5G53770 | Bradi2g46260 | NA |
| 325 | ortho-1526 | AT1G36320 | Bradi2g40070 | NA |

| | | | | |
|---|---|---|---|---|
| 326 | **ortho-1533** | AT5G48840 | Bradi1g01460 | Medtr7g058920 |
| 327 | **ortho-1534** | AT3G59490 | Bradi4g35162 | Medtr2g098960 |
| 328 | **ortho-1535** | AT5G38890 | Bradi4g37550 | NA |
| 329 | **ortho-1537** | AT1G19130 | Bradi3g37590 | Medtr4g125100 |
| 330 | **ortho-1540** | AT3G09210 | Bradi1g03400 | NA |
| 331 | **ortho-1542** | AT5G53580 | Bradi3g30847 | Medtr3g092140 |
| 332 | **ortho-163** | AT1G26550 | Bradi4g30050 | Medtr3g037570 |
| 333 | **ortho-164** | AT1G50480 | Bradi4g31680 | Medtr5g059390 |
| 334 | **ortho-165** | AT1G50480 | Bradi4g31680 | Medtr1g116140 |
| 335 | **ortho-165** | AT1G50480 | Bradi4g31680 | Medtr5g059390 |
| 336 | **ortho-166** | AT1G77670 | NA | NA |
| 337 | **ortho-230** | AT2G29630 | Bradi1g12550 | Medtr3g462950 |
| 338 | **ortho-248** | NA | Bradi1g19650 | Medtr5g022000 |
| 339 | **ortho-256** | NA | Bradi1g23600 | NA |
| 340 | **ortho-260** | AT3G43300 | NA | Medtr4g124430 |
| 341 | **ortho-262** | AT5G54770 | Bradi1g25860 | Medtr1g105480 |
| 342 | **ortho-283** | AT3G45100 | Bradi1g32610 | Medtr2g095220 |
| 343 | **ortho-311** | AT2G22125 | Bradi1g45400 | Medtr3g087800 |
| 344 | **ortho-313** | AT2G22125 | Bradi1g45400 | NA |
| 345 | **ortho-330** | NA | Bradi1g50050 | NA |
| 346 | **ortho-336** | NA | NA | NA |
| 347 | **ortho-341** | AT5G08530 | NA | Medtr1g042300 |
| 348 | **ortho-350** | NA | Bradi1g60190 | NA |
| 349 | **ortho-354** | NA | Bradi1g64190 | Medtr1g086640 |
| 350 | **ortho-357** | AT2G47330 | NA | Medtr7g109720 |
| 351 | **ortho-371** | NA | NA | Medtr8g107400 |
| 352 | **ortho-382** | NA | Bradi1g71800 | NA |
| 353 | **ortho-401** | NA | Bradi2g04847 | NA |
| 354 | **ortho-402** | NA | NA | NA |
| 355 | **ortho-459** | NA | NA | NA |
| 356 | **ortho-460** | NA | NA | NA |
| 357 | **ortho-473** | NA | NA | Medtr8g020210 |
| 358 | **ortho-49** | AT2G18710 | Bradi3g19100 | Medtr2g460600 |
| 359 | **ortho-492** | NA | Bradi2g47090 | Medtr8g098675 |
| 360 | **ortho-524** | NA | NA | NA |
| 361 | **ortho-540** | NA | Bradi5g05077 | Medtr4g084660 |
| 362 | **ortho-541** | NA | NA | Medtr1g019500 |
| 363 | **ortho-542** | AT2G05170 | Bradi5g07790 | Medtr1g019500 |
| 364 | **ortho-550** | AT4G04930 | Bradi5g16510 | NA |
| 365 | **ortho-570** | AT5G23110 | Bradi5g26017 | Medtr1g037310 |
| 366 | **ortho-67** | AT4G16340 | Bradi5g24268 | Medtr8g056900 |
| 367 | **uce-11001000** | AT1G22850 | Bradi2g57150 | Medtr4g032670 |
| 368 | **uce-11001086** | AT1G08030 | Bradi1g48757 | Medtr4g058890 |
| 369 | **uce-11001091** | AT1G07970 | Bradi1g65650 | Medtr4g060500 |
| 370 | **uce-11001193** | NA | Bradi1g62210 | Medtr4g076210 |
| 371 | **uce-11001198** | AT5G04590 | Bradi2g20550 | Medtr4g077190 |
| 372 | **uce-11001264** | AT2G35110 | Bradi3g42020 | Medtr4g084140 |
| 373 | **uce-11001281** | AT3G59770 | Bradi2g27230 | Medtr4g087530 |
| 374 | **uce-11001295** | AT1G65070 | Bradi5g26420 | Medtr4g088920 |
| 375 | **uce-11001333** | AT5G37830 | NA | Medtr4g093870 |
| 376 | **uce-11001382** | AT2G21070 | Bradi3g01970 | NA |
| 377 | **uce-11001394** | NA | NA | Medtr4g094860 |
| 378 | **uce-11001396** | AT5G67170 | Bradi5g25650 | Medtr4g094860 |
| 379 | **uce-11001406** | NA | Bradi2g50850 | Medtr4g095660 |
| 380 | **uce-11001423** | AT4G17100 | Bradi5g17235 | Medtr4g099100 |
| 381 | **uce-11001423** | AT4G17100 | Bradi5g17240 | Medtr4g099100 |
| 382 | **uce-11001444** | AT4G16570 | Bradi1g52270 | Medtr4g100980 |
| 383 | **uce-11001445** | AT4G16510 | Bradi3g50910 | Medtr4g101100 |
| 384 | **uce-1100148** | AT5G56290 | Bradi3g39510 | Medtr3g009230 |
| 385 | **uce-11001491** | AT5G62760 | Bradi3g08040 | Medtr4g107890 |
| 386 | **uce-11001498** | AT1G74580 | Bradi1g59781 | Medtr4g108650 |
| 387 | **uce-11001503** | NA | Bradi1g22590 | Medtr4g109230 |
| 388 | **uce-11001520** | AT3G18524 | Bradi1g15260 | Medtr4g111945 |
| 389 | **uce-11001525** | AT2G26890 | Bradi3g34450 | Medtr4g112045 |
| 390 | **uce-11001529** | AT2G26900 | Bradi2g45100 | Medtr4g113090 |
| 391 | **uce-11001573** | AT4G15840 | Bradi2g59770 | Medtr4g119960 |
| 392 | **uce-11001576** | NA | Bradi2g31140 | Medtr4g120340 |
| 393 | **uce-11001586** | AT3G06510 | Bradi4g09920 | Medtr4g122980 |
| 394 | **uce-1100162** | AT3G07530 | Bradi2g58200 | Medtr3g019010 |
| 395 | **uce-11001675** | NA | NA | Medtr4g131750 |
| 396 | **uce-11001677** | AT1G61850 | Bradi1g26060 | Medtr4g131850 |
| 397 | **uce-11001710** | AT4G32910 | Bradi2g49627 | Medtr1g006690 |
| 398 | **uce-11001712** | AT4G32910 | NA | Medtr1g006690 |
| 399 | **uce-11001731** | NA | NA | Medtr1g009980 |
| 400 | **uce-11001739** | AT4G32620 | Bradi2g05030 | Medtr1g010290 |
| 401 | **uce-11001780** | AT5G26040 | Bradi1g37510 | Medtr1g016440 |
| 402 | **uce-11001869** | AT3G10690 | Bradi1g04210 | Medtr1g031690 |
| 403 | **uce-11001871** | AT1G23230 | Bradi3g56910 | Medtr1g031910 |
| 404 | **uce-11001922** | NA | Bradi1g42030 | Medtr1g040145 |
| 405 | **uce-11001923** | AT3G22990 | Bradi3g47466 | Medtr1g041115 |
| 406 | **uce-11001924** | AT3G55760 | Bradi4g15010 | Medtr1g041275 |
| 407 | **uce-11001928** | AT3G23490 | Bradi3g28470 | Medtr1g041795 |
| 408 | **uce-11001946** | AT3G24190 | Bradi3g55317 | Medtr1g046620 |
| 409 | **uce-11001967** | AT3G56160 | Bradi3g44850 | Medtr1g050462 |
| 410 | **uce-11001969** | AT3G56160 | Bradi3g44850 | Medtr1g050462 |
| 411 | **uce-11001970** | NA | NA | Medtr1g050462 |
| 412 | **uce-11001971** | AT1G42990 | Bradi1g35790 | Medtr1g050502 |
| 413 | **uce-11001986** | AT2G41020 | Bradi2g58667 | Medtr1g051155 |
| 414 | **uce-11001993** | AT3G57630 | Bradi1g73000 | Medtr1g052730 |
| 415 | **uce-11002009** | NA | Bradi3g10360 | Medtr1g054625 |
| 416 | **uce-11002062** | AT5G03070 | Bradi2g00320 | Medtr1g066760 |
| 417 | **uce-11002066** | AT5G12130 | Bradi2g37780 | Medtr1g067200 |
| 418 | **uce-11002087** | AT2G20360 | Bradi3g55340 | Medtr1g069455 |
| 419 | **uce-11002090** | AT2G20420 | Bradi3g49070 | Medtr1g069645 |
| 420 | **uce-11002094** | AT2G20495 | NA | Medtr1g069790 |
| 421 | **uce-11002095** | AT2G20495 | Bradi1g29077 | Medtr1g069790 |
| 422 | **uce-11002115** | AT4G25120 | Bradi1g27110 | Medtr1g072800 |
| 423 | **uce-11002119** | AT4G17740 | Bradi4g37555 | Medtr1g073130 |
| 424 | **uce-1100219** | NA | Bradi1g20410 | Medtr3g435140 |
| 425 | **uce-11002249** | AT2G38680 | NA | Medtr1g090380 |
| 426 | **uce-11002256** | NA | NA | Medtr1g090693 |
| 427 | **uce-11002271** | AT2G36360 | Bradi1g02610 | Medtr1g492850 |
| 428 | **uce-11002284** | AT5G17530 | Bradi1g28790 | Medtr1g094980 |
| 429 | **uce-11002285** | NA | Bradi5g15930 | Medtr1g095040 |
| 430 | **uce-11002317** | AT4G09620 | Bradi1g02400 | Medtr1g096340 |
| 431 | **uce-11002334** | AT1G22770 | Bradi2g05226 | Medtr1g098160 |
| 432 | **uce-11002367** | AT4G21800 | Bradi4g36230 | Medtr1g100240 |
| 433 | **uce-11002425** | NA | Bradi4g08260 | Medtr1g107485 |
| 434 | **uce-11002469** | AT2G17020 | Bradi1g76050 | Medtr1g114010 |
| 435 | **uce-11002471** | AT2G17020 | Bradi1g76050 | Medtr1g114010 |
| 436 | **uce-11002544** | NA | NA | Medtr1g116820 |
| 437 | **uce-11002601** | AT2G14050 | Bradi1g45695 | Medtr2g009190 |
| 438 | **uce-11002602** | AT2G14050 | NA | Medtr2g009190 |
| 439 | **uce-11002604** | AT2G14050 | Bradi1g45695 | Medtr2g009190 |
| 440 | **uce-11002619** | AT1G28120 | Bradi3g41530 | Medtr2g011420 |
| 441 | **uce-11002659** | AT2G25950 | Bradi2g41230 | Medtr2g018310 |
| 442 | **uce-11002669** | AT5G46210 | Bradi1g06387 | Medtr2g019260 |
| 443 | **uce-11002693** | AT4G04870 | Bradi2g52187 | Medtr2g022840 |
| 444 | **uce-11002703** | AT5G47790 | Bradi3g20620 | Medtr2g027390 |
| 445 | **uce-11002731** | AT5G49930 | Bradi4g04650 | NA |
| 446 | **uce-11002751** | AT5G49030 | Bradi3g57220 | Medtr2g436760 |
| 447 | **uce-11002757** | AT4G18460 | Bradi2g08490 | Medtr2g036040 |
| 448 | **uce-11002784** | AT4G35740 | Bradi3g57087 | NA |
| 449 | **uce-11002785** | AT4G35740 | Bradi3g57087 | Medtr2g042440 |
| 450 | **uce-11002800** | AT1G34630 | Bradi1g71820 | Medtr2g044600 |
| 451 | **uce-11002801** | AT1G34630 | Bradi1g71820 | Medtr2g044600 |
| 452 | **uce-11002833** | AT3G42660 | Bradi2g18270 | Medtr2g059200 |
| 453 | **uce-11002842** | AT2G18710 | Bradi3g19100 | Medtr2g460600 |
| 454 | **uce-11002844** | AT2G18710 | Bradi3g19100 | Medtr2g460600 |
| 455 | **uce-1100286** | NA | NA | Medtr3g053220 |
| 456 | **uce-11002870** | AT1G54350 | Bradi2g07076 | Medtr2g062790 |
| 457 | **uce-11002913** | AT1G16650 | Bradi4g02070 | Medtr2g064450 |
| 458 | **uce-11002988** | AT5G54590 | NA | Medtr2g082430 |
| 459 | **uce-11003046** | AT5G60600 | Bradi3g48080 | Medtr2g094160 |
| 460 | **uce-11003047** | AT5G60600 | Bradi3g48080 | Medtr2g094160 |
| 461 | **uce-11003063** | AT3G45830 | Bradi2g37740 | Medtr2g096000 |
| 462 | **uce-11003082** | AT3G20780 | Bradi4g27057 | Medtr2g098510 |
| 463 | **uce-11003083** | AT3G20780 | Bradi4g27057 | Medtr2g098510 |
| 464 | **uce-11003101** | AT1G17650 | Bradi2g41630 | Medtr2g099910 |
| 465 | **uce-11003104** | AT1G72990 | Bradi2g18440 | Medtr2g100000 |
| 466 | **uce-11003164** | AT4G02580 | Bradi2g20370 | Medtr2g104110 |
| 467 | **uce-11003238** | AT2G35450 | Bradi5g08360 | Medtr7g022480 |
| 468 | **uce-11003253** | AT3G04020 | Bradi1g66580 | Medtr7g024560 |
| 469 | **uce-11003276** | AT5G14220 | Bradi5g14120 | Medtr7g031310 |
| 470 | **uce-11003279** | AT5G14220 | Bradi5g14120 | Medtr7g031310 |
| 471 | **uce-11003283** | AT2G36810 | Bradi1g75250 | NA |
| 472 | **uce-11003286** | AT3G01660 | NA | Medtr7g032900 |
| 473 | **uce-11003293** | AT3G28030 | Bradi1g70850 | Medtr7g034405 |
| 474 | **uce-11003302** | AT1G08840 | Bradi5g19741 | Medtr7g034575 |
| 475 | **uce-11003322** | NA | Bradi1g25830 | Medtr7g050860 |
| 476 | **uce-11003324** | AT5G15170 | Bradi1g25830 | Medtr7g050860 |
| 477 | **uce-11003333** | AT3G25410 | Bradi3g43450 | Medtr7g053090 |
| 478 | **uce-11003461** | AT5G36230 | Bradi4g19650 | Medtr7g080110 |
| 479 | **uce-11003478** | AT1G59760 | Bradi1g52230 | Medtr7g080730 |
| 480 | **uce-11003510** | AT2G38130 | Bradi2g32930 | Medtr7g084140 |
| 481 | **uce-11003596** | AT5G06680 | Bradi4g31640 | Medtr7g093560 |
| 482 | **uce-1100362** | AT5G64250 | Bradi3g38730 | Medtr3g069190 |
| 483 | **uce-11003641** | AT5G51200 | Bradi3g07490 | Medtr7g099880 |
| 484 | **uce-11003643** | AT5G51200 | Bradi3g07490 | Medtr7g099880 |
| 485 | **uce-11003655** | AT2G37240 | Bradi1g65880 | Medtr7g100540 |
| 486 | **uce-11003675** | AT2G36810 | Bradi1g75250 | Medtr7g102430 |
| 487 | **uce-11003678** | AT2G36810 | NA | Medtr7g102430 |
| 488 | **uce-11003723** | AT5G06550 | Bradi4g16020 | Medtr7g106990 |
| 489 | **uce-11003764** | AT4G01860 | NA | Medtr7g110260 |
| 490 | **uce-11003811** | AT3G04480 | Bradi3g54847 | Medtr7g111650 |
| 491 | **uce-11003827** | AT3G05510 | Bradi2g59710 | Medtr7g112680 |
| 492 | **uce-11003848** | AT3G04650 | Bradi1g23880 | Medtr7g115310 |
| 493 | **uce-11003905** | AT2G04560 | Bradi2g50040 | Medtr8g012965 |

| | | | | |
|---|---|---|---|---|
| **494** | **uce-11003939** | NA | Bradi2g41580 | Medtr8g020210 |
| **495** | **uce-11003944** | AT1G17410 | Bradi3g46220 | Medtr8g021233 |
| **496** | **uce-11003969** | AT5G12370 | Bradi4g39550 | Medtr8g023330 |
| **497** | **uce-11004007** | AT1G01770 | Bradi1g28600 | Medtr8g031480 |
| **498** | **uce-11004045** | NA | Bradi2g57967 | Medtr8g037325 |
| **499** | **uce-11004050** | AT3G54440 | Bradi2g61010 | Medtr8g039160 |
| **500** | **uce-11004054** | AT2G38730 | Bradi1g04250 | Medtr8g040180 |
| **501** | **uce-11004055** | AT5G06970 | Bradi1g20630 | Medtr8g040190 |
| **502** | **uce-11004063** | NA | Bradi2g21790 | Medtr8g043970 |
| **503** | **uce-11004085** | AT1G78630 | Bradi2g49850 | Medtr8g054450 |
| **504** | **uce-1100411** | AT1G77930 | Bradi1g20730 | Medtr3g073560 |
| **505** | **uce-11004158** | AT1G70570 | Bradi3g02900 | Medtr8g069525 |
| **506** | **uce-11004173** | AT3G54230 | Bradi2g43127 | Medtr8g070990 |
| **507** | **uce-11004182** | AT5G48340 | Bradi2g57597 | Medtr8g071270 |
| **508** | **uce-11004203** | AT5G11980 | Bradi4g05710 | Medtr8g074590 |
| **509** | **uce-11004290** | AT1G55350 | Bradi3g53020 | Medtr8g088520 |
| **510** | **uce-11004293** | AT1G55350 | Bradi3g53020 | Medtr8g088520 |
| **511** | **uce-11004308** | AT3G26085 | Bradi3g04310 | Medtr8g089550 |
| **512** | **uce-11004373** | AT1G63660 | Bradi3g20590 | Medtr8g098325 |
| **513** | **uce-11004395** | AT2G07360 | Bradi1g67140 | Medtr8g100105 |
| **514** | **uce-11004403** | AT4G12750 | Bradi4g03303 | NA |
| **515** | **uce-11004422** | AT1G49040 | Bradi2g41690 | Medtr8g104010 |
| **516** | **uce-11004430** | AT1G14850 | Bradi1g69510 | Medtr8g105290 |
| **517** | **uce-11004457** | AT1G76630 | Bradi3g32780 | Medtr5g004660 |
| **518** | **uce-11004459** | AT1G76630 | Bradi3g32780 | Medtr5g004660 |
| **519** | **uce-11004463** | AT3G19210 | Bradi3g58092 | Medtr5g004720 |
| **520** | **uce-11004465** | AT3G19210 | Bradi3g58092 | Medtr5g004720 |
| **521** | **uce-11004479** | AT5G44750 | NA | Medtr5g004960 |
| **522** | **uce-11004488** | NA | Bradi5g20597 | Medtr5g005440 |
| **523** | **uce-1100450** | AT5G17990 | Bradi1g76800 | Medtr3g077590 |
| **524** | **uce-11004502** | AT5G42390 | Bradi1g35670 | Medtr5g007380 |
| **525** | **uce-11004521** | AT1G71500 | Bradi4g21270 | Medtr5g008750 |
| **526** | **uce-11004546** | AT1G12250 | Bradi3g50060 | Medtr5g012110 |
| **527** | **uce-11004564** | AT5G41150 | Bradi1g78620 | Medtr5g013480 |
| **528** | **uce-11004565** | AT5G41150 | Bradi1g78620 | Medtr5g013480 |
| **529** | **uce-11004567** | AT4G17140 | Bradi3g33740 | Medtr5g013780 |
| **530** | **uce-11004573** | AT5G47010 | Bradi1g27027 | Medtr5g013820 |
| **531** | **uce-11004575** | AT5G47010 | Bradi1g27027 | Medtr5g013820 |
| **532** | **uce-11004576** | AT5G47010 | Bradi1g27027 | Medtr5g013820 |
| **533** | **uce-11004579** | AT5G46630 | Bradi3g52130 | Medtr5g014330 |
| **534** | **uce-11004594** | NA | Bradi2g57476 | Medtr5g015680 |
| **535** | **uce-11004604** | AT2G42240 | NA | Medtr5g016680 |
| **536** | **uce-11004606** | AT4G36980 | NA | Medtr5g017430 |
| **537** | **uce-11004609** | NA | NA | Medtr5g017430 |
| **538** | **uce-11004631** | AT5G66530 | Bradi4g28160 | Medtr5g020640 |
| **539** | **uce-11004631** | AT5G66530 | Bradi4g28150 | Medtr5g020640 |
| **540** | **uce-11004678** | AT1G31780 | Bradi1g69350 | Medtr5g025410 |
| **541** | **uce-11004685** | AT1G31360 | NA | Medtr5g026590 |
| **542** | **uce-11004689** | AT5G45380 | Bradi3g21640 | Medtr5g026640 |
| **543** | **uce-11004704** | AT3G07300 | Bradi3g23230 | Medtr5g031790 |
| **544** | **uce-11004705** | AT2G22530 | Bradi3g60040 | Medtr5g032210 |
| **545** | **uce-11004715** | AT2G22530 | Bradi3g60040 | Medtr5g032210 |
| **546** | **uce-11004733** | AT1G70070 | Bradi3g59790 | Medtr5g036660 |
| **547** | **uce-11004736** | NA | NA | Medtr5g037050 |
| **548** | **uce-11004754** | AT1G10520 | Bradi1g44820 | Medtr7g039450 |
| **549** | **uce-11004766** | AT1G26640 | Bradi2g50820 | Medtr5g042980 |
| **550** | **uce-11004854** | AT4G02990 | Bradi2g25830 | Medtr5g068860 |
| **551** | **uce-11004863** | AT4G27590 | NA | Medtr5g069205 |
| **552** | **uce-11004960** | AT1G06560 | Bradi3g21700 | Medtr5g091110 |
| **553** | **uce-11004963** | AT1G06560 | Bradi3g21700 | Medtr5g091110 |
| **554** | **uce-11004974** | AT4G26980 | Bradi1g32747 | Medtr5g092970 |
| **555** | **uce-11005008** | AT1G80680 | Bradi1g73110 | Medtr5g097890 |
| **556** | **uce-11005009** | AT1G80680 | Bradi1g73110 | Medtr5g097890 |
| **557** | **uce-11005026** | AT1G66520 | NA | Medtr6g003980 |
| **558** | **uce-1100503** | AT2G04740 | Bradi1g42730 | Medtr3g082870 |
| **559** | **uce-11005074** | AT5G39940 | NA | Medtr6g011270 |
| **560** | **uce-11005123** | AT5G15680 | Bradi3g60610 | Medtr6g016215 |
| **561** | **uce-11005138** | AT5G38460 | Bradi1g63540 | Medtr6g017315 |
| **562** | **uce-11005152** | AT5G14260 | Bradi4g39580 | Medtr6g023320 |
| **563** | **uce-11005158** | AT1G17760 | Bradi4g03980 | Medtr6g026900 |
| **564** | **uce-11005168** | AT2G33260 | Bradi1g53780 | Medtr6g032995 |
| **565** | **uce-11005228** | AT1G73350 | Bradi4g10720 | Medtr6g068990 |
| **566** | **uce-11005234** | NA | Bradi3g59890 | Medtr6g071210 |
| **567** | **uce-11005240** | AT3G17750 | Bradi1g10530 | Medtr6g074905 |
| **568** | **uce-11005253** | NA | Bradi3g01527 | Medtr6g081020 |
| **569** | **uce-1100526** | NA | Bradi2g42017 | Medtr3g083970 |
| **570** | **uce-11005260** | AT1G34130 | Bradi5g26030 | Medtr6g077750 |
| **571** | **uce-11005266** | AT1G18030 | Bradi4g27880 | Medtr6g081850 |
| **572** | **uce-1100542** | AT4G30990 | Bradi2g42017 | Medtr3g083970 |
| **573** | **uce-1100560** | AT2G32000 | Bradi4g34290 | Medtr3g084800 |
| **574** | **uce-1100563** | AT2G32000 | Bradi4g34290 | Medtr3g084800 |
| **575** | **uce-1100565** | AT2G32000 | Bradi4g34290 | Medtr3g084800 |
| **576** | **uce-1100572** | AT1G24310 | Bradi1g52600 | Medtr3g085590 |
| **577** | **uce-1100615** | NA | NA | NA |
| **578** | **uce-1100626** | AT5G53580 | Bradi3g30847 | Medtr3g092140 |
| **579** | **uce-1100644** | NA | Bradi1g20790 | Medtr3g094840 |
| **580** | **uce-1100658** | AT5G63920 | Bradi1g74120 | Medtr3g096640 |
| **581** | **uce-1100693** | AT5G10920 | Bradi1g64597 | Medtr3g100220 |
| **582** | **uce-1100696** | AT5G49970 | Bradi3g23340 | Medtr3g100890 |
| **583** | **uce-1100712** | AT1G10130 | Bradi1g09810 | Medtr3g103270 |
| **584** | **uce-1100733** | AT2G45700 | Bradi4g31227 | Medtr3g105470 |
| **585** | **uce-1100733** | AT2G45700 | Bradi4g31220 | Medtr3g105470 |
| **586** | **uce-1100740** | AT4G31770 | Bradi1g10190 | Medtr3g105810 |
| **587** | **uce-1100752** | AT2G25320 | Bradi3g48647 | Medtr3g106700 |
| **588** | **uce-1100755** | NA | NA | Medtr3g107060 |
| **589** | **uce-1100757** | AT4G32320 | Bradi3g40330 | Medtr3g107060 |
| **590** | **uce-1100789** | AT4G31600 | Bradi1g46960 | Medtr3g109070 |
| **591** | **uce-1100804** | AT2G25840 | Bradi2g49520 | Medtr3g110360 |
| **592** | **uce-1100842** | AT1G42430 | Bradi1g76190 | Medtr3g115210 |
| **593** | **uce-1100843** | AT1G63610 | Bradi3g60850 | Medtr3g115240 |
| **594** | **uce-1100845** | NA | Bradi1g11317 | Medtr3g116250 |
| **595** | **uce-1100848** | AT1G49340 | Bradi1g11317 | Medtr3g116250 |
| **596** | **uce-1100849** | AT1G49340 | Bradi1g11317 | Medtr3g116250 |
| **597** | **uce-1100856** | AT4G34730 | Bradi1g51470 | Medtr3g117600 |
| **598** | **uce-1100889** | AT1G16080 | Bradi3g19310 | Medtr4g007910 |
| **599** | **uce-1100959** | AT5G02250 | Bradi2g37760 | Medtr4g021750 |
| **600** | **uce-1100965** | AT1G48050 | Bradi1g01007 | Medtr4g023560 |
| **601** | **uce-1100971** | AT2G45270 | Bradi2g11360 | Medtr4g024920 |
| **602** | **uce-1100982** | NA | Bradi5g16830 | Medtr4g026500 |
| **603** | **uce-1100982** | NA | Bradi5g16840 | Medtr4g026500 |
| **604** | **uce-1100998** | AT1G33810 | Bradi3g13560 | Medtr4g031650 |
| **605** | **uce-12001086** | AT3G16200 | Bradi2g13630 | Medtr2g101480 |
| **606** | **uce-12001183** | AT5G23630 | Bradi2g25857 | Medtr3g097560 |
| **607** | **uce-12001288** | AT4G33060 | Bradi2g41910 | Medtr2g023120 |
| **608** | **uce-1200141** | NA | Bradi5g23940 | NA |
| **609** | **uce-12001449** | AT1G27340 | Bradi2g59200 | Medtr3g085160 |
| **610** | **uce-12001605** | AT5G17170 | Bradi3g16187 | Medtr5g008050 |
| **611** | **uce-12001743** | AT4G34030 | Bradi3g36140 | Medtr4g085890 |
| **612** | **uce-12001936** | AT3G12040 | Bradi3g57425 | Medtr7g093540 |
| **613** | **uce-1200199** | AT2G36360 | Bradi1g02610 | Medtr1g492850 |
| **614** | **uce-12002031** | AT4G32700 | Bradi4g08150 | Medtr1g008600 |
| **615** | **uce-12002186** | AT1G26550 | Bradi4g30050 | Medtr3g037570 |
| **616** | **uce-12002218** | AT2G32000 | Bradi4g34290 | Medtr3g084800 |
| **617** | **uce-12002225** | AT1G05910 | Bradi4g35150 | Medtr4g067100 |
| **618** | **uce-1200272** | AT1G75200 | Bradi1g07101 | Medtr1g022235 |
| **619** | **uce-1200293** | NA | Bradi1g08917 | Medtr1g071260 |
| **620** | **uce-1200294** | NA | Bradi1g08917 | Medtr1g071260 |
| **621** | **uce-1200301** | AT1G10130 | Bradi1g09810 | Medtr3g103270 |
| **622** | **uce-1200306** | NA | Bradi1g10447 | NA |
| **623** | **uce-1200413** | AT1G67500 | Bradi1g18510 | Medtr7g113380 |
| **624** | **uce-1200581** | NA | Bradi1g36340 | Medtr4g116430 |
| **625** | **uce-1200660** | AT1G16780 | Bradi1g46330 | Medtr4g005280 |
| **626** | **uce-120069** | AT1G58050 | Bradi5g10330 | Medtr4g134790 |
| **627** | **uce-1200740** | AT5G08530 | Bradi1g53800 | Medtr1g042300 |
| **628** | **uce-1200772** | AT3G03710 | Bradi1g56280 | Medtr7g086640 |
| **629** | **uce-1200935** | AT5G18410 | Bradi1g75470 | Medtr7g116590 |
| **630** | **uce-120094** | AT1G05055 | Bradi5g15270 | Medtr7g089440 |
| **631** | **uce-1200979** | AT4G04350 | Bradi2g01504 | Medtr4g132500 |
| **632** | **uce-13001080** | AT4G11380 | Bradi1g62210 | Medtr4g076210 |
| **633** | **uce-13001080** | AT4G23460 | Bradi1g62210 | Medtr4g076210 |
| **634** | **uce-13001153** | AT4G33520 | Bradi3g38790 | Medtr4g094232 |
| **635** | **uce-1300245** | AT5G61140 | Bradi1g70057 | Medtr3g025300 |
| **636** | **uce-1300502** | AT1G35660 | Bradi2g56490 | Medtr1g050670 |

**Table 3** Captured loci mapping to the same gene models or to more than one gene model in the reference genomes of three model species

| Locus | *A. thaliana* | *B. distachyon* | *M. truncatula* |
|---|---|---|---|
| **ortho-1219** | AT5G02710 | Bradi1g61764 | Medtr0016s0020/Medtr1g067330 |
| **uce-11001710/uce-11001712** | AT4G32910 | Bradi2g49627 | Medtr1g006690 |
| **ortho-1290/uce-11001780** | AT5G26040 | Bradi1g37510 | Medtr1g016440 |
| **ortho-541/ortho-542** | AT2G05170 | Bradi5g07790 | Medtr1g019500 |
| **ortho-1343/uce-11001922** | AT5G17560 | Bradi1g42030 | Medtr1g040145 |
| **ortho-1236/uce-11001928** | AT3G23490 | Bradi3g28470 | Medtr1g041795 |
| **ortho-341/uce-1200740** | AT5G08530 | Bradi1g53800 | Medtr1g042300 |
| **uce-11001967/uce-11001969/uce-11001970** | AT3G56160 | Bradi3g44850 | Medtr1g050462 |
| **ortho-1022/uce-11002009** | AT3G58530 | Bradi3g10360 | Medtr1g054625 |
| **ortho-1046/uce-11002062** | AT5G03070 | Bradi2g00320 | Medtr1g066760 |
| **ortho-1468/uce-11002066** | AT5G12130 | Bradi2g37780 | Medtr1g067200 |
| **ortho-1055/uce-11002087** | AT2G20360 | Bradi3g55340 | Medtr1g069455 |
| **uce-11002094/uce-11002095** | AT2G20495 | Bradi1g29077 | Medtr1g069790 |
| **uce-1200293/uce-1200294** | NA | Bradi1g08917 | Medtr1g071260 |
| **ortho-1297/uce-11002249** | AT2G38680 | Bradi1g13590 | Medtr1g090380 |
| **ortho-1487** | AT1G72090/AT5G03940 | Bradi1g54570 | Medtr1g090690 |
| **ortho-1400/uce-11002256** | AT5G04360 | Bradi5g00540 | Medtr1g090693 |
| **ortho-1360/uce-11002317** | AT4G09620 | Bradi1g02400 | Medtr1g096340 |
| **uce-11002469/uce-11002471** | AT2G17020 | Bradi1g76050 | Medtr1g114010 |
| **uce-11002271/uce-1200199** | AT2G36360 | Bradi1g02610 | Medtr1g492850 |
| **uce-11002601/uce-11002604/uce-11002602** | AT2G14050 | Bradi1g45695 | Medtr2g009190 |
| **ortho-1045/uce-11002693** | AT4G04870 | Bradi2g52187 | Medtr2g022840 |
| **ortho-1124/uce-11002757** | AT4G18460 | Bradi2g08490 | Medtr2g036040 |
| **uce-11002785/uce-11002784** | AT4G35740 | Bradi3g57087 | Medtr2g042440 |
| **uce-11002800/uce-11002801** | AT1G34630 | Bradi1g71820 | Medtr2g044600 |
| **ortho-1475** | AT3G24200 | Bradi1g02380 | Medtr2g060550/Medtr7g051960 |
| **ortho-1375/uce-11002913** | AT1G16650 | Bradi4g02070 | Medtr2g064450 |
| **ortho-1462** | AT1G28100/AT1G28110 | NA | Medtr2g072940 |
| **uce-11003046/uce-11003047** | AT5G60600 | Bradi3g48080 | Medtr2g094160 |
| **uce-11003082/uce-11003083** | AT3G20780 | Bradi4g27057 | Medtr2g098510 |
| **ortho-1011/uce-11002751** | AT5G49030 | Bradi3g57220 | Medtr2g436760 |
| **ortho-1148/ortho-49/uce-11002842/uce-11002844** | AT2G18710 | Bradi3g19100 | Medtr2g460600 |
| **ortho-1146** | AT5G07960 | Bradi1g08201/Bradi1g08210 | Medtr3g024170/Medtr3g024190 |
| **ortho-1341** | AT3G52030/AT3G52040 | Bradi4g23700 | Medtr3g034210 |
| **ortho-163/uce-12002186** | AT1G26550 | Bradi4g30050 | Medtr3g037570 |
| **ortho-1110/uce-1100286** | AT3G20440 | Bradi1g41970 | Medtr3g053220 |
| **ortho-1505/uce-1100362** | AT5G64250 | Bradi3g38730 | Medtr3g069190 |
| **ortho-1455/uce-1100411** | AT1G77930 | Bradi1g20730 | Medtr3g073560 |
| **ortho-1281/uce-1100503** | AT2G04740 | Bradi1g42730 | Medtr3g082870 |
| **ortho-1425/uce-1100542/uce-1100526** | AT3G25430/AT4G30990 | Bradi2g42017 | Medtr3g083970 |
| **uce-1100560/uce-1100563/uce-1100565/uce-12002218** | AT2G32000 | Bradi4g34290 | Medtr3g084800 |
| **ortho-311/ortho-313** | AT2G22125 | Bradi1g45400 | Medtr3g087800 |
| **ortho-1542/uce-1100626** | AT5G53580 | Bradi3g30847 | Medtr3g092140 |
| **uce-1100712/uce-1200301** | AT1G10130 | Bradi1g09810 | Medtr3g103270 |
| **uce-1100733** | AT2G45700 | Bradi4g31220/Bradi4g31227 | Medtr3g105470 |
| **uce-1100757/uce-1100755** | AT4G32320 | Bradi3g40330 | Medtr3g107060 |
| **ortho-1274/uce-1100789** | AT4G31600 | Bradi1g46960 | Medtr3g109070 |
| **uce-1100848/uce-1100849/uce-1100845** | AT1G49340 | Bradi1g11317 | Medtr3g116250 |
| **ortho-1064/uce-1100959** | AT5G02250 | Bradi2g37760 | Medtr4g021750 |
| **ortho-1480/uce-1100971** | AT2G45270 | Bradi2g11360 | Medtr4g024920 |
| **uce-1100982** | NA | Bradi5g16830/Bradi5g16840 | Medtr4g026500 |
| **ortho-1032/uce-1100998** | AT1G33810 | Bradi3g13560 | Medtr4g031650 |
| **uce-13001080/uce-11001193** | AT4G11380/AT4G23460 | Bradi1g62210 | Medtr4g076210 |
| **ortho-1222/uce-11001396/uce-11001394** | AT5G67170 | Bradi5g25650 | Medtr4g094860 |
| **uce-11001423** | AT4G17100 | Bradi5g17235/Bradi5g17240 | Medtr4g099100 |
| **ortho-1364/uce-11001444** | AT4G16570 | Bradi1g52270 | Medtr4g100980 |
| **ortho-1053/uce-11001445** | AT4G16510 | Bradi3g50910 | Medtr4g101100 |
| **ortho-1348** | AT5G50290 | Bradi3g39950/Bradi3g39960 | Medtr4g105550 |
| **ortho-1476/uce-11001520** | AT3G18524 | Bradi1g15260 | Medtr4g111945 |
| **uce-11004457/uce-11004459** | AT1G76630 | Bradi3g32780 | Medtr5g004660 |
| **uce-11004463/uce-11004465** | AT3G19210 | Bradi3g58092 | Medtr5g004720 |
| **ortho-1115/uce-11004521** | AT1G71500 | Bradi4g21270 | Medtr5g008750 |
| **ortho-1069** | AT1G50490/AT3G20060 | Bradi2g10220 | Medtr5g009020 |
| **ortho-1440/uce-11004546** | AT1G12250 | Bradi3g50060 | Medtr5g012110 |
| **uce-11004564/uce-11004565** | AT5G41150 | Bradi1g78620 | Medtr5g013480 |
| **uce-11004573/uce-11004575/uce-11004576** | AT5G47010 | Bradi1g27027 | Medtr5g013820 |
| **uce-11004606/uce-11004609** | AT4G36980 | NA | Medtr5g017430 |
| **uce-11004631** | AT5G66530 | Bradi4g28150/Bradi4g28160 | Medtr5g020640 |
| **ortho-1278/uce-11004689** | AT5G45380 | Bradi3g21640 | Medtr5g026640 |
| **ortho-1289/uce-11004705/uce-11004715** | AT2G22530 | Bradi3g60040 | Medtr5g032210 |
| **ortho-1242/uce-11004736** | AT1G59840 | Bradi5g24257 | Medtr5g037050 |
| **ortho-164/ortho-165** | AT1G50480 | Bradi4g31680 | Medtr5g059390/Medtr1g116140 |
| **ortho-1402/uce-11004854** | AT4G02990 | Bradi2g25830 | Medtr5g068860 |
| **ortho-1105** | AT5G17460 | Bradi2g01810 | Medtr5g086010/Medtr5g087440 |
| **ortho-1315/uce-11004960/uce-11004963** | AT1G06560 | Bradi3g21700 | Medtr5g091110 |
| **uce-11005008/uce-11005009** | AT1G80680 | Bradi1g73110 | Medtr5g097890 |
| **ortho-1106/uce-11005026** | AT1G66520/AT1G66510 | NA | Medtr6g003980 |

| | | | |
|---|---|---|---|
| **ortho-1198/uce-11005074** | AT5G39940 | Bradi4g00497 | Medtr6g011270 |
| **ortho-1288/uce-11005138** | AT5G38460 | Bradi1g63540 | Medtr6g017315 |
| **ortho-1112/uce-11005266** | AT1G18030 | Bradi4g27880 | Medtr6g081850 |
| **ortho-1122/uce-11003276/uce-11003279** | AT5G14220 | Bradi5g14120 | Medtr7g031310 |
| **ortho-1120/uce-11003302** | AT1G08840 | Bradi5g19741 | Medtr7g034575 |
| **ortho-1019/uce-11004754** | AT1G10520 | Bradi1g44820 | Medtr7g039450/Medtr5g040170 |
| **ortho-1212/uce-11003324/uce-11003322** | AT5G15170 | Bradi1g25830 | Medtr7g050860 |
| **ortho-1186/uce-11003333** | AT3G25410 | Bradi3g43450 | Medtr7g053090 |
| **ortho-1200/uce-11003510** | AT2G38130 | Bradi2g32930 | Medtr7g084140 |
| **ortho-1084/uce-12001936** | AT3G12040 | Bradi3g57425 | Medtr7g093540 |
| **ortho-1518/uce-11003641/uce-11003643** | AT5G51200 | Bradi3g07490 | Medtr7g099880 |
| **ortho-1007/uce-11003655** | AT2G37240 | Bradi1g65880 | Medtr7g100540 |
| **uce-11003675/uce-11003283/uce-11003678** | AT2G36810 | Bradi1g75250 | Medtr7g102430 |
| **ortho-1503** | AT3G11620 | Bradi2g61350/Bradi2g61360 | Medtr7g105800 |
| **ortho-1361** | AT4G03140/AT4G03150 | NA | Medtr7g107680 |
| **ortho-1263/uce-11003764** | AT4G01860 | Bradi1g77780 | Medtr7g110260 |
| **ortho-1342/uce-11003811** | AT3G04480 | Bradi3g54847 | Medtr7g111650 |
| **ortho-1052/uce-11003848** | AT3G04650 | Bradi1g23880 | Medtr7g115310 |
| **ortho-473/uce-11003939** | NA | Bradi2g41580 | Medtr8g020210 |
| **ortho-1401/uce-11003944** | AT1G17410 | Bradi3g46220 | Medtr8g021233 |
| **ortho-1519/uce-11004158** | AT1G70570 | Bradi3g02900 | Medtr8g069525 |
| **ortho-1497/uce-11004182** | AT5G48340 | Bradi2g57597 | Medtr8g071270 |
| **uce-11004290/uce-11004293** | AT1G55350 | Bradi3g53020 | Medtr8g088520 |
| **ortho-1322** | AT1G08220 | Bradi2g34156/Bradi2g34159 | NA |
| **ortho-1203** | AT5G44740 | Bradi2g50340/Bradi2g50350 | NA |
| **ortho-1241** | AT2G25530/AT3G15290 | Bradi2g52350 | NA |

Total loci count with shared genes: 85
Total loci mapping to more than one gene in any species: 23
Total loci mapping to more than one gene in *A. thaliana*: 9
Total loci mapping to more than one gene in *B. distachyon*: 9
Total loci mapping to more than one gene in *M. truncatula*: 6

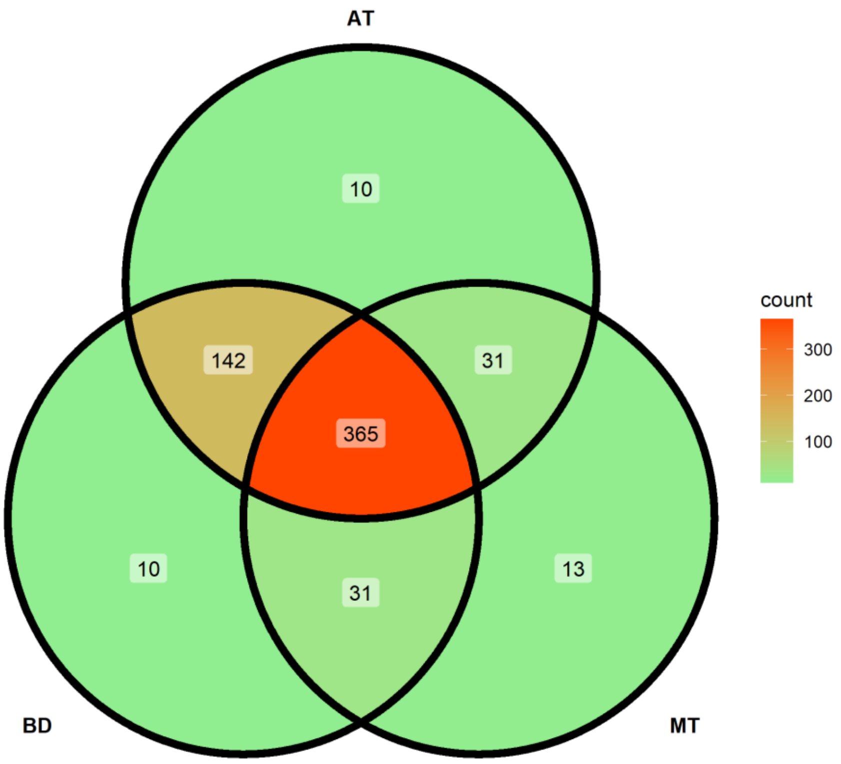


**Figure 1 Venn diagram of the captured loci present in three reference genomes.** Reference genomes: AT: *Arabidopsis thaliana*, BD: *Brachipodium distachyon*, MT: *Medicago truncatula.* Non-mapped loci: 9 (ortho-1003, ortho-143, ortho-145, ortho-336, ortho-402, ortho-459, ortho-460, ortho-524, uce-1100615)

### Validation phase

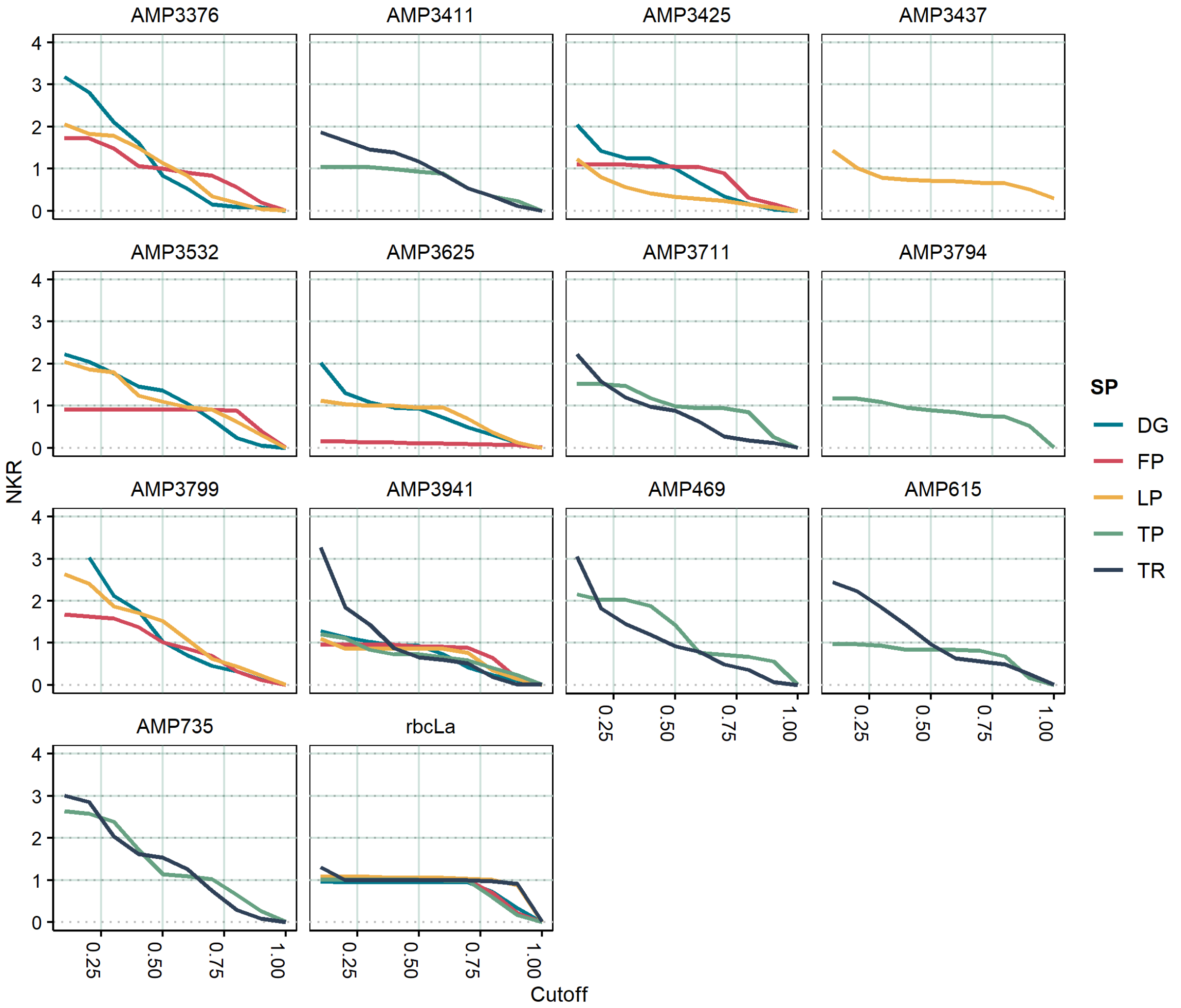


**Figure 2 Effect of *k*-mer filtering threshold on normalized *k*-mer richness (NKR).** Normalized *k*-mer richness is dependent on a cutoff set for *k*-mer frequencies. The *k*-mer frequencies were calculated in proportion to the maximum *k*-mer count per amplicon and species. The *k*-mer frequencies ranged from 5 (*k*-mers with a frequency of 5% the maximum *k*-mer count per amplicon and species) to 1 (only *k*-mers with the maximum *k*-mer count per amplicon and species are considered). Each species-amplicon pair is represented by a total of 16 genotypes, with 100 sub-sampled reads per genotype.

**Table 4** Single-plant material used for amplicon sequencing and SSR analysis

| Index | Sample name | Species | Cultivar |
|---|---|---|---|
| 1 | DG-BRE-1 | *Dactylis glomerata* L. | 'Brennus' (R2n, FR) |
| 2 | DG-BRE-2 | *Dactylis glomerata* L. | 'Brennus' (R2n, FR) |
| 3 | DG-BRE-3 | *Dactylis glomerata* L. | 'Brennus' (R2n, FR) |
| 4 | DG-BRE-4 | *Dactylis glomerata* L. | 'Brennus' (R2n, FR) |
| 5 | DG-BRE-5 | *Dactylis glomerata* L. | 'Brennus' (R2n, FR) |
| 6 | DG-BXL-1 | *Dactylis glomerata* L. | 'Barexcel' (Barenbrug, NL) |
| 7 | DG-BXL-2 | *Dactylis glomerata* L. | 'Barexcel' (Barenbrug, NL) |
| 8 | DG-BXL-3 | *Dactylis glomerata* L. | 'Barexcel' (Barenbrug, NL) |
| 9 | DG-BXL-4 | *Dactylis glomerata* L. | 'Barexcel' (Barenbrug, NL) |
| 10 | DG-BXL-5 | *Dactylis glomerata* L. | 'Barexcel' (Barenbrug, NL) |
| 11 | DG-BXL-6 | *Dactylis glomerata* L. | 'Barexcel' (Barenbrug, NL) |
| 12 | DG-RED-1 | *Dactylis glomerata* L. | 'Reda' (DSP/Agroscope , CH) |
| 13 | DG-RED-2 | *Dactylis glomerata* L. | 'Reda' (DSP/Agroscope , CH) |
| 14 | DG-RED-3 | *Dactylis glomerata* L. | 'Reda' (DSP/Agroscope , CH) |
| 15 | DG-RED-4 | *Dactylis glomerata* L. | 'Reda' (DSP/Agroscope , CH) |
| 16 | DG-RED-5 | *Dactylis glomerata* L. | 'Reda' (DSP/Agroscope , CH) |
| 17 | FP-COS-1 | *Festuca pratensis* Huds. | 'Cosmolit' (Saatszucht Steinach, DE) |
| 18 | FP-COS-2 | *Festuca pratensis* Huds. | 'Cosmolit' (Saatszucht Steinach, DE) |
| 19 | FP-COS-3 | *Festuca pratensis* Huds. | 'Cosmolit' (Saatszucht Steinach, DE) |
| 20 | FP-COS-4 | *Festuca pratensis* Huds. | 'Cosmolit' (Saatszucht Steinach, DE) |
| 21 | FP-COS-5 | *Festuca pratensis* Huds. | 'Cosmolit' (Saatszucht Steinach, DE) |
| 22 | FP-PAR-1 | *Festuca pratensis* Huds. | 'Paradisia' (DSP/Agroscope , CH) |
| 23 | FP-PAR-2 | *Festuca pratensis* Huds. | 'Paradisia' (DSP/Agroscope , CH) |
| 24 | FP-PAR-3 | *Festuca pratensis* Huds. | 'Paradisia' (DSP/Agroscope , CH) |
| 25 | FP-PAR-4 | *Festuca pratensis* Huds. | 'Paradisia' (DSP/Agroscope , CH) |
| 26 | FP-PAR-5 | *Festuca pratensis* Huds. | 'Paradisia' (DSP/Agroscope , CH) |
| 27 | FP-PAR-6 | *Festuca pratensis* Huds. | 'Paradisia' (DSP/Agroscope , CH) |
| 28 | FP-PRA-1 | *Festuca pratensis* Huds. | 'Pradel' (DSP/Agroscope , CH) |
| 29 | FP-PRA-2 | *Festuca pratensis* Huds. | 'Pradel' (DSP/Agroscope , CH) |
| 30 | FP-PRA-3 | *Festuca pratensis* Huds. | 'Pradel' (DSP/Agroscope , CH) |
| 31 | FP-PRA-4 | *Festuca pratensis* Huds. | 'Pradel' (DSP/Agroscope , CH) |
| 32 | FP-PRA-5 | *Festuca pratensis* Huds. | 'Pradel' (DSP/Agroscope , CH) |
| 33 | LP-ARA-1 | *Lolium perenne* L. | 'Arara' (DSP/Agroscope, CH) |
| 34 | LP-ARA-2 | *Lolium perenne* L. | 'Arara' (DSP/Agroscope, CH) |
| 35 | LP-ARA-3 | *Lolium perenne* L. | 'Arara' (DSP/Agroscope, CH) |
| 36 | LP-ARA-4 | *Lolium perenne* L. | 'Arara' (DSP/Agroscope, CH) |
| 37 | LP-ARA-5 | *Lolium perenne* L. | 'Arara' (DSP/Agroscope, CH) |
| 38 | LP-LAC-1 | *Lolium perenne* L. | 'Lacerta' (DSP/Agroscope, CH) |
| 39 | LP-LAC-2 | *Lolium perenne* L. | 'Lacerta' (DSP/Agroscope, CH) |
| 40 | LP-LAC-3 | *Lolium perenne* L. | 'Lacerta' (DSP/Agroscope, CH) |
| 41 | LP-LAC-4 | *Lolium perenne* L. | 'Lacerta' (DSP/Agroscope, CH) |
| 42 | LP-LAC-5 | *Lolium perenne* L. | 'Lacerta' (DSP/Agroscope, CH) |
| 43 | LP-LAC-6 | *Lolium perenne* L. | 'Lacerta' (DSP/Agroscope, CH) |
| 44 | LP-LIP-1 | *Lolium perenne* L. | 'Lipresso' (Euro Grass, DE) |
| 45 | LP-LIP-2 | *Lolium perenne* L. | 'Lipresso' (Euro Grass, DE) |
| 46 | LP-LIP-3 | *Lolium perenne* L. | 'Lipresso' (Euro Grass, DE) |
| 47 | LP-LIP-4 | *Lolium perenne* L. | 'Lipresso' (Euro Grass, DE) |
| 48 | LP-LIP-5 | *Lolium perenne* L. | 'Lipresso' (Euro Grass, DE) |
| 49 | TP-BNS-1 | *Trifolium pratense* L. | 'Bonus' (Selgen, CZ) |
| 50 | TP-BNS-2 | *Trifolium pratense* L. | 'Bonus' (Selgen, CZ) |
| 51 | TP-BNS-3 | *Trifolium pratense* L. | 'Bonus' (Selgen, CZ) |
| 52 | TP-BNS-4 | *Trifolium pratense* L. | 'Bonus' (Selgen, CZ) |
| 53 | TP-BNS-5 | *Trifolium pratense* L. | 'Bonus' (Selgen, CZ) |
| 54 | TP-DIP-1 | *Trifolium pratense* L. | 'Diplomat' (DSV, DE) |
| 55 | TP-DIP-2 | *Trifolium pratense* L. | 'Diplomat' (DSV, DE) |
| 56 | TP-DIP-3 | *Trifolium pratense* L. | 'Diplomat' (DSV, DE) |
| 57 | TP-DIP-4 | *Trifolium pratense* L. | 'Diplomat' (DSV, DE) |
| 58 | TP-DIP-5 | *Trifolium pratense* L. | 'Diplomat' (DSV, DE) |
| 59 | TP-DIP-6 | *Trifolium pratense* L. | 'Diplomat' (DSV, DE) |
| 60 | TP-PAV-1 | *Trifolium pratense* L. | 'Pavo' (DSP/Agroscope, CH) |
| 61 | TP-PAV-2 | *Trifolium pratense* L. | 'Pavo' (DSP/Agroscope, CH) |
| 62 | TP-PAV-3 | *Trifolium pratense* L. | 'Pavo' (DSP/Agroscope, CH) |
| 63 | TP-PAV-4 | *Trifolium pratense* L. | 'Pavo' (DSP/Agroscope, CH) |
| 64 | TP-PAV-5 | *Trifolium pratense* L. | 'Pavo' (DSP/Agroscope, CH) |
| 65 | TR-BEA-1 | *Trifolium repens* L. | 'Beaumont' (CW 090; Barenbrug, NL) |
| 66 | TR-BEA-2 | *Trifolium repens* L. | 'Beaumont' (CW 090; Barenbrug, NL) |
| 67 | TR-BEA-3 | *Trifolium repens* L. | 'Beaumont' (CW 090; Barenbrug, NL) |
| 68 | TR-BEA-4 | *Trifolium repens* L. | 'Beaumont' (CW 090; Barenbrug, NL) |
| 69 | TR-BEA-5 | *Trifolium repens* L. | 'Beaumont' (CW 090; Barenbrug, NL) |
| 70 | TR-BEA-6 | *Trifolium repens* L. | 'Beaumont' (CW 090; Barenbrug, NL) |
| 71 | TR-BOM-1 | *Trifolium repens* L. | 'Bombus' (DSP/Agroscope, CH) |
| 72 | TR-BOM-2 | *Trifolium repens* L. | 'Bombus' (DSP/Agroscope, CH) |
| 73 | TR-BOM-3 | *Trifolium repens* L. | 'Bombus' (DSP/Agroscope, CH) |
| 74 | TR-BOM-4 | *Trifolium repens* L. | 'Bombus' (DSP/Agroscope, CH) |
| 75 | TR-BOM-5 | *Trifolium repens* L. | 'Bombus' (DSP/Agroscope, CH) |
| 76 | TR-HEB-1 | *Trifolium repens* L. | 'Hebe' (Svalöf-Weibull, SE) |
| 77 | TR-HEB-2 | *Trifolium repens* L. | 'Hebe' (Svalöf-Weibull, SE) |
| 78 | TR-HEB-3 | *Trifolium repens* L. | 'Hebe' (Svalöf-Weibull, SE) |
| 79 | TR-HEB-4 | *Trifolium repens* L. | 'Hebe' (Svalöf-Weibull, SE) |
| 80 | TR-HEB-5 | *Trifolium repens* L. | 'Hebe' (Svalöf-Weibull, SE) |

**Table 5** Microsatellite primer sequences, melting temperatures and sequence repeat motifs

| Species | Locus | | Primer sequence 5'-3' | Tm (°C) | SSR (motif) repeats | Reference |
|---|---|---|---|---|---|---|
| *Dactylis glomerata* L. | DGSSR004 | F | GCTGTGGAGAAAAAAATGA | 52 | (TG)18(GA)8 | A01E02 |
| | | R | GATGCCATTAAGTTCAAAATG | | | (Xie et al., 2010) |
| | DGSSR006 | F | ATGCTGTTTGATCACAGTCA | 48 | (TG)23 | A01I13 |
| | | R | GTTGGACTGCCATTACTAGC | | | (Xie et al., 2010) |
| | DGSSR007 | F | TGGACTACATGATGAACCAGTACC | 52 | (GCC)15 | Dg_Contig1483 |
| | | R | GGTTCTCTTCCATGCTCATGTT | | | (Bushman et al., 2011) |
| | DGSSR008 | F | GTGCACATGTTTTTCAAGTAGTCC | 52 | (GTG)18 | Dg_Contig12060 |
| | | R | GCAATACCCTGAGATGGTTTACAT | | | (Bushman et al., 2011) |
| | DGSSR009 | F | CTCGTCGTCCCACAGAAACTAC | 56 | (GCTC)16 | Dg_Contig12302 |
| | | R | ATGACGGAGGAGCTGGAAAC | | | (Bushman et al., 2011) |
| | DGSSR010 | F | GCTGTTGCACAGTTAGTAGTTCGT | 60 | (GTTTG)20 | Dg_Contig1744 |
| | | R | CTAACACTGACAGCGTGTTCTTCT | | | (Bushman et al., 2011) |
| | DGSSR011 | F | AGGTAAACATTGGAGAGAAAGGCT | 60 | (GAA)18 | Dg_Contig66 |
| | | R | CTACTAGTTCCGTCATCTCCTGGT | | | (Bushman et al., 2011) |
| | DGSSR012 | F | CCAATAACACTGGACTCTCTTCCT | 60 | (TCC)12 | BG04063B2F01.r1 |
| | | R | TGGCATTTGAAGAGTTACCTATGA | | | (Bushman et al., 2011) |
| *Festuca pratensis* Huds. | FASSR44 | F | TCATTTGACGCCACTTGAAC | 51 | (GAG)8 | NFA064 |
| | | R | GTTTCTTGTCTTAGCGCCTTCCTTGGT | | | (Saha et al., 2004) |
| | FASSR49 | F | TCCTAAGCAGAGCTCGATCC | 51 | (GA)10 | NFA071 |
| | | R | GTTTCTTGAGGTTGGCGAACTTCCTC | | | (Saha et al., 2004) |
| | LMSSR15 | F | TCATTGCCCTGACTACCAGG | 56 | (GT)5 | LM15 |
| | | R | GCATCGCCTTTCTCAGAGTG | | | (Studer, Widmer, Enkerli, & Kölliker, 2006) |
| | LMSSR26 | F | AAGAAAATGCAATGTCTAAACAGATTAGTT | 54 | (GT)15(GCGT)5(GT)11 | LM26 |
| | | R | GTTTCTTCGCCTCATGAACACTTTATATATTCTAGAA | | | (Studer et al., 2006) |
| | LMSSR29 | F | AGAGAGAGTTTGCTCACAC | 46 | (GT)5 | LM29 |
| | | R | TTCAAGTGCAAGCAAAACA | | | (Studer et al., 2006) |
| | LPSSR01 | F | CACCTCCCGCTGCATGGCATGT | 55 | (GT)8(AGGT) | M4213 |
| | | R | TACAACGACATGTCAAGG | | | (Kubik, Sawkins, Meyer, & Gaut, 2001) |
| | LPSSR27 | F | GTATAGTACCCATTCCGT | 54 | (CA)22 | PR3 |
| | | R | GCCGCCCTGCCATGCTG | | | (Kubik et al., 2001) |
| | LPSSR47 | F | AGCCACACTTTACCTAATGCTG | 55 | (AC)17 | Uni001 |
| | | R | CCCGCAAAACTTACAATTAAA | | | (Jensen et al., 2005) |
| *Lolium perenne* L. | LPSSR07 | F | AAAGACCGCATACGAAGT | 50 | (CA)27 | LPSSRH01A02 |
| | | R | AACCAAAGCCTCAAGACA | | | (Jones, Dupal, Kölliker, Drayton, & Forster, 2001) |
| | LPSSR09 | F | GAGGCACCGGCCATGGAG | 58 | (CTT)20 | LPSSRH01A10 |
| | | R | AGGACGAGCCACTCACTTG | | | (Jones et al., 2001) |
| | LPSSR14 | F | TGGAATAACGATGAAAAG | 46 | (CA)4(TACA)4 | LPSSRH02C11 |
| | | R | CATCACGAATTAACAAGAG | | | (Jones et al., 2001) |
| | LPSSR17 | F | TGACTTCTCTCGATCCT | 55 | (CT)17 | LP8a |
| | | R | ATGTGACTACAAAACCA | | | (Jensen et al., 2005) |
| | LPSSR20 | F | GCGTAAGAGAGAGGGCGAT | 55 | (GA)10GG(GA)GG(GA) | LP194 |
| | | R | ACGTATGTCCAACAGGT | | | (Kubik et al., 2001) |
| | LPSSR23 | F | TAGCTTTCTATGCAAAGCT | 52 | (GT)8 | M844 (Kubik et al., 2001) |
| | | R | CACTTCACTTTTCTTGCA | | | |
| | LPSSR13 | F | ATTGACTGGCTTCCGTGTT | 56 | (CA)9 | LPSSRH01H06 (Jones et al., 2001) |
| | | R | CGCGATTGCAGATTCTTG | | | |
| | LPSSR16 | F | CGGCCACCCTTGATAGAG | 59 | (CA)21 | LPSSRK01A11 (Jones et al., 2001) |
| | | R | TCGTCAAGGATCCGGAGA | | | |
| *Trifolium pratense* L. | TPSSR010 | F | TGGACATCATGGTTCCCACG | 65 | (CT)28 | TPSSR10 |
| | | R | TCAAAGAAGCAAGGAACGGTG | | | (Roland Kölliker, Enkerli, & Widmer, 2006) |
| | TPSSR017 | F | AAGCAGCGAGACTTCCCTTTG | 60 | (CTT)22 | TPSSR17 |
| | | R | TGGAAGGTTAACATCGAGAGCA | | | (Roland Kölliker et al., 2006) |
| | TPSSR034 | F | GTTAGTGCGCGAAAGGAAGG | 60 | (CT)7; (CT)22; (CT)8 | TPSSR34 |
| | | R | TTGTTCAGTGGATCAGTGAGTAACACAA | | | (Roland Kölliker et al., 2006) |
| | TPSSR044 | F | TCTGGCTTTCTTGCCGATATC | 60 | (ATTG)7; (GA)20; (GA)6 | TPSSR44 |
| | | R | TTACACCCTGTTTCGTGAAGCAC | | | (Roland Kölliker et al., 2006) |
| | TPSSR045 | F | TGTGTTATGGTGAAGTTCAAAATATAATTTC | 60 | (GA)27; (GA)5 | TPSSR45 |
| | | R | TTCCAATGGCGTCAATGGTCTC | | | (Roland Kölliker et al., 2006) |
| | TPSSR046 | F | TCAAATAAAACTTTCATAACGTTCATCTC | 60 | (TC)28 | TPSSR46 |
| | | R | TTTCCGAAGAAACCATTATCTACGTTG | | | (Roland Kölliker et al., 2006) |
| | TPSSR050 | F | AAGGCCCATGTTGAAACTGC | 60 | (CT)23 | TPSSR50 |
| | | R | TTTTTGTTCAGGAAAATGAGGCG | | | (Roland Kölliker et al., 2006) |
| | TPSSR052 | F | ATTCCTTCCATCTTCTCTATGT | 60 | (CT)31 | TPSSR52 |
| | | R | TTTTATATTAATGGGAGTTAGTATGATCTA | | | (Roland Kölliker et al., 2006) |
| *Trifolium repens* L. | TRSSR04 | F | AGAAAGGTGAATGATGAAA | 50.9 | (GAA)20 | TPSSRA01H11 |
| | | R | TCTAATTCTTCCAATAGGG | | | (R. Kölliker, Jones, Drayton, Dupal, & Forster, 2001) |
| | TRSSR05 | F | TTTTGCTAATAAGTAATGCTGC | 57 | (TG)11 | TRSSRA02B08 |
| | | R | GGACATTATGCAATGGTGAG | | | (R. Kölliker et al., 2001) |
| | TRSSR09 | F | AAGTGTTGGACAAGGAAACTAGG | 62 | (TA)7(CA)19 | TRSSRDXX16 |
| | | R | TCTCTAGATCACCGGCATTG | | | (R. Kölliker et al., 2001) |
| | TRSSR10 | F | GAGGTTAGTCAATTAGGACTTCTCT | 62 | (ACACC)10 | NF207316 |
| | | R | TCATTAGGGGACCATTGCTTT | | | (Zhang, He, Zhao, Bouton, & Monteros, 2008) |
| | TRSSR11 | F | AAAATCTCTACCATGACTTTGCTTTA | 60 | (TGTGT)5 | NF207415 |

| | | | | | |
|---|---|---|---|---|---|
| | R | CCAGAACAATAGAAAGATCATCA | | | (Zhang et al., 2008) |
| TRSSR12 | F | TTGGTATGGTTTCATGTCCCT | 62 | (TATTT)4 | NF207986 |
| | R | GCTCTGGTAAATAGATTGGATCGT | | | (Zhang et al., 2008) |
| TRSSR13 | F | ACAGCTCCCCCTTAACACCT | 60 | (ATCAC)4 | NF208653 |
| | R | ATGATCTGCACTGCTCTGGAT | | | (Zhang et al., 2008) |
| TRSSR14 | F | CAGACCACAAAACAATCAATG | 57 | (AC)9 | NF206403 |
| | R | CCAAACGATGTTGTGTGCTAA | | | (Zhang et al., 2008) |

PCR reactions were conducted in a C1000 Touch™ Thermal Cycler (Bio-Rad Laboratories, Inc., Hercules CA, USA). To increase specificity of the PCR reactions, touchdown PCR with the following conditions was employed: 5 min at 94°C for initial denaturation, then 11 cycles of 30s at 94 °C (denaturation), 1 min initially 12 °C above annealing temperature, decreasing 1 °C every cycle, and 1 min at 72 °C (elongation) followed by 29 cycles of 30s at 94 °C (denaturation), 1 min at the primer specific annealing temperature and 1 min at 72 °C (elongation). Final extension was performed at 72 °C for 15 min.

Analysis of SSR fragments was performed on an ABI PRISM® 3130xl Genetic Analyzer (Applied Biosystems®, Waltham MA, USA). One µl of a 1:20 dilution of each PCR product was added to 18 µl Hi-Di™Formamide and 1 µl GeneScan™ 400 HD [ROX™ DYE] size standard (Applied Biosystems®, Waltham MA, USA), then incubated for 3 min at 94 °C in a PTC-200 Peltier Thermal Cycler (Bio-Rad Laboratories, Inc., Hercules CA, USA) and then immediately chilled on ice for a few minutes. Genotyping of SSRs was done using microsatellite analysis tools of the Thermo Fisher Connect online application (Thermo Fisher Scientific Inc., Waltham MA, USA).

**Table 6** Summary of diversity statistics of each SSR marker.

| **Species name** | **SSR†** | **Number of observed alleles** | **Simpson index** | **Nei's 1978 gene diversity** | **Evenness** |
|---|---|---|---|---|---|
| *Dactylis glomerata* L. | **DGSSR004** | 17 | 0.919271 | 0.93883 | 0.866505 |
| | **DGSSR006** | 14 | 0.869638 | 0.887755 | 0.749432 |
| | DGSSR007 | 6 | 0.59808 | 0.621083 | 0.612239 |
| | DGSSR008 | 2 | 0.5 | 0.516129 | 1 |
| | DGSSR009 | 4 | 0.606421 | 0.628079 | 0.779961 |
| | **DGSSR010** | 10 | 0.815193 | 0.835075 | 0.738955 |
| | **DGSSR011** | 9 | 0.81213 | 0.828054 | 0.795003 |
| | **DGSSR012** | 15 | 0.8632 | 0.880816 | 0.723261 |
| | mean | 9.63 | 0.747991 | 0.766978 | 0.78317 |
| *Festuca pratensis* Huds. | FASSR44 | 2 | 0.142012 | 0.153846 | 0.531316 |
| | **FASSR49** | 4 | 0.577778 | 0.619048 | 0.706266 |
| | LMSSR15 | 3 | 0.580499 | 0.609524 | 0.886511 |
| | **LMSSR26** | 5 | 0.698225 | 0.75641 | 0.780019 |
| | **LMSSR29** | 4 | 0.576389 | 0.601449 | 0.8175 |
| | LPSSR01 | 3 | 0.538194 | 0.561594 | 0.8907 |
| | **LPSSR27** | 7 | 0.801512 | 0.837945 | 0.850622 |
| | **LPSSR47** | 11 | 0.853186 | 0.900585 | 0.760914 |
| | mean | 4.88 | 0.595974 | 0.63005 | 0.777981 |
| *Lolium perenne* L. | **LPSSR07** | 17 | 0.920118 | 0.956923 | 0.843652 |
| | LPSSR09 | 2 | 0.142012 | 0.153846 | 0.531316 |
| | LPSSR14 | 5 | 0.7392 | 0.77 | 0.849816 |
| | LPSSR17 | 3 | 0.650284 | 0.679842 | 0.963862 |
| | **LPSSR20** | 12 | 0.862004 | 0.901186 | 0.758189 |
| | **LPSSR23** | 6 | 0.7648 | 0.796667 | 0.847979 |
| | **LPSSR13** | 13 | 0.872428 | 0.905983 | 0.753738 |
| | **LPSSR16** | 7 | 0.81405 | 0.852814 | 0.884596 |
| | mean | 8.13 | 0.720612 | 0.752158 | 0.804143 |
| *Trifolium pratense* L. | TPSSR17 | 4 | 0.485207 | 0.525641 | 0.607021 |
| | **TPSSR45** | 19 | 0.940972 | 0.981884 | 0.938838 |
| | **TPSSR46** | 14 | 0.903592 | 0.944664 | 0.849868 |
| | **TPSSR50** | 16 | 0.920139 | 0.960145 | 0.868223 |
| | TPSSR44 | 10 | 0.833333 | 0.869565 | 0.77264 |
| | **TPSSR34** | 13 | 0.903047 | 0.953216 | 0.878151 |
| | **TPSSR10** | 11 | 0.87 | 0.915789 | 0.814824 |
| | TPSSR52 | 8 | 0.65625 | 0.7 | 0.523183 |
| | mean | 11.88 | 0.814068 | 0.856363 | 0.781593 |
| *Trifolium repens* L. | **TRSSR04** | 22 | 0.917695 | 0.93501 | 0.75259 |
| | TRSSR05 | 7 | 0.824263 | 0.844367 | 0.898048 |
| | **TRSSR09** | 16 | 0.915 | 0.938462 | 0.869889 |
| | **TRSSR10** | 11 | 0.882842 | 0.897081 | 0.901989 |
| | TRSSR11 | 6 | 0.777284 | 0.794949 | 0.88005 |
| | TRSSR12 | 9 | 0.7976 | 0.813878 | 0.79665 |
| | **TRSSR13** | 13 | 0.868229 | 0.882232 | 0.79331 |
| | TRSSR14 | 10 | 0.871304 | 0.889456 | 0.876394 |
| | mean | 11.75 | 0.856777 | 0.874429 | 0.846115 |

† Highlighted SSRs were used for pairwise distance calculations (Fig. 4 in main text). This table was produced using poppr. An explanation of the columns "number of observed alleles", "Simpson index", "Nei's 1978 gene diversity" and "eveness" is found in https://grunwaldlab.github.io/Population_Genetics_in_R/Genotypic_EvenRichDiv.html

**Table 7** Genome size of the species analyzed in the discovery phase

| Family | Species | 1C genome size (mean, Mbp) † | Reference |
|---|---|---|---|
| Fabaceae | *Lotus corniculatus* L. | 846 | (João Loureiro, Castro, Cerca de Oliveira, Mota, & Torices, 2013; Pustahija et al., 2013) |
| | *Medicago sativa* L. | 1,264 | (Blondon, Marie, Brown, & Kondorosi, 1994; Pustahija et al., 2013) |
| | *Onobrychis viciifolia* Scop. | 1,225 | (Leitch, Johnston, Pellicer, Hidalgo, & Bennett, 2019) |
| | *Trifolium pratense* L. | 418 | (Vižintin, Javornik, & Bohanec, 2006) |
| | *Trifolium repens* L. | 1,039 | (Pustahija et al., 2013) |
| Poaceae | *Arrenatherum elatius* L. | 7,840 | (Grime & Mowforth, 1982) |
| | *Alopecurus pratensis* L. | 6,664 | (Olszewska & Osiecka, 1982) |
| | *Cynosurus cristatus* L. | 2,987 | (Smarda, Bures, Horova, Foggi, & Rossi, 2008) |
| | *Dactilys glomerata* L. | 3,953 | (Creber, Davies, Francis, & Walker, 1994) |
| | *Festuca pratensis* Huds. | 4,778 | (Kopecký et al., 2010) |
| | *Festuca rubra* L. | 7,678 | (J. Loureiro, Kopecký, Castro, Santos, & Silveira, 2007; Smarda et al., 2008) |
| | *Lolium multiflorum* Lam. | 3,979 | (Kopecký et al., 2010) |
| | *Lolium perenne* L. | 4,055 | (Kopecký et al., 2010) |
| | *Phleum pratense* L. | 4,018 | (Leitch et al., 2019) |
| | *Poa pratensis* L. | 4,724 | (Arumuganathan, Tallury, Fraser, Bruneau, & Qu, 1999; Bennett, Smith, & Smith, 1982) |
| | *Trisetum flavescens* L. | 2,548 | (Leitch et al., 2019) |
| Fabaceae | Average | 958.4 | |
| Poaceae | Average | 4,838.5 | |

† Genome sizes according to the Plant DNA C-Value Database (https://cvalues.science.kew.org; Leitch et al., 2019)

**Figure 3** Distribution of indels in sequence capture and amplicon sequencing data

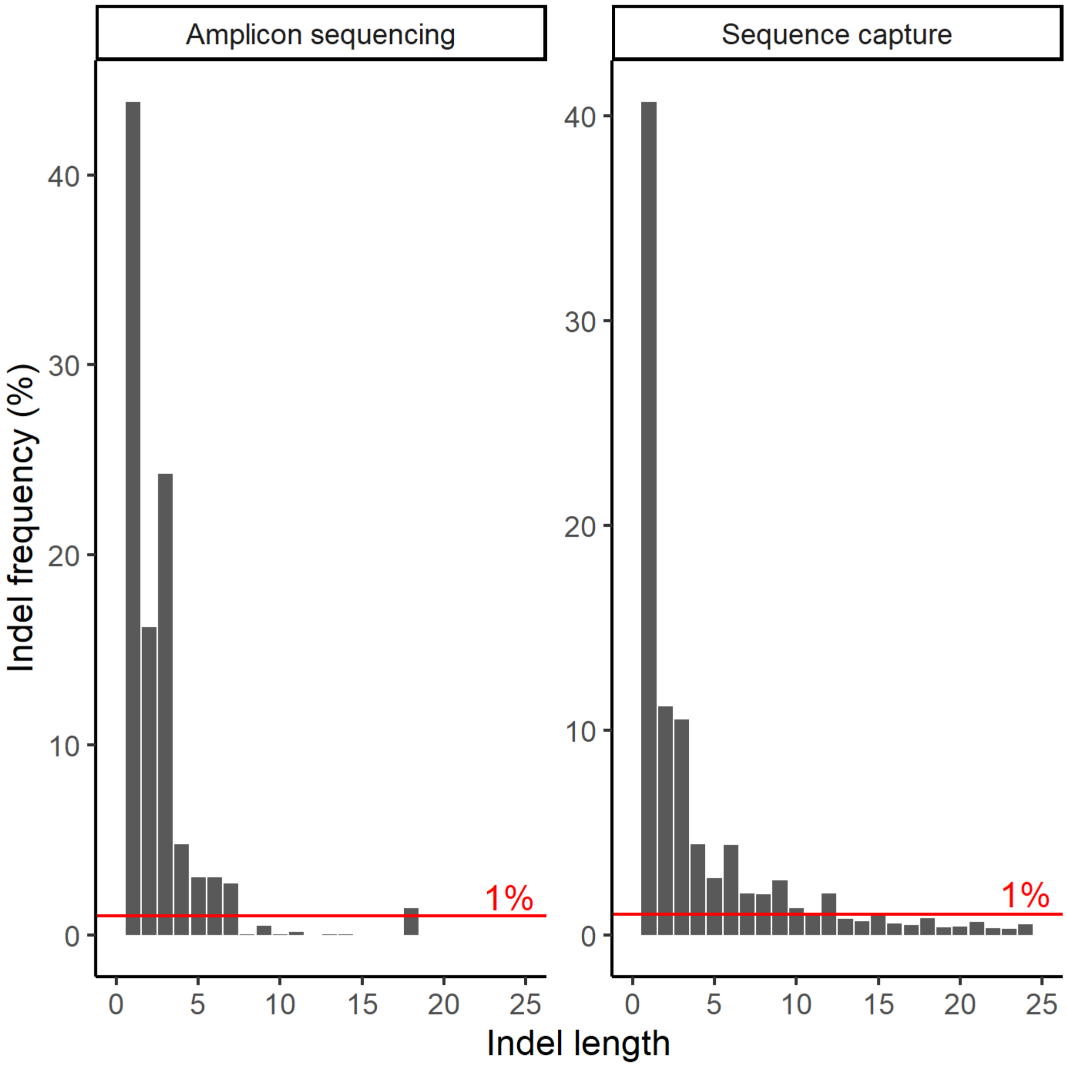
